# In their sister’s footsteps: Taxonomic divergence obscures substantial functional overlap among the metabolically diverse symbiotic gut communities of adult and larval turtle ants

**DOI:** 10.1101/2021.09.08.459499

**Authors:** Benoît Béchade, Yi Hu, Jon G. Sanders, Christian S. Cabuslay, Piotr Łukasik, Bethany R. Williams, Valerie J. Fiers, Richard Lu, John T. Wertz, Jacob A. Russell

## Abstract

Gut bacterial symbionts can support animal nutrition by facilitating digestion and providing valuable metabolites. While the composition of gut symbiont communities shifts with host development in holometabolous insects, changes in symbiotic roles between immature and adult stages are not well documented, especially in ants. Here, we explored the metabolic capabilities of microbiomes sampled from herbivorous turtle ant (*Cephalotes* sp.) larvae and adult workers through genomic and metagenomic screenings and targeted *in vitro* metabolic assays. We reveal that larval guts harbor bacterial symbionts from the Enterobacteriales, Lactobacillales and Rhizobiales orders, with impressive metabolic capabilities, including catabolism of plant and fungal recalcitrant fibers common in turtle ant diets, and energy-generating fermentation. Additionally, several members of the specialized turtle ant adult gut microbiome, sampled downstream of an anatomical barrier that dams large food particles, show a conserved potential to depolymerize many dietary fibers and other carbohydrates. Symbionts from both life stages have the genomic capacity to recycle nitrogen, synthesize amino acids and B-vitamins, and perform several key aspects of sulfur metabolism. We also document, for the first time in ants, an adult-associated Campylobacterales symbiont with an apparent capacity to anaerobically oxidize sulfide, reduce nitrate, and fix carbon dioxide. With help of their gut symbionts, including several bacteria likely acquired from the environment, turtle ant larvae appear as an important component of turtle ant colony digestion and nutrition. In addition, the conserved nature of the digestive, energy-generating, and nutritive capacities among adult-enriched symbionts suggests that nutritional ecology of turtle ant colonies has long been shaped by specialized, behaviorally-transferred gut bacteria with over 46 million years of residency.

## Introduction

Microbial symbioses widely influence the survival and evolution of eukaryotes, and microbe-driven nutritional mutualisms comprise a well-characterized set of adaptive strategies [1–5]. In animals, these microbially contributed benefits include nitrogen (N)-fixation and N-recycling [6–9], in addition to the synthesis of dietarily rare, essential metabolites, such as amino acids [10] or B-vitamins [11–13]. In addition to nutrient concentration, conservation, and provisioning, microbial symbionts can contribute to beneficial digestive functions, helping to catabolize recalcitrant carbohydrates and polyphenols that are indigestible for many animals, yielding forms of carbon which can be readily utilized by these hosts [14–17]. Thanks to this array of potential benefits, microbial symbionts have expanded the range of natural food sources utilized by animal hosts, granting access to previously inhospitable dietary niches [18–21].

In several animal species the microbial symbiont community changes with the age of the host. This is especially the case in animals with dramatic shifts in the diet throughout their development, such as mammary gland-feeding mammals [22, 23]. Holometabolous insects undergo a complete reorganization of their body plan during metamorphosis, producing adults with a transformed physiology and ecology, and, often, with distinct gut microbiomes bearing little resemblance to those in larvae [24]. Such stage-specific symbioses are common in some eusocial hymenopterans, like honeybees and ants [25–28], despite the possibility of social transfer of microbes between life stages. The digestive roles of gut microbial symbionts in early life stages have been studied in few insects from other orders, where bacteria occasionally show capacities for *in vitro* degradation of recalcitrant, plant-derived fibers in some lepidopterans, orthopterans and dipterans [29–33], and for N-fixation and N-recycling in some coleopteran larvae [34, 35]. However, eusocial hymenopteran larvae have rarely been studied in functional analyses of gut microbial symbionts [but see 36,37].

In ants, larvae are assumed to play the role of an external stomach for older colony members, since adult ants possess anatomical restrictions constraining their digestion of solid food, such as a narrow esophagus and a small opening of <0.2 micrometers through the proventriculus, an organ that separates the crop from the midgut [38]. It has been shown that ant larvae can liquefy solid food and regurgitate it to adult workers or queens in predatory and omnivorous species [39–41]. This larval digestive role was also reported in fungus-growing ant larvae, which possess a simplified gut and have a higher enzymatic activity against complex carbohydrates than adult workers [42]. Digestive roles of symbionts in some non-hymenopteran insects therefore raise the question of whether gut bacteria support this ‘digestive caste’ function of ant larvae.

Turtle ants of the genus *Cephalotes* are a group of eusocial, neotropical arboreal insects. While their diets sometimes contain simple sources of carbohydrates, like extrafloral nectar, plant wound secretions, or insect honeydew [43–45], they are also enriched for recalcitrant forms of N and carbohydrates. Common turtle ant food sources containing such recalcitrant nutrients include wind-dispersed pollen, mammalian urine, bird droppings, and potentially other cryptic materials such as lichen and fungi [44–50]. Among other pollen feeders, honeybees harbor pectin-degrading gut symbionts, and symbionts that synthesize organic acids (including short chain fatty acids and lactic acid), which are thought to be a key energy source for these eusocial insects [15,51,52]. Insects feeding on urine and bird droppings, including olive fruit flies, are thought to benefit from N-recycling conducted by gut symbionts [53, 54]. Regarding ants with low trophic levels [55], impactful contributions of symbiont N-recycling to amino acid pools were suggested in *Dolichoderus* ants through genomic inference [56], but have been directly demonstrated for *Camponotus* ants [57] and, more recently, for turtle ants [58].

Matching the expectations from their limited diet, turtle ant adult workers harbor populous bacterial communities in their midgut and hindgut lumen [59, 60]. These communities are conserved across the turtle ant phylogeny [61], and are comprised of strains from over a dozen seemingly specialized lineages, spanning eight bacterial orders (Burkholderiales, Campylobacterales, Flavobacteriales, Opitutales, Pseudomonadales, Rhizobiales, Sphingobacteriales and Xanthomonadales) [58, 62]. A recent investigation revealed symbiotic N-recycling by a subset of this gut bacterial community, and worker ant acquisition of symbiont-synthesized amino acids, made from dietarily acquired, and symbiont-recycled N-wastes [58]. This ancient nutritional mutualism has likely been key to the turtle ants’ prolonged exploitation of marginal diets, bearing resemblance to microbial mutualisms of termites [63, 64].

What is not known for this symbiosis in adult turtle ants is whether conserved symbionts impact the use of the cell wall biomass from plant and fungal tissue encountered in the turtle ant diet, aiding in the breakdown of recalcitrant plant cell wall materials – here referred to as ‘fibers’ – including lignin, pectin, xylan/hemicellulose and cellulose. Due to the aforementioned anatomical restrictions in adult ants, any processing of solid, macromolecular food matter would need to occur outside of adult midguts and hindguts (ileum and rectum) where the bulk of conserved bacteria live [26], raising questions of whether turtle ant larval guts might be active sites for such digestion. Turtle ant larvae share some bacteria with adult workers [65], but evidence from other studies suggests the overall bacterial composition of larval guts differs from that in the guts of adults [26, 66]. Key digestive and nutritional functions in turtle ant larval gut microbiome have never been explored thus far and the degree of metabolic complementarity/overlap between adult and larval gut microbiomes had yet to be measured.

Given the capacity for trophallactic food exchange between developmental stages in ants, the expectation that ant larvae are the digestive caste and the possibility that turtle ant larvae harbor a taxonomically unique bacterial community, we asked: (1) do microbiomes vary in their overall metabolic potential between larval and adult turtle ants, and (2) are predicted functional differences consistent with the partitioning of host-beneficial N metabolisms and fiber catabolism into complementary, stage-specific nutritional symbioses? We also asked: (3) do symbionts from adult and larval turtle ant gut microbiomes encode other nutritive functions, shown to be fulfilled by microbial symbionts from other invertebrates, including the synthesis of B-vitamins [*e.g.* 66–69], the metabolism of sulfur [*e.g.* 4,70], or the production of energetic organic acids [*e.g.* 71,72]? We addressed these questions by exploring metabolic potentials in turtle ant larvae and adult workers at the whole gut microbiome and symbiont levels, using shotgun metagenomics, bacterial isolate genomics, and *in vitro* metabolic assays (**S1 Fig**). Through our investigation, we gained unique insights into the potential nutritional benefits that the remarkably conserved extracellular gut symbionts of turtle ants provide to their host.

## Materials and methods

Detailed protocols for cultivation procedures, (meta)genomic bioinformatic analyses and statistical tests can be found in **S1 Text**.

### Genomic and metagenomic sequencing

In addition to 18 previously published metagenomes from adult workers across 17 *Cephalotes* species (including *C. varians* colonies PL005 and PL010), and to previously published genome sequences from 14 cultured, worker-derived symbionts [58], we generated two new metagenomes and two new bacterial cultured isolate genomes (IGs) from turtle ant larvae.

For the larval gut metagenomes, larvae from two *C. varians* colonies, PL005 and PL010 (collected in Key Largo, FL, USA, in 2012: 25°7’24.59’N, 80°24’11.88’W and 25°11’36.66’N, 80°21’25.26’W, respectively), were washed in 70% ethanol and rinsed in sterile water before dissection. Larval guts were dissected under a light microscope using bleach-sterilized fine forceps. Dissected guts from ten larvae were pooled, separately for each colony, then flash-frozen in liquid nitrogen, before grinding with a sterile pestle in a 1.5 ml tube. DNA was subsequently extracted using the Qiagen DNeasy Blood and Tissue kit (Qiagen Ltd., Hilden, Germany), following the manufacturer’s protocol for Gram-positive bacteria. For metagenomic library preparation, fragmented genomic DNA from the resulting samples was size-selected, ligated with adapters and then further amplified before sequencing (2×100bp or 2×150bp) on an Illumina HiSeq2500 machine (Biopolymers Facility of Harvard University, Boston, MA, USA). Low quality reads and adapters were removed using Trimmomatic [74] after de-multiplexing. Assembly of reads from individual libraries was performed by using IDBA-UD software [75] with *k*-values of 20, 40, 60, 80, and 100. Assembly statistics were then calculated with QUAST [76]. Additional information on the metagenomes used can be found in **S1 Table**.

For genome sequencing of cultured isolates from larvae, specimens of *C. texanus* were collected from Texas in 2015 and 2016 (colony JR029a from Floresville, TX, USA – 29°07’47.31’N, 98°09’02.22’W; colony JDR108 from Gonzales, TX, USA – 29°27’58.82’N, 97°34’2.32’W). After larval gut dissection and bacterial cultivation and isolation (see **S1 Text**), two cultured isolates from *C. texanus* larvae were selected for genome sequencing: *Enterococcus* sp. JR029-101 and *Staphylococcus* sp. JDR108L-110-1. DNA was extracting following the same protocol as detailed above. DNA preparations were sent for sequencing on a PacBio RS II system (Pacific Biosciences, Menlo Park, CA, USA) at the Genomics Core Facility of the College of Medicine at Drexel University (Philadelphia, PA, USA). The PacBio reads were assembled using HGAP 2.3 [77]. Circlator 1.0.2 [78] was used to attempt circularizing bacterial genomes, which also involves permutation of the circle to place a particular gene (*dnaA*) at the beginning of all assemblies. Errors were corrected, and methylation motifs were computed, using RS Modification and Motif Analysis within the SMRT Analysis System (v2.3, January 2015; PacBio, Menlo Park, CA, USA). The assembled scaffold data were run through QUAST [76] to estimate assembly statistics. Additional information on the IGs used can be found in **S2 Table**.

Like the previously published datasets, the scaffolds of the newly assembled *C. varians* PL005 and PL010 larval metagenomes, and those from the two aforementioned IGs from *C. texanus* larvae – *Enterococcus* sp. JR029-101 and *Staphylococcus* sp. JDR108L-110-1 – were uploaded to the IMG/M-ER database for annotation and, in the former case, taxonomic classification [79].

### Taxonomic assignment and metagenome-assembled genome binning

For the assembled *C. varians* larval and adult worker gut metagenomes, taxonomic assignment was performed in IMG/M-ER which uses USEARCH [80] to screen genes against the IMG/M-ER identified genome database [81].

To confirm that we sampled typical gut bacterial communities in our turtle ant larval gut metagenomes, we examined results of Hu et al. [66], which characterized dominant taxa across larvae of varying size for 12 *Cephalotes* species through 16S rRNA amplicon sequencing (**S2 Text**). Our shotgun-based data suggested a high abundance of Rhizobiales for both PL005 (**S2 Fig**) and PL010 (Fig 1A) colony larvae – matching overall trends of the amplicon-based study. But the colony PL005 differed in exhibiting a high abundance of common adult-derived symbionts and low abundance of Enterobacteriales and Lactobacillales. The amplicon-based study revealed that Lactobacillales and Enterobacteriales were abundant in larvae (**S3 Fig**). Larval gut communities enriched with these two bacterial orders, like our PL010 larval gut metagenome (Fig 1A), are typical of older larval gut microbiomes [66]. For these reasons, we placed most of our focus on the PL010 larval metagenome to infer functions most likely representative of late larval stages, when larvae possess large masses of solid food in their gut.

**Fig 1.**
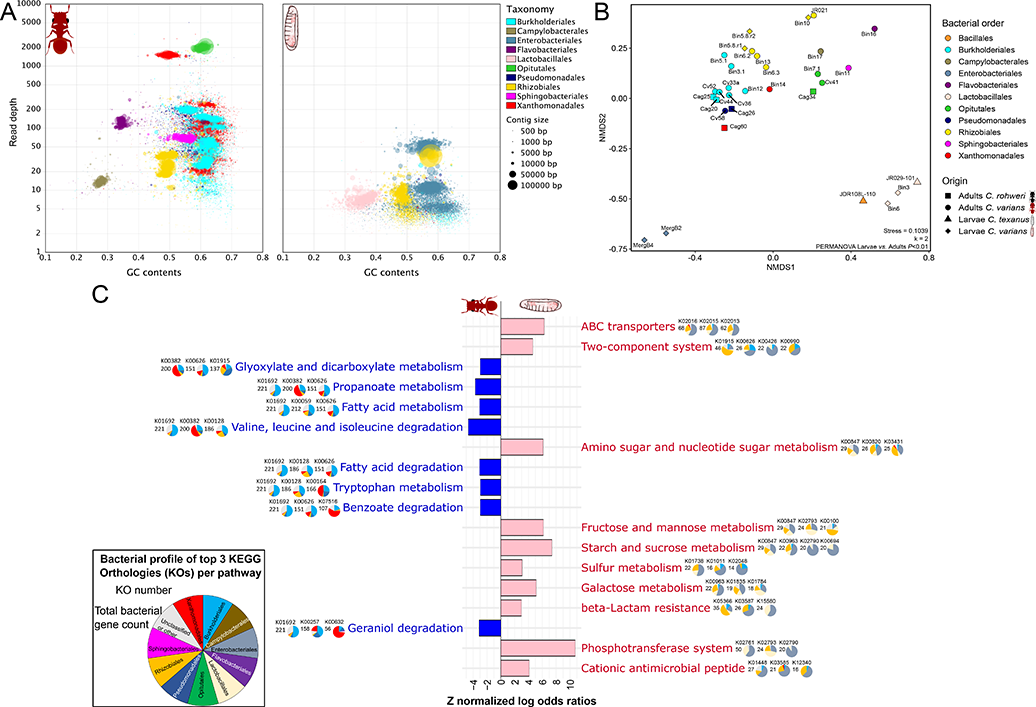
Taxonomic and functional overview of the turtle ant gut microbiomes. **(A)** Taxon-annotated GC-coverage blobplots illustrating the bacterial community composition of the *Cephalotes varians* PL010 adult worker (left) and larval (right) gut metagenomes. Assembled scaffolds from both metagenomes were taxonomically annotated using the USEARCH-IMG/M-ER pipeline [79–81]. Guanine (G) and cytosine (C) content, which varies among core bacterial genomes, is displayed on the x-axis. Depth of sequencing coverage is a proxy for the relative abundance of core symbionts, shown on the y-axis. The *C. varians* PL010 adult worker is similar as in [58]. **(B)** Non-Metric Multi-Dimensional Scaling (NMDS) plot based on a Bray-Curtis distance matrix calculated from normalized KO-encoding gene counts for each larval and adult symbiont genome. The Permutation Multivariate Analysis of Variance (PERMANOVA) was computed between larval and adult symbiont genomes with 9,999 permutations. **(C)** KEGG pathways with significantly different normalized gene counts in the *C. varians* PL010 larval gut metagenome compared to the adult worker metagenome. KEGG pathways are sorted from the most (top) to the least (bottom) enriched in both larval and adult metagenomes. Gene counts per KEGG pathway were normalized by the total count of bacterial KO-encoding genes. Corresponding *Z*-normalized log odds ratios between adult and larval metagenomes were calculated. Significantly enriched KEGG pathways are indicated by colored bars. Pink bars correspond to pathways significantly enriched in the larval metagenome compared to the adult one, while blue bars are for pathways significantly enriched in the adult metagenome compared to the larval one. Significance was evaluated by applying the False Discovery Rate correction of Benjamini-Hochberg [92] with a critical value of 0.05. For each of these KEGG pathways, the top three most encoded KOs are shown along with pie charts depicting which bacterial symbionts code for these KOs. The full list of gene counts per KEGG pathway can be found in **S6 Table** and the list of KOs in significantly enriched pathways can be found in **S7 Table**.

To enable functional characterization of individual symbiont strains, a subset of the assembled scaffolds from the *C. varians* PL010 larval gut metagenome were binned into metagenome-assembled genomes (MAGs) using CONCOCT [82] within the Anvi’o metagenome visualization and annotation pipeline v2.2.2 [83], following the workflow described online (http://merenlab.org/2016/06/22/anvio-tutorial-v2/). Metagenomic bins were classified as MAGs if they had more than 60% completeness and less than 10% redundancy (**S3 Table**). In total, 7 MAGs were produced from the PL010 larval metagenome. This added to 11 MAGs produced, in a prior study, from the PL010 worker gut metagenome [58].

### Genome comparisons

To assess the levels of similarity among 32 *Cephalotes* gut symbiont genomes (*i.e.* 2 new larval IGs from *C. texanus* larvae, 7 MAGs from *C. varians* larvae, 7 IGs from *C. varians* workers, 5 IGs from *C. rohweri* workers, and 11 MAGs from *C. varians* workers), pairwise orthoANI (*i.e.* Average Nucleotide Identity) values [84] were calculated using the OrthoANIu algorithm [85]. Three *Ventosimonas* (Peudomonadales) IGs (Cag26, Cag27 and Cag32) from *C. rohweri* were found to be highly similar to one another (orthoANI>98.5%). To minimize pseudoreplication, only one of these three IGs (Cag26) was included in our remaining analyses. For illustration purposes, a maximum likelihood phylogeny was constructed in RAxML Black Box [86] on the CIPRES Science Gateway [87] to show relatedness among these bacteria using seven concatenated bacterial marker genes (*rplA*, *rplB*, *rplC*, *rpsB*, *rps*C, *rpsE* and *tsf*) with 999 bootstrap iterations.

### *In vitro* metabolic assays

To test whether bacterial symbionts isolated from larval and adult worker guts of turtle ants can complete specific nutritional functions, we performed *in vitro* metabolic assays, focusing on functions including pectin degradation (representative of the broader potential for fiber degradation), carbohydrate and organic acid catabolism, and urea hydrolysis (representative of N-recycling). First, we followed a previously described pectin plate assay method [15] to assess pectin depolymerization by cultured bacteria. Test media were prepared by combining poly-d-galacturonic acid methyl-ester from apple, with agar in Tris-HCl pH 8.6. The solution was sterilized and pipetted into the wells of a 24-well plate. After symbiont inoculation into the agar wells and two days of growth, each well was flooded with cetyltrimethylammonium bromide. Development of a clear zone around the bacterial colony was considered evidence of pectinase activity.

In separate assays for carbohydrate and organic acid catabolism, testing either depolymerization (as above for pectin) or oxidation activity, we utilized GN3 microplates (Biolog, Hayward, CA, USA) according to the manufacturer’s instructions. Cultured bacterial isolates were inoculated by taking a small sample from a Tryptic Soy Agar plate and resuspending it in inoculating fluid A until a transmittance of 98% was achieved. A 150 µl volume of the inoculating fluid was placed into each well on the plate.

Finally, to test for a key N-metabolic function, we performed urease assays on a subset of cultured bacterial strains. Briefly, bacteria were inoculated into Rapid Urea Broth (Becton Dickinson, Sparks, MD, USA) containing phenol red, a pH indicator. Urease-induced production of ammonia (NH_3_) from urea renders the culture media alkaline, leading to a resulting color change of red to bright purple.

### General functional genomic analyses

In our first genomic test of functional dissimilarities between *Cephalotes* larval and adult gut symbiont strains, we generated a matrix of symbiont gene counts and their assignments to Kyoto Encyclopedia of Genes and Genomes (KEGG) Orthology identifiers (KOs), normalized by the total count of KO-encoding genes, for the 18 MAGs and 14 IGs. We used this matrix for Non-Metric Multi-Dimensional Scaling (NMDS) analysis and for a Permutational Multivariate Analysis of Variance (PERMANOVA) in R v3.6.3 [88], applying the ‘vegan’ and ‘ggplot2’ packages [89, 90], to test whether the overall sets of functions encoded by individual *Cephalotes* symbionts differ between strains isolated from larvae *vs.* adults.

Next, in order to identify functions differentially enriched in larval *vs.* adult symbiont communities, we extracted bacterial gene counts per KEGG pathway [91] for the *C. varians* PL010 larval and adult worker gut metagenomes from IMG/M-ER [79], using these to calculate *Z*-normalized Log Odds Ratios (*Z*-LOR). Significance was evaluated by applying the False Discovery Rate correction of Benjamini-Hochberg [92], with a critical threshold value of 0.05. This approach differed from the first by focusing on all bacterial genes in our *C. varians* PL010 larval and adult gut metagenomes (no IGs), which included those from MAGs and all genes on scaffolds not assigned to MAGs.

### Full pathway genomic explorations

The challenges of functional genome annotation have been well-documented within the field of genomics [93–95]. To alleviate some of these issues, we operated beyond automated annotation pipelines, developing a methodological framework to screen and curate gene annotations from genomes and metagenomes for genes and pathways involved in fiber catabolism, N-recycling, amino acid biosynthesis, B-vitamin biosynthesis, organic acid metabolism and sulfur metabolism (**S4 Fig**).

To begin, we first searched *C. varians* PL010 larval and adult gut metagenomes, and all 14 *Cephalotes*-derived IGs, by entering KO and Enzyme Commission number (EC) assignments as keywords in the IMG/M-ER Function Search tool, repeating this for all steps of focal KEGG pathways falling within the six aforementioned metabolic functions. With hundreds of genes yielded from these Function Searches, we screened annotation evidence for each gene, seeking to retain those with modest-to-high confidence assignments to the metabolic and transport steps of interest. To achieve this, all search-returned genes with KOs or ECs matching those used in our targeted search (*i.e.* and, hence, most likely to encode the focal step) were kept for further analysis. For Function Search-returned genes with KO and EC annotations that did not correspond to the targeted pathway steps, we examined COG and Pfam annotations, keeping genes when their annotations matched those expected for the targeted pathway steps, and discarding all others.

Following the guilt-by-association assumption, stating that in prokaryotic genomes, clustered genes often share related functions [96, 97], we examined functional annotations of genes found in proximity, on the same genomic scaffolds as our Function Search-returned genes. For each of our metabolic pathways screened, we first identified scaffolds with more than three search-returned genes with related pathway step annotations on a single scaffold. For these, gene annotations on the entire scaffolds were visually inspected in the IMG/M-ER Chromosome Viewer, and BLASTp analyses were conducted against the NCBI Non-Redundant (nr) protein database for at least three genes located on each side of the search-returned genes, when available. Additional genes of interest discovered through these BLASTs were added to our tables of Function Search-returned genes.

For fiber catabolism and N-recycling pathways (reviewed in [98, 99]), deemed to be of known [58], or likely, importance for these symbioses, we examined the reliability of functional annotations in IMG/M-ER by comparing these to results from two BLASTp searches performed for each retained search-returned gene – one against the SWISS-PROT database and the second against the NCBI nr database.

Three exceptions to the approach detailed in the previous paragraphs were made. (1) Due to the large number of steps in biosynthetic pathways for the 20 genetically encoded amino acids, Function Search analyses for these metabolisms used only KOs (*i.e.* not ECs). (2) To supplement our assessment of fiber and other carbohydrate catabolism in larvae and adults from *C. varians* colony PL010, the two gut metagenomes were additionally screened against the HMMER, DIAMOND and Hotpep databases for Carbohydrate Active Enzyme (CAZy) families (here referred to as ‘CAZymes’) [100], using the dbCAN2 metaserver (v7, January 2019) [101]. Catabolic CAZymes found in these metagenomes were categorized based on the type of substrates the enzymes can degrade from a review of their CAZypedia descriptions or CAZyme activities in the CAZy website [100]. (3) To expand the taxonomic breadth of hosts assessed for the beneficial function of N-recycling, our Function Search analyses targeted worker gut metagenomes from 5 additional *Cephalotes* species (*C. grandinosus*, *C. maculatus*, *C. minutus*, *C. pusillus*, *C. rohweri*) [58]. These 5 turtle ant species were chosen because they diverged in excess of 20, and in some cases, 40 million years ago, hailing from several divergent lineages of the *Cephalotes* phylogeny [102]. These species were also included in the 16S rRNA amplicon sequencing survey of Hu et al. [66], giving us functional insight into microbiomes characterized at a compositional level.

### Screening for plant and fungal cell wall degrading enzymes and other carbohydrate degrading enzymes across *Cephalotes* species

To further examine symbiont-encoded digestive capacities, we screened for 48 plant and fungal cell wall degrading enzymes (PFCWDEs) and other carbohydrate degrading enzymes across symbiont (meta)genomes from multiple turtle ant species (**S4 Table**). The enzymes on this list were identified by (i) examining KEGG pathway steps for key aspects of carbohydrate metabolism, and (ii) by including enzymes screened in other studies investigating digestive roles of social insect symbionts [15, 103]. After identification, KO, EC, COG and Pfam annotations for the targeted PFCWDEs were imported into IMG/M-ER as a Function Set. An IMG/M-ER Function Profile analysis was then performed to display the genes encoding any of these 48 PFCWDEs and other carbohydrate degrading enzymes across adult worker gut metagenomes from the six *Cephalotes* species studied above for N-recycling (*C. grandinosus*, *C. maculatus*, *C. minutus*, *C. pusillus*, *C. rohweri*, and *C. varians* (PL010)), the larval gut metagenome from *C. varians* colony PL010, and the 14 *Cephalotes* IGs.

Focusing on 16 key PFCWDEs that are involved in the depolymerization of fibers expected to be of significance in turtle ant diets, we attempted to find additional instances of these enzyme-encoding genes in a broader set of *Cephalotes* species (*i.e.* 11 additional species represented with a metagenome in IMG/M-ER [58]), while also assessing the possibility of false positive results. We used sequences of genes annotated as the 16 key PFCWDEs from the 6 focal *Cephalotes* species as queries in IMG/M-ER BLAST searches against all *Cephalotes* metagenomes and IGs. To filter out false positives, BLASTp searches were first performed using these same queries against custom NCBI Identical Protein Group databases, in April-July 2019. After aligning top hits with our queries and the sequences retrieved from IMG/M-ER BLASTs against the 11 additional *Cephalotes* species, we performed phylogenetic analysis using RAxML Black Box [86] on the CIPRES Science Gateway [87] to generate maximum likelihood/rapid bootstrapping trees. Visual assessments of single protein phylogenetic trees constructed in the interactive Tree Of Life website (iTOL) [104] were then performed as shown in **S5 Fig**. When sequences from *Cephalotes*-derived symbionts clustered with sequences previously annotated to encode other enzymes – *i.e.* not the expected PFCWDEs – the functional annotation of symbiont-encoded genes was judged to be not well supported. Such genes were not included in the final version of the trees.

Screening results, trees for the phylogenetic analyses and R scripts are saved as data repository, available at 10.6084/m9.figshare.15173730.

## Results

A summary of the assembly statistics for the metagenomes, metagenome-assembled genomes (MAGs) and cultured isolate genomes (IGs) used in our study can be found in **S1-S3 Tables**.

### 1. Taxonomic and functional overviews

#### 1.1. Bacterial community composition in larval and adult worker guts of *Cephalotes varians*

The *C. varians* larval gut metagenome from colony PL010 was dominated by bacteria from the Enterobacteriales, Lactobacillales and Rhizobiales orders (Fig 1A). Among these bacterial groups, only members of the Rhizobiales are known as core symbionts of *Cephalotes* adult worker guts, with those from the Enterobacteriales and Lactobacillales being largely confined to larval guts [66]. Also found in the PL010 larval gut metagenome, though at lower relative abundance, were bacteria from the Burkholderiales, Xanthomonadales and Pseudomonadales orders, which are common core symbionts of *Cephalotes* adults.

Rare, if not absent, from PL010 larvae were relatives of the *Staphylococcus* sp. (Bacillales) isolated from *C. texanus* larval guts (**S6 Fig**). *Staphylococcus* bacteria did, however, comprise averages of ∼1.0 and 1.6% in the larvae of single colonies of *C. varians* and *C. texanus*, based on 16s rRNA amplicon data [66]. Being found also in larvae of *C. palta* (1%), and seemingly rarer or absent from the remaining 14 colonies in that study, these bacteria, thus, appear to be occasional residents of the turtle ant larval microbiome.

Relatives of *Enterococcus* sampled from our *C. varians* larval gut PL010 metagenome – exhibiting high relatedness to *Enterococcus faecalis* – were more common than *Staphylococcus* associates in the 16S rRNA amplicon study across larvae of 12 turtle ant species, with >1% average relative abundance in larvae from seven out of the 17 studied colonies and a maximum of 16% within larvae of *C. setulifer* [66]. Through that method, *Enterococcus* comprised up to 4.7% of the microbiome when averaged across *C. varians* larvae of single colonies [66].

#### 1.2. MAGs of larvae in relation to cultured isolate genomes and MAGs of adults

Assembled scaffolds from the PL010 larval gut metagenome annotated as belonging to the three larval dominant bacterial orders were binned into 7 MAGs (**S3 and S5 Tables**), including two assigning to the Enterobacteriales (MergB2 and MergB4), two from the Lactobacillales (Bin3 and Bin6), and three from the order Rhizobiales (Bin10, Bin5.8.r1 and Bin5.8.r2). These added to the 11 MAGs previously generated from the PL010 adult worker metagenome [58], and to the 12 adult and 2 larval cultured isolate genomes (IGs). Large proportions of scaffolds from several MAGs were classified to lower taxonomic levels with good confidence, based on IMG/M-ER analysis. For example, Burkholderiales Bin12 was likely to be a bacterium from the Comamonadaceae family, with 89% of Bin12 scaffolds assigning to this family. Burkholderiales-assigned Bin3.1 and Bin5.1 were more likely to be from the Alcaligenaceae family, with 65% and 71% their scaffolds respectively assigning to this family (**S7 Fig; S5 Table**). For larval symbionts, MergB2 was deemed likely to represent an *Enterobacter* sp., with 97% of its scaffolds assigning to this genus. Sixty-one percent of MergB4 assigned to the *Klebsiella* genus, suggesting a more tentative assignment for this MAG. The larva-derived Bin6, introduced in the previous section, hailed from the *Enterococcus* genus, with 99% of its scaffolds classifying accordingly. The similarly larva-derived Bin3 was, alternatively, concluded to belong to the genus *Lactococcus*, as supported by the assignment of 77% of the Bin3 scaffolds (**S6 Fig; S5 Table**).

Results from pairwise average nucleotide identity computation indicated first that the cultured *Enterococcus* symbiont of *C. texanus* (JR029-101) was closely related to MAG Bin6 from *C. varians*. Given the high pairwise orthoANI value (98%), the larva-derived symbionts from these two turtle ant species are likely from the same bacterial species (*i.e.* pairs of genomes with orthoANI >95% [105]). Second, two Rhizobiales species appeared to be shared between *C. varians* larvae and adults (**S8 Fig**). One of them was the adult-derived IG JR021-5 and the larval MAG Bin10, with over 96% nucleotide identity across their genomes. The second pair included the adult-associated MAG Bin13 and the larva-derived MAG Bin5.8.r2, which were over 99% identical. This indicates that, while largely distinct, microbiomes in adults and larvae do show some overlap.

#### 1.3. Overall predicted metabolic functions in larval *vs.* adult worker microbiomes

Differences in functional investment of turtle ant gut bacterial symbionts from the two developmental stages were measured as the differences in normalized counts of KO-encoding genes. Overall, predicted functional capabilities of symbionts from larval guts were significantly different from those represented in adult worker guts (PERMANOVA: 9,999 permutations, *F_1,31_* = 2.44, *P*-value <0.01; Fig 1B). In a non-metric multi-dimensional scaling (NMDS) analysis focused on IGs and MAGs, core symbionts that are common in adults, and in some cases, also larvae (*i.e.* Rhizobiales), loosely clustered together on the second axis, at the exclusion of larvae-specific symbionts from the Enterobacteriales and Lactobacillales (Fig 1B). Bacteria from the same taxonomic orders showed a strong tendency to cluster on both axes, with an exception exhibited for Rhizobiales, due to functional divergence of the JR021-5/Bin10 species from the remaining Rhizobiales representatives.

To explore in greater depth the predicted dissimilarities in metabolic investment between larval and adult turtle ant gut microbiomes, we compared bacterial gene counts for KEGG metabolic pathways in *C. varians* PL010 larval and adult gut metagenomes. Among the ten KEGG pathways that were significantly more abundant in the larval gut metagenome (*Z*-LOR >2.97, adjusted *P*-value <0.05; Fig 1C; **S6 Table**) were four carbohydrate metabolism pathways involved in the metabolism of fibers and simple carbohydrates, such as xylan (hemicellulose) and chitin (KEGG pathway: ‘Amino sugar and nucleotide sugar metabolism’), fructose and mannose (‘Fructose and mannose metabolism’), cellulose and starches (‘Starch and sucrose metabolism’) and galactose (‘Galactose metabolism’). Three of the remaining six pathways related to membrane transport and two-component systems. Overall, KEGG pathways significantly more enriched in the larval gut metagenome included transporters (‘ABC transporters’, Phosphotransferase system’), differing components of carbohydrate and sulfur metabolism, defensive functions (‘beta-Lactam resistance’, ‘Cationic antimicrobial peptide’), and sensory mechanisms (‘Two-component system’). The most abundant genes in the larval microbiome were encoding an iron complex transporter, which had the highest number of gene copies in the larval gut metagenome (*fhuD*: 68, *fhuB*: 87, *fhuC*: 62; **S7 Table**).

In contrast, the adult gut metagenome was significantly enriched in eight KEGG pathways involved in fatty acid, amino acid, other carbohydrate, and carboxylate metabolism (*Z*-LOR <-2.95, adjusted *P*-value <0.05; Fig 1C; **S6 Table**). Many of the enriched genes in the adult gut metagenome encoded central, sometimes versatile, metabolic enzymes functioning in several KEGG pathways (**S7 Table**). For example, the most encoded KO in the adult metagenome was for the enoyl-CoA hydratase (K01692, EC:4.2.1.17, 221 gene copies; **S7 Table**), a central metabolic enzyme that catalyzes a step in the beta-oxidation of fatty acids to produce FADH_2_ and acetyl Coenzyme A (acetyl-CoA). This enzyme can participate in seventeen different KEGG pathways. Other key adult-enriched bacterial enzyme-encoding genes included the acetyl-CoA synthetase (K01895, EC:6.2.1.1), acetyl-CoA C-acetyltransferase (K00626, EC:2.3.1.9) and acetyl-CoA carboxylase (K01961, EC: 6.4.1.2), all involved in the conversion of acetate and acetyl-CoA into substrates for the oxidative tricarboxylic acid cycle (oTCA) or fatty acids.

Taken together, these results suggest that larvae-enriched bacterial predicted functions, with investment in transporters, carbohydrate and sulfur metabolism, and microbial defenses, could be characteristic of microbes regularly interacting with a threatening, but sometimes fiber- and simple carbohydrate-rich, environment. In contrast, adult-enriched bacterial predicted functions, with investment in central carbon, amino acid, fatty acid and organic acid metabolism, may be symptomatic of a more confined system, where derivatives of dietary substrates (*e.g.* organic acids, some amino acids and fatty acids) are commonly converted into useful metabolites like amino acids or stored as fatty acids.

### 2. Targeted screenings of metabolic capabilities in larval *vs.* adult worker microbiomes in turtle ants

To disentangle symbiont impacts and for evidence that these gut symbionts play similar roles to those of symbionts in other animal systems, including B-vitamin and amino acid biosynthesis, and the metabolism of both sulfur and organic acids, we measured the completeness of these biosynthetic and metabolic pathways encoded in the 32 MAGs and IGs from *Cephalotes* larval and adult worker guts.

#### 2.1. B-vitamin and amino acid biosynthesis

Several insects with plant-based diets depend on their microbial symbionts to acquire essential B-vitamins [67, 106] and amino acids [107]. Among the 32 analyzed MAGs and IGs of *Cephalotes*, we found that gut symbionts collectively encode the genetic capacities to completely synthesize almost all B-vitamins (Fig 2A; **S9 Fig**). In particular, we found that the MAG of a larva-associated *Klebsiella* sp. (MergB4; Enterobacteriales) encoded complete pathways to synthesize all B-vitamins except thiamine (B1) and cobalamin (B12). Also of note was the finding that *Ventosimonas* symbionts (Pseudomonadales) from adult worker guts encode complete, or near-complete, pathways for B-vitamins including riboflavin (B2), pantothenate (B5), pyridoxine (B6), biotin (B7) and folate (B9).

**Fig 2.**
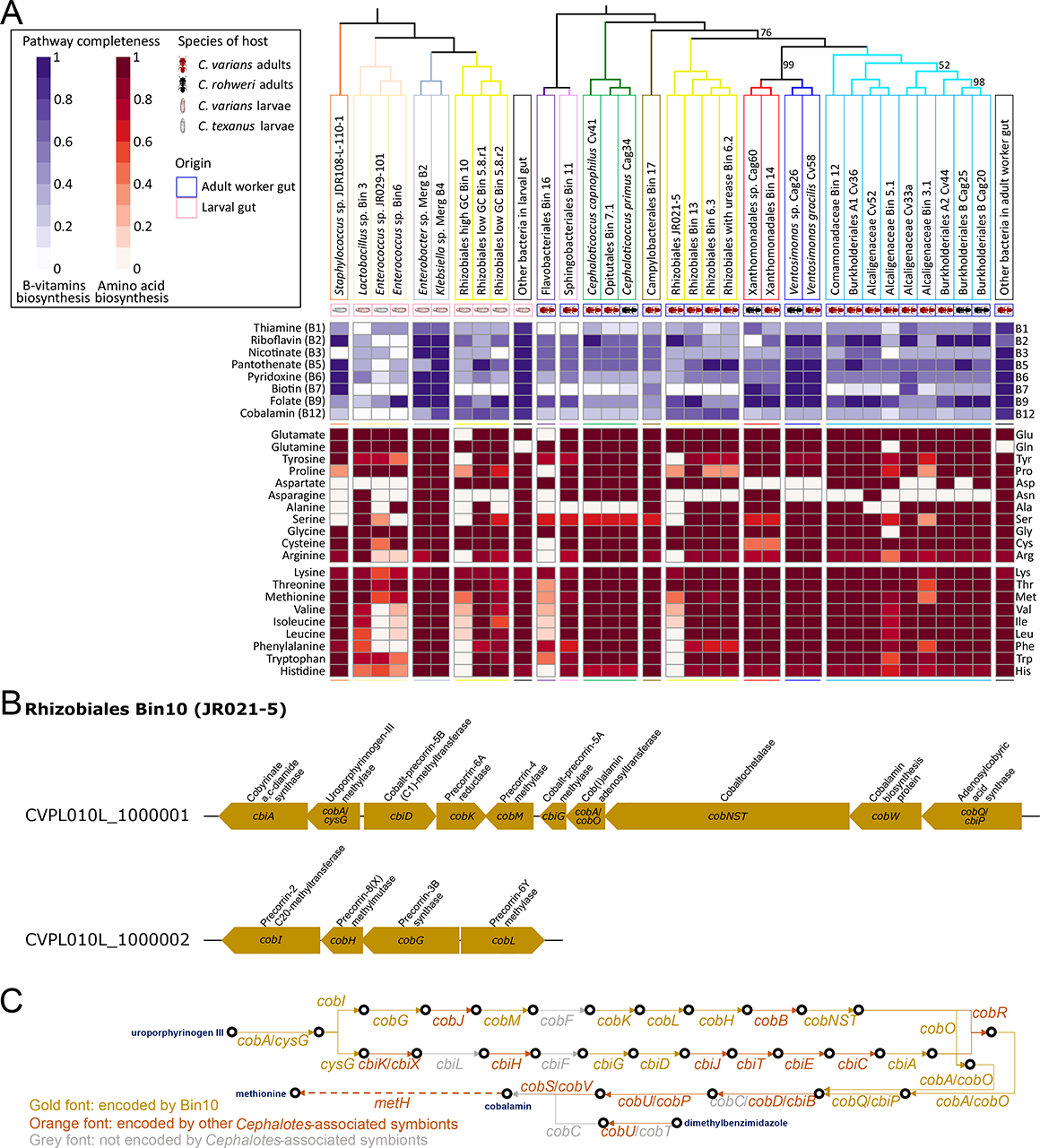
B-vitamin and amino acid biosynthesis pathways encoded in adult and larval *Cephalotes*-associated symbiont genomes. **(A)** Heatmap depicting the completeness of biosynthetic pathways encoded in genomes of distinct *Cephalotes*-associated bacterial gut symbionts. Proportions of step categories encoded in symbiont genomes (columns) to perform various metabolisms (rows) is given by the shade within the squares of the heatmap, the darker the more complete the biosynthetic step is. Symbiont genomes were screened in IMG/M-ER for the presence of key enzyme-encoding genes. Bacterial genes from the *C. varians* PL010 metagenomes that were not binned to Metagenome-Assembled Genomes (MAGs, with the mention “Bin” or “Merg”) but having a predicted relevant function are pooled in the “Other bacteria in larval gut” and “Other bacteria from adult worker gut” columns, respectively. Additional cultured Isolated Genomes (IGs) from *C. varians*, *C. texanus* and *C. rohweri* were included. The bacterial phylogenetic trees were inferred based on seven bacterial marker gene sequences using a maximum likelihood (ML) method and 999 bootstrap iterations. Bootstrap values <100 are shown on the tree. B-vitamin detailed pathways encoded by *Cephalotes*-associated symbionts can be found in **S9 Fig**. **(B)** Cobalamin (B12-vitamin) gene clusters encoded in the larva-derived Rhizobiales Bin10 MAG (JR021-5). **(C)** Pathways for cobalamin production encoded by Bin10 or other *Cephalotes*-associated symbionts. The pathway following the *cob* route, is encoded in *Hodgkinia* genomes from cicadas [108], leading to cobalamin-dependent production of methionine.

Encoding more than 50% of the cobalamin (B12) biosynthetic pathway, the Rhizobiales symbionts from larvae and adults possessed clusters of genes coding for the production of this B-vitamin (Figs 2B and 2C). Along with *Klebsiella* sp. MergB4, which similarly appeared to lack a few cobalamin pathway steps, it is possible that Rhizobiales symbionts synthesize pseudo-cobalamin corrinoids, with activities related to cobalamin. In fact, as seen for the genomes of *Hodgkinia* endosymbionts of cicadas [108], the *metH* gene coding for a cobalamin-dependent version of methionine synthase (EC:2.1.1.13) was encoded by all *Cephalotes*-associated Rhizobiales genomes, except for JR021-5/Bin10 (Fig 2C). Also encoding this gene was the adult-enriched *Cephaloticoccus capnophilus* (Opitutales), and two larva-derived bacteria, *Enterobacter* sp. MergB2 and *Klebsiella* sp. MergB4. This indicates that these symbionts may utilize cobalamin-like cofactors to synthesize methionine.

Nearly all core symbionts from adult turtle ants encode complete pathways to synthesize essential and non-essential amino acids [58]. With an interest in whether adult symbionts are unique in these properties we expanded our search of amino acid biosynthesis in larval symbionts. Newly discovered here was the possession of genes required to synthesize all amino acids in *Klebsiella* sp. MergB4 from larval guts, and a majority of amino acids in other larval-associated symbionts (Fig 2A). MergB4 was predicted to synthesize amino acids that subsets of individual adult-associated symbionts cannot produce on their own, including, for example, alanine, serine, and histidine that *Cephaloticoccus* sp. cannot make, or aspartate, arginine and cysteine that adult-associated Xanthomonadales cannot synthesize. It is, thus, conceivable that N-metabolism of abundant larval symbionts could make contributions to colony-level N-budgets.

#### 2.2. Sulfur and coupled metabolism

As encoded in the *Blochmannia* symbiont genome from carpenter ants [109] and of use in the production of sulfur-containing amino acids, many symbionts from the larval and adult *Cephalotes* guts possessed the genes to import and reduce extracellular sulfate down to sulfide, using the assimilatory pathway to synthesize cysteine and methionine (Figs 3A and 3B). Some symbionts appeared to lack just one step of the pathway. For example, in Alcaligenaceae Cv33a (Burkholderiales), only the gene for adenylyl-sulfate kinase (*cysC*: EC:2.7.1.25) was missing, while *Cephaloticoccus capnophilus* appeared to have a complete set of genes for this pathway (Fig 3C). Other symbionts could potentially use alternative routes to produce sulfide: the *C. texanus* larval *Staphylococcus* sp. JDR108L-110-1 may utilize homocysteine and the adult Burkholderiales Cv44 and Cag20 might reduce extracellular taurine (Fig 3B).

**Fig 3.**
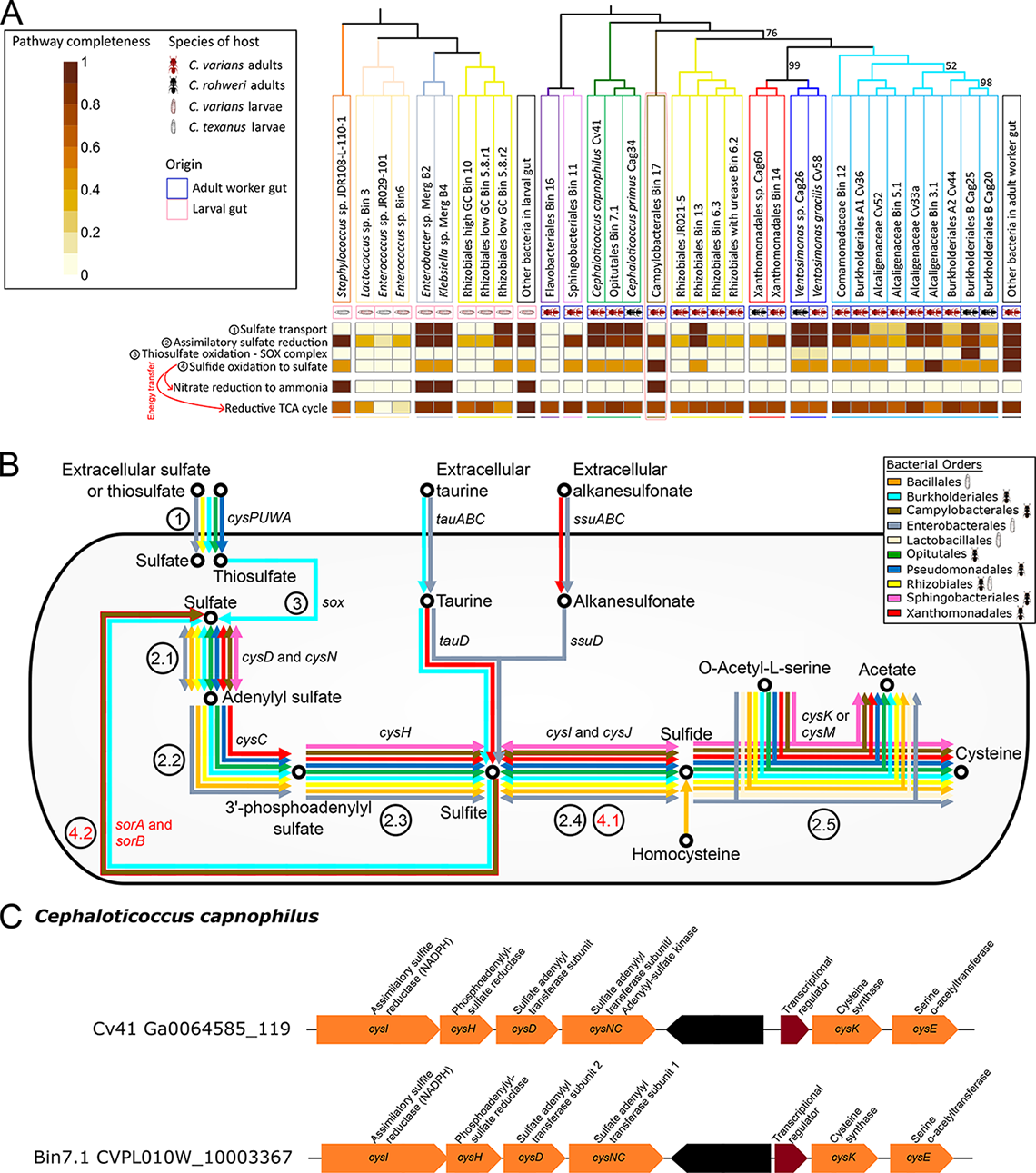
Sulfur metabolism encoded in adult and larval *Cephalotes*-associated symbiont genomes. **(A)** Heatmap depicting the completeness of sulfur metabolism pathways encoded in genomes of distinct *Cephalotes*-associated bacterial gut symbionts, as well as the nitrate reduction to ammonia pathway and the reductive TCA (rTCA) cycle, which could be coupled with sulfide oxidation. See legend from Fig 2A for a complete description of the heatmap. **(B)** Sulfur metabolism pathways encoded in *Cephalotes*-associated genomes. Colors of the arrows correspond to the taxonomic order of the bacterial symbionts possessing the genes coding for the focal steps. Note that many *Cephalotes*-associated symbionts have retained genes for sulfate reduction, but few can also oxidize sulfide. **(C)** Sulfate reduction gene cluster from two symbionts associated with *C. varians*. The scaffolds contain multiple genes coding for different subunits and enzymes involved in sulfate reduction. Note that the gene coding for a sulfate adenylyltransferase subunit 1 in *Cephaloticoccus capnophilus* Bin7.1 MAG matches a putative adenylyl-sulfate kinase-encoding gene in Cv41 IG.

Seemingly unique to the gut of adult turtle ants when compared to larvae is the potential to generate sulfate in two specific oxidative ways. In the first case, the genome of adult-associated Burkholderiales Cag25 encodes enzymes that could use thiosulfate in the sulfur-oxidation system (SOX) route (Fig 3B). Another route could be taken by Alcaligenaceae Bin3.1 (Cv33a strain; Burkholderiales) and Campylobacterales Bin17, which appear to encode capacities to oxidize sulfide to sulfate in only two energy-generating steps (Figs 3B and 4). Biological sulfide oxidation is typically conducted in oxic/anoxic interfaces (high/no dioxygen (O_2_) concentrations) with high levels of sulfide [110]. It is not yet known if such microaerobic to anaerobic conditions are common in portions of *Cephalotes* guts. Indeed, representatives from nearly all major adult-symbiont lineages, with a notable exception of midgut-enriched Campylobacterales [111], have been grown under aerobic conditions [58,112–114], raising further questions on the context for use of sulfide-oxidizing genes by these specialized bacteria.

**Fig 4.**
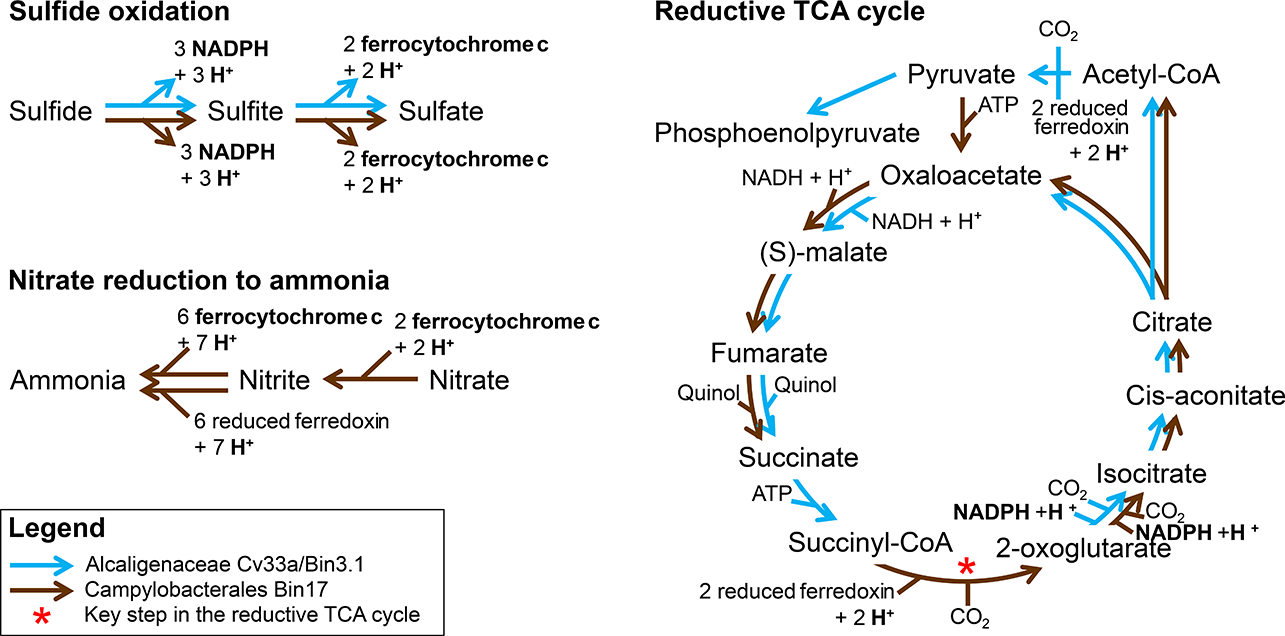
Sulfide oxidation and coupled pathways encoded in two sulfide oxidizers from *Cephalotes varians*. Only the Campylobacterales Bin17 MAG can complete both nitrate reduction and the reductive TCA (rTCA) cycle, by using electron donors generated by sulfide oxidation. The red asterisk indicates an essential step to complete the rTCA cycle.

Sulfide oxidative processes in prokaryotes are commonly coupled with nitrate reduction or carbon dioxide (CO_2_) fixation, providing electron donors that participate in these energy-demanding reductive reactions [115]. For example, free-living members of the *Arcobacter* oxidize sulfide while either reducing or denitrifying nitrate and fixing CO_2_ [116–118]. In light of this, and the fact that *Arcobacter* species are the closest relatives of specialized *Cephalotes*-associated Campylobacterales symbionts [58], we investigated the genomic potentials for coupling of sulfide oxidation with other metabolic mechanisms potentially encoded by Campylobacterales Bin17. To achieve this, we expanded our targeted screenings to two new metabolic pathways commonly utilized by this symbiont’s free-living relatives: nitrate reduction to NH_3_ and the reductive (*i.e.* reverse) TCA (rTCA) cycle for CO_2_ fixation. In adult turtle ant guts, Campylobacterales Bin17 was the only symbiont with the genetic potential to reduce nitrate to NH_3_ via an anaerobic respiratory dissimilatory pathway (Figs 3A and 4). In larval guts, this pathway was encoded by the two Enterobacteriales MAGs MergB2 and MergB4 and by the *C. texanus* larval *Staphylococcus* sp. JDR108L-110-1 IG. The larvae-associated bacteria do not appear to couple this nitrate reduction pathway with sulfide oxidation, as the latter pathway is incomplete in the genomes of these bacteria (Fig 3A).

Many enzymes from the oTCA cycle are reversible and can serve in CO_2_ fixation via the rTCA cycle [119–121]. In our screenings, multiple symbionts from turtle ant guts encoded reversible steps of both the rTCA and oTCA cycles (Figs 3A and 4). Among them, both of the candidate sulfide oxidizers from adult guts, Alcaligenaceae Bin3.1/Cv33a and Campylobacterales Bin17, encoded nine out of eleven steps of the rTCA cycle. Only Bin17, however, appeared to encode 2-oxoglutarate synthase (*i.e.* 2-oxoglutarate:ferredoxin oxidoreductase; EC:1.2.7.3), a key step in the rTCA cycle (Fig 4) [122, 123].

Altogether, these results indicate that many symbionts, including the midgut-dominant *Cephaloticoccus* sp., appear to have the genetic capability to produce sulfide via assimilatory sulfate reduction. Our genetic evidence further suggests that this metabolite is, then, either used for amino acid synthesis or made available to other bacteria with the capacities for sulfide oxidation. If the midgut microenvironment where Campylobacterales Bin17 settles is one of low/no dioxygen, this symbiont could oxidize sulfide, obtained directly from another midgut bacterium, *Cephaloticoccus* sp. [111]. In the process this symbiont would produce electron donors such as NADPH or ferrocytochrome c, which could fuel nitrate reduction to NH_3_ and the conversion of CO_2_ to organic carbon, molecules via the rTCA cycle, by this same symbiont.

#### 2.3. Organic acid metabolism

Organic acids are produced by mammalian gut microbiomes, and serve as a source of symbiont-derived energy in a number of social insects [98]. Through our analyses we ascertained that such molecules – including lactate, acetate, formate, and citrate – can be produced from pyruvate and/or further metabolized by a variety of turtle ant symbionts (Fig 5A; **S10 Fig**). Production of butyrate and propionate through pyruvate fermentation pathways could also be possible, though only if completed by a collaboration of symbionts, each encoding portions of the full pathway. Only in larvae did we find evidence of symbiont capacities to produce formate from pyruvate fermentation via formate acetyltransferase (EC:2.3.1.54). Among such symbionts were larva-associated Enterobacteriales and Lactobacillales. Encoding the potential to derive lactate, formate, or acetate through the fermentation of pyruvate, bacteria from these groups, including *Enterobacter* sp. MergB2, *Enterococcus* sp. JR029-101/Bin6, Lactococcus sp. Bin3, *Klebsiella* sp. MergB4 and *Staphylococcus* sp. JDR108L-110-1, hence exhibited genomic similitudes with their free-living and pathogenic relatives [124–126]. Although many symbionts in both larvae and adults had the genomic potential to oxidize formate to CO_2_ and H^+^ using formate dehydrogenase (EC:1.17.1.9), only some larva-isolated Enterobacteriales could grow on formate (**S8 Table**). Importantly, this NAD-dependent formate dehydrogenase is reversible. Therefore, it could possibly be utilized by Campylobacterales Bin17 and Alcaligenaceae Cv33a/Bin3.1, as well as other members of the turtle ant microbiomes, to reduce CO_2_ into formate, as an alternative means of fixing CO_2_ [127].

**Fig 5.**
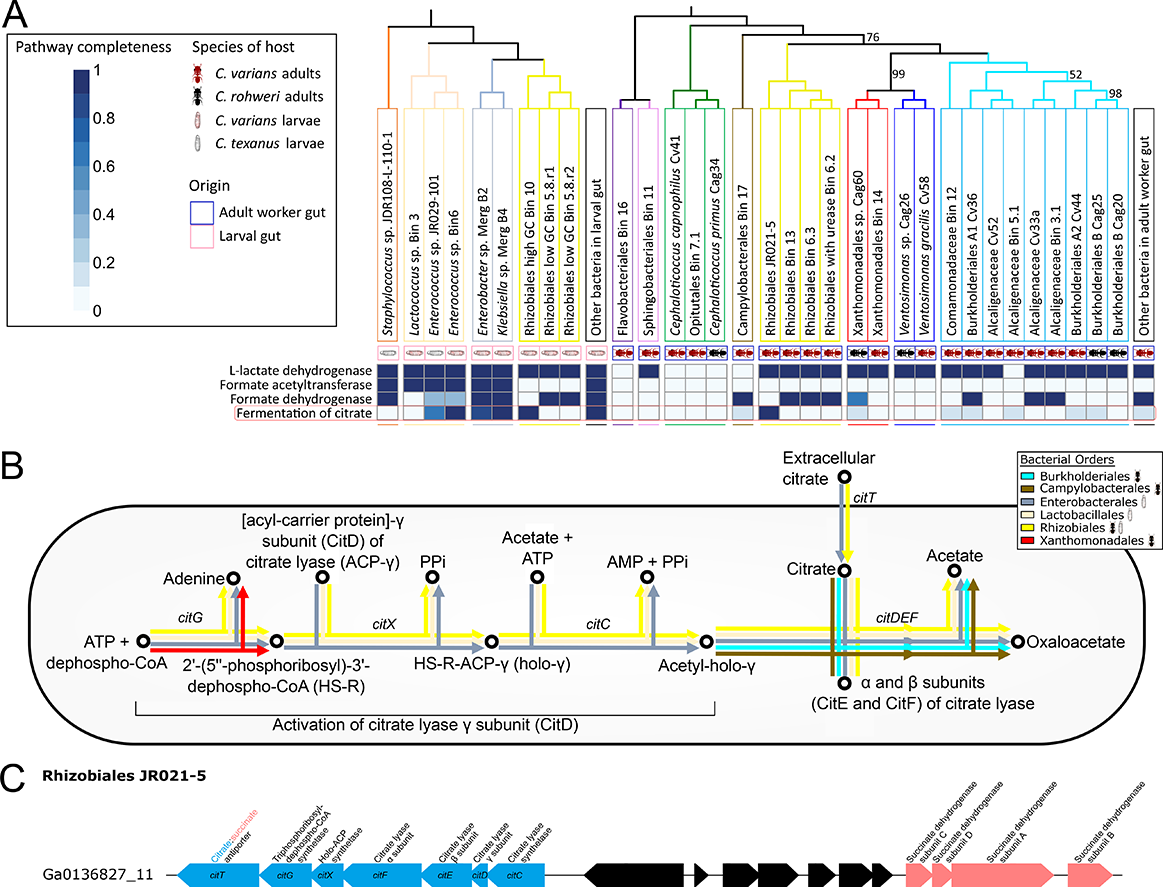
Organic acid metabolic enzymes and pathways encoded in adult and larval *Cephalotes*-associated symbiont genomes. **(A)** Heatmap depicting the presence of genes encoding key enzymes and pathways in organic acid metabolisms encoded in genomes of distinct *Cephalotes*-associated bacterial gut symbionts. See legend from Fig 2A for a complete description of the heatmap. Organic acid pathways encoded by *Cephalotes*-associated symbionts can be found in **S10 Fig**. **(B)** Citrate fermentation pathway encoded in *Cephalotes*-associated genomes. Colors of the arrows correspond to the taxonomic order of the bacterial symbionts possessing the genes coding for the focal steps. **(C)** Gene cluster from Rhizobiales JR021-5 IG including genes to ferment citrate. The scaffold contains multiple genes coding for different subunits of citrate lyase and other enzymes involved in citrate fermentation. A citrate:succinate antiporter-encoding gene is also found in this cluster, in proximity of genes coding for succinate dehydrogenase.

Larva-specific and Rhizobiales symbionts possessed the genes coding for the three citrate lyase subunits (*citDEF*: EC:4.1.3.6) essential to ferment citrate into acetate and oxaloacetate, as well as for the citrate transporter-encoding genes (*citT*) and for the activation of citrate lyase (*citG*: EC:2.4.2.52, *citX*: EC:2.7.7.61, and *citC*: EC:6.2.1.22) (Fig 5B). It is interesting to note that multiple larva-derived symbionts (plus some adult-isolated symbionts) could grow on citrate, *in vitro*, under oxic conditions (**S8 Table**). All genes coding for citrate transport and fermentation were found in a cluster on a scaffold from the IG of Rhizobiales JR021-5 (Fig 5C), one of the two genome-sequenced symbiont species that is clearly shared between larvae and adults (**S8 Fig**). This cluster was similar to the *cit* gene cluster from various Enterobacteriaceae [128, 129]. The citrate transporter-encoding gene (*citT*) was annotated as a citrate:succinate antiporter [129], which coincides with, on the same scaffold, a cluster of genes coding for all the subunits of succinate dehydrogenase, involved in the reversible production of succinate. Our results show that only turtle ant larval symbionts can ferment pyruvate to formate, but both life stages host gut symbionts that have the genetic potential to ferment pyruvate into lactate and acetate, and share the putative citrate-fermenting Rhizobiales, which may be of use in pollen digestion [130].

### 3. N-recycling in larval and adult turtle ants

N is difficult to access for turtle ants, given that, in their diet, it is either physically well-protected by a tough cell wall, like pollen [48, 131], or found in animal waste-based recalcitrant molecules, including uric acid or urea [44,49,132] (Fig 6A). Because immature insects require N to a greater extent than adult insects [133, 134], we hypothesized that turtle ant larval symbionts would encode genes for N-fixation and/or recycling of N wastes. In adult turtle ants, microbial-driven fixation of atmospheric dinitrogen was not found to be a mechanism that could compensate this lack of easily accessed N [58]. Similarly, we found that in the *C. varians* PL010 larval gut metagenome, none of the *nif*, *anf*, and *vnf* genes involved in N-fixation was present. N-recycling, in contrast, was recently found to be an important microbial contribution to turtle ant adult nutritional requirements [58].

**Fig 6.**
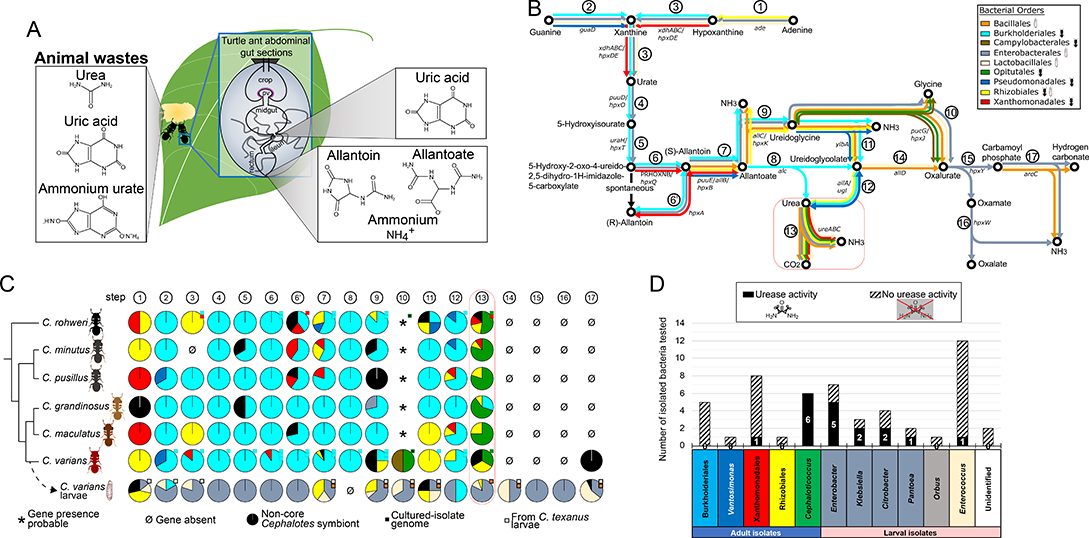
Nitrogen- (N-)recycling by turtle ant larval and adult gut symbionts. **(A)** N sources from the *Cephalotes* diet. The diet of turtle ants was historically posited as deficient in available N [55]. Primary sources of N in turtle ant’s diets include wind-dispersed pollen [171], with internal protein and amino acids difficult to access [131], droppings from birds, reptiles and insects, and mammalian urine [44, 49]. While mammalian, reptilian and avian wastes mainly contain urea, uric acid and ammonium urate, respectively [191–193], invertebrate excreta can be diverse, and composed of allantoin, allantoate and ammonium in ants [194]. Sources of N for turtle ants could also include their own waste, as adult-to-adult proctodeal trophallaxes (*i.e.* anal-oral trophallaxes [195]) are common behaviors in these ants [132]. **(B)** N-recycling pathways encoded by larval and adult *Cephalotes* gut symbionts. Colors of the arrows correspond to the taxonomic order of the bacterial symbionts possessing the genes coding for the focal steps. Numbers and gene names associated with the arrows link to the pathway steps from **S11 Fig**. **(C)** Symbiont-encoded N-recycling across *Cephalotes* species. The pie charts show the proportion of genes from bacteria identified at the taxonomic order (colors) to code for specific steps of N-recycling (columns) in given *Cephalotes* gut metagenomes (rows). The small squares indicate whether a gene coding for a given enzyme was found in IGs identified at the order level by their color. Squares with thick black borders correspond to IGs from *C. texanus* larvae. The *Cephalotes* phylogeny is adapted from [102]. Asterisks indicate the probable presence of genes coding for (S)-ureidoglycine-glyoxylate aminotransferase from the abundant *Cephaloticoccus* sp. (Opitutales) associated with all *Cephalotes* species screened to date [58], and not restricted to *C. varians*. **(D)** Results of *in vitro* urease assays conducted on cultured bacteria isolated from *C. varians* and *C. rohweri* adult guts and bacteria from *C. varians* and *C. texanus* larval guts. Each assay was run in triplicates. Full results of *in vitro* metabolic assays can be found in **S8 Table**.

#### 3.1. Uricolytic pathways

To infer microbial contribution to N-recycling in larval guts and to explore extended N-recycling pathways in adult worker gut microbiomes, we applied both metagenomic screening on (meta)genome-derived data and urease assays on cultured symbiont isolates. One key finding was the discovery of the *puuD* gene in a member of the adult-derived Comamonadaceae (Burkholderiales), encoding an enzyme (EC:1.7.3.3) that catabolizes uric acid (Fig 6B; **S11 Fig**) [135]. The encoding symbiont Comamonadaceae Bin12 emerges as the second uric acid degrading symbiont in the adult turtle ant system, joining Cv33a -a previously identified, symbiont from the related Alcaligenaceae [58].

Newly discovered in the larval microbiome, from MAGs of *Enterobacter* sp. MergB2 and *Klebsiella* sp. MergB4 (Enterobacteriales), was the possession of all genes required to produce uric acid from purines, to recycle uric acid into urea, and then NH_3_ (Fig 6B; **S11 Fig**). This NH_3_ could then be assimilated into the process of amino acid synthesis, since these larval-associated symbiont MAGs encoded all genes required to synthesize 18 to 20 of the protein-building amino acids (Fig 2A).

Interestingly, the uricolytic route taken by these Enterobacteriales is different from the one taken by the adult Alcaligenaceae Cv33a. In particular, although conserved in adults among *Cephalotes* species (Fig 6C), the Cv33a allantoicase-encoding *alc* genes (producing urea and ureidoglycolate from allantoate; EC:3.5.3.4) were absent from larval gut symbionts (Figs 6B and 6C; **S11 Fig**). Gene screening results suggested, instead, that allantoate could alternatively be utilized by MergB2 and MergB4, in addition to other larval and adult symbionts, to directly produce CO_2_, NH_3_ and ureidoglycine. Some symbionts, including the MergB2 and MergB4, appeared then capable of hydrolyzing ureidoglycine into glycine and oxalurate, which they could use to produce oxamate or carbamoyl phosphate, followed by NH_3_.

Interestingly, adult-associated *Cephaloticoccus capnophilus* and Campylobacterales Bin17, both abundant in the midgut of *C. varians* [111,136,137], possessed the genes encoding (S)-ureidoglycine-glyoxylate aminotransferase (EC:2.6.1.112), to produce glycine and oxalurate from ureidoglycine (Figs 6B and 6C; **S11 Fig**), thus unlocking an alternative exit for the uricolytic route that had not been recognized in a prior study on *Cephalotes* adult-associated symbionts [58].

Almost all of the larval Enterobacteriales genes coding for the uricolytic pathway were grouped on one scaffold from the MergB2 MAG, and four were grouped on scaffolds binning to the MergB4 MAG (**S12 Fig**). The structural organization of these genes was highly similar to the *hpxA-Z* supraoperonic cluster from *Klebsiella oxytoca* M5al and *K. pneumoniae* MGH78578 (corresponding to MergB2 and MergB4, respectively) previously documented [138]. This spatial proximity likely facilitates the co-expression of these functionally related genes [97]. Genes coding for allantoin and allantoate permeases found in these clusters indicate that exogenous sources of N may be integrated into the uricolytic pathway of these Enterobacteriales, and/or that the Enterobacteriales may release intermediates of the uricolytic pathway in the larval gut lumen. The larval *Enterococcus* sp. JR029-101/Bin6 and *Staphylococcus* sp. JDR108L-110-1 may also participate in intermediate uricolytic steps since they encode the genes to utilize allantoin to produce ureidoglycine, glycine and oxalurate (Figs 6B and 6C; **S11 and S12 Figs**).

#### 3.2. Urea hydrolysis

Urease-encoding genes, involved in the hydrolysis of urea into NH_3_ and CO_2_, were previously found in the adult-derived *Cephaloticoccus capnophilus* (strain Cv41), *Cephaloticoccus primus* (strain Cag34), Xanthomonadales sp. strain Cag60 and a Rhizobiales symbiont represented by MAG Bin6.2 ([58]; Figs 6B and 6C; **S11 and S12 Figs**). In the present study, we also found genes coding for the three urease subunits in the larval *Staphylococcus* sp. JDR108L-110-1 and the MergB2 and MergB4 MAGs. Assays on larva-derived symbionts from the Enterobacteriales (*Enterobacter*, *Klebsiella*, *Citrobacter* and *Pantoea*) revealed common, but non-ubiquitous instances of urease capacities, as well, since 10 out of 16 larval isolates from this bacterial order were positive for *in vitro* urease activity (Fig 6D; **S7 Table**). In addition, one *Enterococcus* sp. isolated from a larval gut out of 12 tested successfully degraded urea *in vitro*, suggesting occasional appearance of this function within this group.

### 4. Fiber digesting potentials in larval and adult turtle ants

Turtle ants have regularly been observed consuming plant materials, such as pollen, nectar, plant wound secretions, as well as collecting fungi and chewing on lichens (Fig 7A) [43,44,48,49,139]. These food sources are rich in carbohydrates, especially in recalcitrant fibers that may be degraded with the participation of symbiont digestive enzymes, like in honeybees and fungus-growing termites [52, 140]. Beyond nutritive functions, since some reports indicate that turtle ants may excavate the inside of their woody nest [141, 142], symbiont digestive enzymes could also conceivably support the modification of recalcitrant lignocellulose that composes wood. Because in ants, larvae are key digestive and liquefying components for the colony [39–41], we hypothesized that the gut of turtle ant larvae was the site of intense plant and fungal fiber digestion. However, because (i) in other ants active digestive enzymes can passage through the adult guts [143] and be deposited on fiber substrates via fecal droplets, and (ii) some turtle ant adult microbes are found in larval guts (Fig 1A; [66]), there is possibly a potential for digestive enzyme encoding genes in the adult microbiome, whether of use in this stage or not.

**Fig 7.**
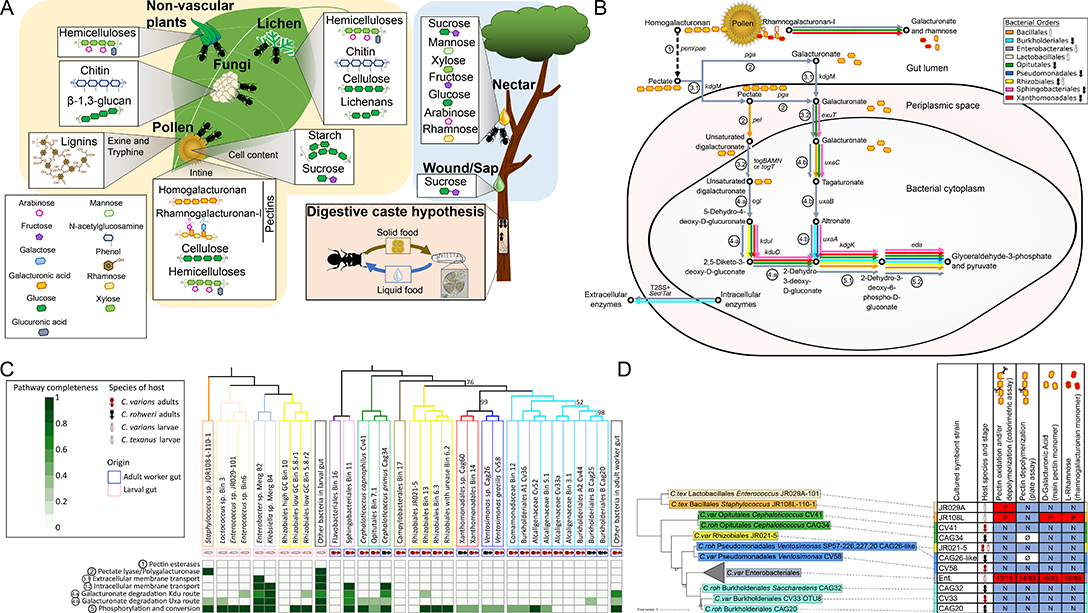
Dietary pectin catabolism by gut symbionts in turtle ants. **(A)** Characteristic food items collected by turtle ants that are sources of fibers and simple carbohydrates. Turtle ants are cryptic herbivores, described as plant foragers and exudate feeders [98]. Their diet is relatively rich in carbohydrates. In ants, while adult workers collect foods, the essential steps of digestion, at least of solid foods, are believed to be conducted by larvae. This is the ‘digestive caste hypothesis’ [39–41]. Adult foragers deliver solid food to larvae which process it, and, after assimilating essential nutrients, regurgitate more digestible elements to elder members of the colony. Pollen grains partially degraded were retrieved in *Cephalotes varians* larval guts, as depicted by the light microscopy image. **(B)** Detailed pathway for pectic homogalacturonan catabolism in the *C. varians* PL010 adult and larval gut metagenomes, starting from homogalacturonan and ending with pyruvate and glyceraldehyde-3-phosphate with two possible routes for galacturonate catabolism. Degradation steps encoded by at least one gene in the metagenome are indicated by big colored arrows. The colors of the big arrows correspond to bacterial orders for genes encoding given steps. Taxonomic annotation was performed in IMG/M-ER. Dotted arrows correspond to steps that were not found encoded in the metagenome. **(C)** Heatmap depicting the presence of genes encoding key enzymes in homogalacturonan catabolic pathways encoded in genomes of distinct *Cephalotes*-associated bacterial gut symbionts. See legend from Fig 2A for a complete description of the heatmap. **(D)** Results of *in vitro* pectinase assays conducted on cultured bacteria isolated from *C. varians* and *C. rohweri* adult guts and bacteria from *C. varians* and *C. texanus* larval guts. Each assay was run in triplicates. The phylogenetic tree was built from 16S rRNA sequences extracted from individual isolates using a ML method and 999 bootstrap iterations. Bootstrap values <100 are shown on the tree. Full results of *in vitro* metabolic assays can be found in **S8 Table**.

#### 4.1. Pectic homogalacturonan catabolism

Pectin, a multi-polymeric fiber mainly composed of homogalacturonan, is abundant in pollen intine [144–146]. Homogalacturonan catabolic pathways commonly used by bacteria (*i.e.* from the depolymerization of the fiber to the conversion into commonly used metabolites) were reconstructed and analyzed in detail in the *C. varians* PL010 larval and adult gut metagenomes. Although our metagenomic screenings returned enterobacterial scaffolds from the larval gut metagenome that possessed genes coding for a pectin methylesterase (EC:3.1.1.11), our phylogenetic verification methods indicated that they instead likely encoded an acyl-CoA thioester hydrolase, with no activity on pectins, as previously reported [147]. Since esterification level is variable in pectic homogalacturonan [148], with pectins from pollen walls showing low degrees of methyl-esterification [149], the absence of pectin methylesterase may not inhibit effective depolymerization of this fiber.

More definitive were findings from larval gut metagenome scaffolds, classified by IMG/M-ER as from the Enterobacteriales, which encoded the potential to depolymerize pectate (*i.e.* non-esterified homogalacturonan) using a polygalacturonase (EC:3.2.1.15) (Fig 7B). Bacteria from this order also encoded each of the type II secretion system subunits that would enable the transport of this enzyme into the extracellular environment [150]. And they further encoded the genes enabling import and transformation of homogalacturonan-derived galacturonate into pyruvate and glyceraldehyde-3-phosphate, through at least one pathway (*i.e.* the Uxa route).

Larval Lactobacillales possessed genes to import unsaturated digalacturonate, but two steps were missing to complete the catabolism of this compound (Fig 7B). In contrast to the larval gut microbiome, none of the bacteria in *C. varians* adult guts possessed genes to depolymerize homogalacturonan or pectate (Figs 7B and 7C), although Sphingobacteriales Bin11 symbionts appeared to be missing just one step of galacturonate degradation.

One cluster of genes coding for homogalacturonan degradation was identified from the larval gut metagenome (**S13 Fig**). Falling on a scaffold from the MergB4 MAG (Enterobacteriales), this grouping contained pectate transporter genes (*kdgM* and *togMNAB*) and genes for enzymes catalyzing the degradation of galacturonate (*kduD* and *ogl*).

Overall, *in vitro* pectinase assays performed on larva- and adult-derived symbionts supported our metagenomic results. First, none of the bacteria isolated from the adult gut and tested for *in vitro* pectinase activity could depolymerize (9 isolates tested) or oxidize (23 isolates tested) homogalacturonan (Fig 7D; **S8 Table**). In contrast, 24 out of 126 (19%) bacteria from the larval gut, including 81% of the larva-isolated Enterobacteriales and 67% of the larva-isolated *Enterococcus*, could depolymerize homogalacturonan. Among the larval isolates, 16 were capable of oxidizing homogalacturonan but not depolymerize this compound. These included 2 *Citrobacter*, 6 *Enterobacter*, 4 *Enterococcus*, 2 *Klebsiella*, 1 *Pantoea* and 1 *Staphylococcus* isolates.

#### 4.2. Plant and fungal cell wall degrading enzymes (PFCWDEs) and other carbohydrate degrading enzymes in *C. varians* metagenomes

To infer symbiont contributions to the catabolism of dietary fibers (including fiber depolymerization and conversion of fiber derivatives, like di- and oligosaccharides), we first screened *C. varians* PL010 larval and adult whole gut metagenomes against the specialized carbohydrate active enzyme (CAZy) database [100], focusing on plant and fungal cell wall degrading enzyme (PFCWDE) and other carbohydrate degrading enzyme families with known activity against fibers (and their derivatives) of predicted abundance in the turtle ant diet [96] (Fig 7A). In the larval gut metagenome, we identified 42 CAZy families (CAZymes) likely involved in fiber catabolism, including glycoside hydrolases (GHs), polysaccharide lyases (PLs), carbohydrate esterases (CEs), and those with auxiliary activities (AAs). Genes encoding such enzymes collectively comprised 0.44% of all the genes in the PL010 larval gut metagenome (547 genes). In the adult gut microbiome from the same *C. varians* ant colony, we found genes encoding 44 CAZymes likely involved in fiber catabolism. The 763 encoding genes accounted for a slightly lower percentage, 0.12%, of the more deeply sampled and/or gene-diverse adult gut microbiome (**S9 Table**). Thirty-one CAZymes were encoded by both larval and adult metagenomes in this studied *C. varians* PL010 colony. While some CAZymes were encoded by various symbionts in both larvae and adults (*e.g.* AA3, GH23, GH13), others were encoded by only a few symbiont taxa (**S14 Fig**), indicating possible symbiont specialization to utilize specific carbohydrates.

#### 4.3. Catabolic pathways of fiber-derived carbohydrates

Following this cataloging of PFCWDEs and other carbohydrate degrading enzymes encoded in turtle ant larval and adult whole guts, we assessed symbiont coding capacity within common bacterial catabolic pathways hypothesized to aid the use of recalcitrant fibers of potential interest for turtle ants (**S15 Fig**). At the whole gut community level, both larval and adult gut metagenomes included symbiont-derived genes for the complete catabolism of most predicted dietary fibers (**S16 Fig**). Two exceptions were homogalacturonan (Figs 7C and 7D) and arabinogalactan that could only be depolymerized (step ‘C1’) by larval symbionts (**S16 Fig**). Overall, bacteria from the larval gut metagenome, especially those encoding scaffolds that were not binned into MAGs, possessed genes to complete most of the catabolic steps of presumed dietary fibers. And in particular, when focusing on MAGs from larval guts, the MergB2 and MergB4 genomes were found to encode genes for complete catabolism of cellulose, xylan (hemicelluloses), resistant starch, chitins and lignins.

Bacteria from the adult worker gut metagenome, including those with genomes not binned into MAGs, possessed genes to catabolize a good array of fibers such as cellulose, xylan, resistant starch, chitins and possibly lignins (**S16 Fig**). This repertoire included putative laccases, or multi-copper oxidase-encoding enzymes – though ambiguous annotations made it unclear as to whether these were involved in bacterial cell wall metabolism or lignin degradation. Clearer, in several cases, were the annotations from many of the unbinned metagenomic bacterial genes, which coded for beta-xylosidase (degrading xylan), chitinase, and alpha-amylase. Genes encoding these capacities were mostly identified from scaffolds assigned to the Xanthomonadales order, with all such genes falling on *Cephalotes*-specific branches in our phylogenetic analyses (**S17 Fig**).

Results from *in vitro* metabolic assays supported many of our genomic inferences, in that several larval and adult symbionts could grow on cellulose-derived cellobiose and starch-derived amylose and maltose, as well as *N*-acetylglucosamine, a chitin monomer (**S18 Fig; S8 Table**). The alpha-amylase function encoded by members of the Xanthomonadales was successfully recapitulated *in vitro*, via strain Cag62 showing amylase activities. Both cultured *Cephaloticoccus* strains (Cv41, Cag34) catabolized cellobiose and maltose *in vitro*. Genome sequencing of the *C. rohweri*-isolated *Cephaloticoccus primus* (Cag34) suggested that this latter function was likely enacted through a maltase enzyme (EC:3.2.1.20; **S16 Fig**). Like the *Cephaloticoccus* isolates, Rhizobiales JR021-5 also catabolized maltose *in vitro*, albeit through an unidentified mechanism.

#### 4.4. Retention of symbiont-encoded PFCWDEs across *Cephalotes* species

To expand the above work, we used IMG/M-ER Function Profile screenings and protein phylogenetic analyses to assess the distributions of PFCWDEs across six divergent *Cephalotes* species. Microbiomes from the six focal congeners of *C. varians* were populated by bacteria with comparable potentials to: (1) depolymerize or debranch arabinosides from hemicellulose or from rhamnogalacturonan-I of pectin, to (2) depolymerize cellulose, chitins, xylan, fructans, resistant starch and amylose, and to (3) potentially oxidize lignins. The taxonomy of the bacteria encoding these specific functions showed modest conservation across these ants, in spite of the millions of years since their common ancestry (Fig 8).

**Fig 8.**
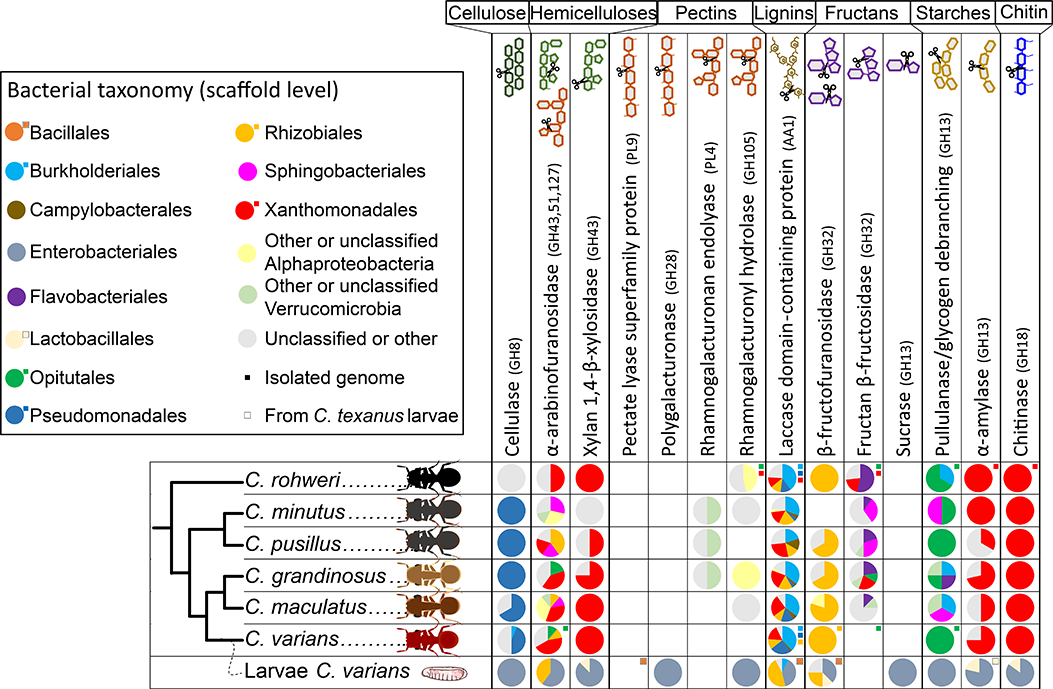
Summary of plant and fungal cell wall digestive enzymes (PFCWDEs) encoded by bacteria in IMG/M-ER metagenomes from adults of six *Cephalotes* species and a larval metagenome. The pie charts show the proportion of genes from bacteria identified at the taxonomic order (colors) to code for specific enzymes (columns) in given *Cephalotes* hosts (rows). The small squares indicate whether a gene coding for a given enzyme was found in IGs identified at the order level by their color. Squares with black borders correspond to IGs from *C. texanus* larvae. The *Cephalotes* phylogeny is adapted from [102].

Although none of the adult-associated symbionts from the six turtle ant species studied here encoded genes to depolymerize homogalacturonan, a major ingredient of most pectins, BLAST searches revealed that bacteria of other *Cephalotes* species, not targeted with our Function Profile screening, possessed polygalacturonase-encoding genes. Such bacteria, however, were often at very low abundance (**S19 Fig**). Interestingly, genes from larval-associated *Staphylococcus* and Enterobacteriales encoding pectate lyase and polygalacturonase, respectively, did not group into *Cephalotes*-specific clades (**S5 Fig**), showing close relatedness instead to those of free-living bacteria.

Xanthomonadales bacteria from the six adult worker gut metagenomes possessed genes, that if expressed, would impart remarkable capacities, being the sole group of adult *Cephalotes*-associated bacteria with genes coding for the depolymerization of xylan and chitins. With just one potential exception, these capacities were conserved, for this taxon, across the six focal turtle ant species as determined through our IMG/M-ER Function Profile screening (Fig 8; **S17 Fig**).

Some *Cephalotes*-specific patterns arose for the enzymes encoding depolymerization of fructans, homogalacturonan, and rhamnogalacturonan-I. This latter molecule is the second most abundant form of pectin, and is depolymerized via multiple enzymes, including two detected through our (meta)genomic analyses (Fig 8). First, rhamnogalacturonan endolyase was encoded by scaffolds assigning to the Verrucomicrobia – *i.e.* the parent phylum of *Cephaloticoccus* sp. – in 3 out of the 6 focal *Cephalotes* species (*C. grandinosus, C. minutus* and *C. pusillus*). These scaffolds possessed similarly high read depths and GC% values in comparison to the dominant *Cephaloticoccus* sp. from the same metagenomes, raising the possibility that they belonged to members of this specialized, symbiont lineage (**S19 Fig**). Second, rhamnogalacturonyl hydrolase-encoding genes were enriched in *C. rohweri* guts (but also found in other *Cephalotes* species), hailing from scaffolds assigned to the Sphingomonadales (a non-core symbiotic group from the Alphaproteobacteria) and Verrucomicrobia. Genomes from two *C. rohweri*-isolated strains, Xanthomonadales Cag60 and *Cephaloticoccus primus* Cag34, also encoded rhamnogalacturonyl hydrolase enzymes (**S19 Fig**).

In addition to these two rhamnogalacturonan-degrading enzymes, an aforementioned adult-specific homogalacturonan-depolymerizing polygalacturonase was encoded by a clade of Verrucomicrobia which had relatively low abundance, potentially derived from, or closely related to, *Cephaloticoccus* spp. Based on the combined results of our gene screening and BLAST searches, verrucomicrobial polygalacturonase-encoding genes were found in the gut of only 4 out of 17 turtle ant species with a sequenced metagenome: *C. minutus*, *C. pallens*, *C. persimplex* and *C. umbraculatus* (**S19 Fig**). Protein phylogenetic trees for these three pectinolytic enzymes in adult microbiomes revealed monophyletic groupings of adult-associated enzymes, without close relatedness to homologs from free-living bacteria. This suggests that their encoding genes have been retained by specific conserved symbionts, possibly including *Cephaloticoccus*spp. in some proportion of *Cephalotes*species.

Despite consistent evidence for annotations suggesting numerous PFCWDEs in adult and larval symbionts, some preliminary results from gene screening analyses were rendered unlikely or ambiguous, upon follow-up examination – as exemplified by cellulase-encoding genes. While strongly evidenced in larval symbionts (**S20 Fig**), detection in adult-associated *Cephaloticoccus* sp. was thrown into question on the basis of phylogenetics, revealing their putative cellulases to group with peptidases, rather than cellulases. Another set of potential cellulases was classified to the Pseudomonadales, an order including core adult-enriched *Ventosimonas* symbionts (Fig 1A; **S8 Fig**) [58, 112]. But examination of the cellulase-associated gene cluster showed architectural similarities with the *Pseudomonas fluorescens* SBW25 *wss* cellulose-synthetase operon, documented previously [151] (**S20 Fig**). While the focal gene was, hence, likely a cellulase-encoding gene, the evidence did not support a digestive function for the predicted cellulase, but instead a role in the regulation of bacterial cellulose biosynthesis, which may be used in the formation of biofilms or in adherence to host tissues.

## Discussion

In this study, we investigated the role of symbionts from turtle ant larval and adult worker guts using a combination of genomic screening and *in vitro* metabolic assays. We show that, despite the differences in taxonomic composition of adult *vs.* larval gut microbiomes, coinciding with apparent diverging genetic investment in overall metabolic potentials, both stages harbor gut symbionts with the genetic machinery to contribute to digestive and nutritional functions for their ant hosts. Our results unveil a clear functional overlap between adult and larval gut microbiomes in key traits of significance to host dietary ecology (N-recycling, dietary fiber degradation, B-vitamins, amino acids and organic acids production, and sulfate reduction). Only a few metabolic pathways appear to be encoded by stage-specific symbionts, like pectic homogalacturonan processing and formate production being specific to larval gut symbionts and sulfide oxidation only encoded by symbionts from the adult gut. While the functional landscape of *Cephalotes* larval guts is dominated by metabolically versatile members of the Enterobacteriales, the adult gut symbiont community displays several patterns of functional complementarity, with variable levels of symbiont specialization.

### Specificities of the *Cephalotes* larval gut community

Larvae play a crucial role in food digestion for ant species with solid diets [40, 41]. Yet, digestive and nutritional functions of ant larval gut symbionts have rarely been explored [but see 152]. Here, we investigated symbiont genomes from larval gut bacteria with high relatedness to symbionts found in abundance in late-stage turtle ant larvae [66]. For a subset of these bacteria, we also performed *in vitro* assays to test genetic predictions on the metabolic potential of cultured isolate symbionts. Through these efforts we document the putative functions of a turtle ant larval gut microbiome for the first time.

We show that symbionts of turtle ant larvae have the genetic potential to assist their hosts through the processing of fungal- and plant-based foods, and to synthesize a variety of essential nutrients, like amino acids, B-vitamins and simple carbohydrates. Likely responsible for these metabolic capabilities in the larval gut microbiome are abundant representatives from the Enterobacteriales, possessing a diverse metabolic machinery. Genes encoded by these bacteria were highly related to those of bacteria from a diversity of non-ant gut habitats (**S5, S17 and S20 Figs**). Combined with analyses in a separate study [66], these trends suggest that such microbes – in addition to at least some Lactobacillales – have not been domesticated as symbiont specialists, unlike the majority of the adult microbiome. Evidence instead suggests that these bacteria are recurrently acquired, quite possibly, from their food [153, 154].

Beside Enterobacteriales-encoded metabolic pathways of potential value to the host, including B-vitamin synthesis, sulfate and nitrate reduction, N-recycling, and amino acid production, their contributions to pectic homogalacturonan depolymerization and catabolism (Fig 7A) may be key to the acquisition of simple pectin-derived carbohydrates. This activity may also, or alternatively, be crucial to the disassembly of pollen intine, yielding access to nutrient-rich internal pollen contents.

In the gut of pollen-feeding adult honeybees, pollen-derived homogalacturonan is depolymerized by gammaproteobacterial *Gilliamella* symbionts [15, 52], which are then able to catabolize pectin-derived galacturonate [155] and ferment such dietary carbohydrates into organic acids [156], useful metabolites for other honeybee symbionts and for the host [51]. We thus hypothesize that the colonization of the turtle ant larval guts by Enterobacteriales enables efficient digestion of recalcitrant pollen grains, a known staple food for turtle ants, which is given to larvae as infrabuccal pellets [48]. Given the greater likelihood for a digestive function of cellulase activity encoded by larva-derived Enterobacteriales (**S20 Fig**), and the seemingly patchy ability of adult symbionts to degrade diverse pectins (Fig 8), we would speculate that these bacteria may indeed bring a broader range of PFCWDE repertoires to the site of solid food digestion. Like a number of other larva-derived symbionts (*e.g.* members of the Rhizobiales, the *Staphylococcus* sp. JDR108L-110-1), Enterobacteriales encode the formate dehydrogenase enzyme (Fig 5A), which may be important in the catabolism of single carbon compounds and generation of energetic NADH molecules, or in certain forms of anaerobic (*e.g.* nitrate) respiration. Research on the O_2_ concentration across different microhabitats within larval and adult guts (as briefly alluded to elsewhere in this manuscript), will be illuminating for this system.

Abundant Enterobacteriales may cooperate or compete with the less abundant, but common, larvae-associated Lactobacillales, which were also found to encode potentially mutualistic functions in the larval gut. Unlike Enterobacteriales, these Gram-positive bacteria do not appear genetically capable of B-vitamin synthesis, sulfur metabolism, or – for two of three strains with sequenced genomes (IGs or MAGs) – amino acid biosynthesis. While their plant and fungal fiber degradative capacities appear more limited, a subset of Lactobacillales in *C. varians* seem capable of catabolizing chitin, xylan (hemicellulose), starches and cellulose-derived cellobiose. Their PFCWDE-encoding genes are closely related to those of a number of non-symbiotic strains in this order (*e.g.* **S17 Fig**). While Hu et al. [66] showed that a subset of Lactobacillales fall into *Cephalotes*-specific lineages, a subset exhibits close relatedness to free-living bacteria based on sequence similarity at 16S rRNA genes. This raises the possibility that, like Enterobacteriales, some of these symbionts are acquired from the environment.

Based on our genome inferences, larvally-localized Lactobacillales symbionts may also engage in forms of anaerobic metabolism, important in energy generation. This is partially evidenced through their encoding of formate acetyltransferase (Fig 5A), an enzyme that generates formate and acetyl-CoA in the presence of pyruvate. It is also evidenced, for one IG, through the possession of genes for citrate fermentation (Fig 5B). This latter property is encoded by other larva-associated Enterobacteriales, and at least some members of the third, larva- dominant group – the Rhizobiales.

### The shared Rhizobiales symbionts

Symbionts from the Rhizobiales contrast with the two other larva-abundant taxa because they can dominate the foregut (*i.e.* infrabuccal pocket, crop and proventriculus) of adult *C. varians* and *C. rohweri*, and can also be found in the hindgut of these adult ants [26,136,137]. They differ, also, through their status as core members of the adult gut microbiome, which form seemingly specialized, *Cephalotes*-specific groups on phylogenies [65, 157]. Showing relatedness to *Bartonella* sp., and including the provisionally described genus *Tokpelaia,* members of this group are among the most prevalent ant-associated bacteria, enriched in some herbivores, but recorded in many unrelated ants with various diets [56,65,152,157–160]. We found here that two Rhizobiales species are shared between turtle ant adult workers and larvae. These symbionts could thus be exchanged between larvae and adults via food transfer or oral-oral trophallaxes.

Strain JR021-5 is of particular interest because it has the smallest genome size among all analyzed *Cephalotes*-derived symbionts sequenced thus far (**S2 Table**). It appears shared between larval and adult guts, based on its high relatedness to Bin10 Rhizobiales MAG from larvae (**S8 Fig**). As proposed in Hu et al. [66], this symbiont and members of other specialized, core *Cephalotes* clades, may be passaged across generations by colony founding queens, and then perpetuated to larvae, and eventually workers, through trophallactic behaviors.

The JR021-5/Bin10 symbiont has the genetic capability to ferment citrate to acetate (Fig 2A). Recently, in another ant-symbiont system, citrate was found to be catabolized by Entomoplasmatales sampled from leafcutter ant guts, likely via fermentative action, which supposedly contributes to the digestion of the host’s fruit-rich diet [161]. Citrate could also be a pollen derivative, or produced as a downstream metabolite from enzymatic pathways catabolizing pollen-derived carbohydrates. Interestingly, genes involved in citrate fermentation in two honeybee-isolated *Lactobacillus* strains are activated in the presence of pollen [130]. The JR021-5/Bin10 symbiont also has the genetic ability to synthesize riboflavin and cobalamin-like corrinoids. These biosynthetic pathways are similarly encoded by the somewhat distantly related alphaproteobacterial *Hodgkinia* symbionts of cicadas [108], but also by mammal-associated gut bacteria [162].

Although more research is needed to infer turtle ant host dependency upon Rhizobiales symbionts, their potential citrate fermentation, and amino acid and B-vitamin biosynthetic functions may be beneficial to such herbivorous insects, feeding on specialized, nutrient-depleted diets [106]. Rhizobiales are more abundant in adult guts when turtle ants feed on a pollen-rich diet [62], indicating that the symbionts might benefit from such a diet. Whether the generation of energy-rich organic acids, through processes like the citrate fermentation pathway, is of key importance for turtle ants’ dietary ecology and a topic in need of further study. Pollen-feeding honeybees and wood-feeding termites present a precedent for symbiont-mediated short-chain fatty acid and other organic acid provisioning [14, 51], revealing the plausibility that *Cephalotes* could similarly benefit from such services.

### Metabolic potentials of the adult gut community in *Cephalotes* ants

Due to the scarcity of usable forms of N in several components of the herbivorous turtle ant diet, symbiont recycling of N wastes, like uric acid and urea, was hypothesized as a key role to their evolutionary success [60, 65]. Bacterial recycling of diet-derived urea into amino acids, followed by assimilation by the adult ant host, was recently demonstrated via isotope labeling experiments [58, 163]. Genomic and metagenomic inferences supported by *in vitro* analyses identified functionally conserved lineages of symbionts as N-recyclers, including *Cephaloticoccus* sp. and Alcaligenaceae strain Cv33a [58]. Yet, further metabolic screening of the adult turtle ant gut community to explore other symbiotic roles was justified, given the imposing diversity and density of these symbionts – a trait found to be uncommon among ants [59].

Expanding on previous work, we here document potential new roles for the turtle ant adult worker gut symbiont community. Beyond the N-recycling function, we show that, at the gut symbiont community level, and compared to the aforementioned larval gut microbiome, which is partially composed of free-living bacterial symbionts, the conserved adult worker gut microbiome is enriched in genes for central, fatty acid and amino acid metabolisms (Fig 1C), with particularly high numbers of genes encoding acetyl-CoA metabolism. We found that some of the most encoded genes in the adult worker gut microbiome are involved in the conversion of acetyl-CoA to/from fatty acids, acetate and pyruvate. Some of these enzymes were encoded by many of the symbionts from adult guts (**S10 Fig**), indicating that acetyl-CoA may be a key metabolite to symbionts residing in the gut of these adult ants. It is also worth noting that acetyl-CoA can be synthesized from pantothenate [164], which can potentially be produced by various larva- and adult-derived symbionts (**S2 Fig**). Acetyl-CoA is involved in many metabolic and biosynthetic processes, such as the energy-yielding oTCA cycle, the fatty acid metabolism and both amino acid and B-vitamin syntheses. Although more research is needed to determine whether symbiont-driven acetyl-CoA metabolism impacts host physiology, it could possibly be involved in increased fatty acid biosynthesis, elongation and storage, which can be linked to insect weight gain [165]. It could, alternatively, be involved in ant cuticle formation [166] or inter-individual chemical communications [167].

With regards to metabolic potentials of individual symbionts screened in the present study, the distributions of the newly explored functions suggest that these ancient symbionts may occupy unique metabolic niches within the micro-ecosystems of the turtle ant gut compartments, similar to the ancient gut bacterial symbionts of herbivorous honeybees [52], termites [168] and possibly passalid beetles [169]. These results contrast with the previously identified near-ubiquity of amino acid biosynthesis among core turtle ant adult-enriched symbionts [58]. Illustrating several novel functions of interest are the urea-recycling *Cephaloticoccus* sp. [58, 113]. Rare in turtle ant larvae [66], modestly abundant in the adult ileum, and dominant in the adult midgut [26,136,137], these bacteria are now predicted to utilize cellulose-derived cellobiose and to deconstruct recalcitrant fibers like resistant starch. In some host species, they may even catabolize pectic homogalacturonan and rhamnogalacturonan-I, which are key pollen compounds [146].

Other members of the adult-associated microbiome also have the potential for diverse metabolisms, including fiber-degrading capabilities, which could aid host digestion. Members of the relatively abundant Xanthomonadales order [58] (Fig 1A) – sampled mainly from the adult turtle ant ileum [111], and rarely from larvae (**S3 Fig;** [66]) – can degrade multiple dietary fibers, including those found in different pollen components, like xylan, arabinosides and amylose (Fig 8). These symbionts also encode chitinases, potentially involved in the digestion of fungal or insect cuticle. This set of enzyme-encoding genes in the Xanthomonadales may allow symbiont-assisted digestion of dietary fibers, with potentially ancient conservation given that the most recent common ancestor of our studied turtle ants lived 46 million years ago [102]. Indeed, in most protein phylogenies, the PFCWDEs encoded by *Cephalotes*-associated Xanthomonadales fell on long branches, far removed from enzymes encoded by non-symbiotic bacteria (**S17 Fig**). Similar patterns were seen for other putative digestive enzymes encoded by specialized members of the adult microbiome, with several trends suggesting long-standing specialization in this proposed mutualistic service. Conservation of symbiont roles was, for example, documented for N-metabolism, with a uricolytic pathway enriched-lineage of Burkholderiales (Fig 6C), and with the ubiquity of urease activity across *Cephaloticoccus* spp. [58].

Despite these trends, we also report here, from our analyses of adult worker gut metagenomes, some sporadic distributions of symbiont-encoded PFCWDEs across turtle ants, hinting at more labile roles for particular symbiont lineages – and in a few cases, the adult gut microbiome, at large. For example, genes coding for homogalacturonan- and rhamnogalacturonan-degrading enzymes – most likely belonging to *Cephaloticoccus* sp. – are found in only a subset of turtle ant adult worker gut microbiomes (Fig 8; **S19 Fig**). This suggests that symbiont abilities to degrade pectins in adult turtle ants vary across host colonies, and plausibly among species. Such variability could be indicative of symbiont responses – plastic or evolved – to short- or long-term changes in host diet – quite possibly due to differing fractions and forms of pectins [21, 170]. While turtle ants are believed to consume wind-dispersed fungal spores and pollen [48,98,139], little is known about dietary variation across *Cephalotes* species. Nevertheless, given (i) the considerable differences in biochemical composition of pollen and fungal tissues across plant and fungal lineages [171–175], (ii) the wide Pan-American distribution of turtle ants, and (iii) the fact that these ants inhabit various habitats (*e.g.* rainforest, savannah, desert, mangrove; [139]), it is likely that the dietary fiber composition differs among turtle ant species. Pectinolytic enzyme-encoding genes of bacterial symbionts vary with the plant diet of folivorous beetles ([21, 170]), suggesting the possibility of adaptive change through expansion in symbiont-encoded digestive enzyme repertoires, or of relaxed selection enabling gene loss.

### Peculiarities of the adult-associated Campylobacterales symbionts

Beyond dietary fibers, some unique forms of sulfur metabolism are encoded in turtle ant microbiomes. Many turtle ant adult-associated symbionts encoded capacities to reduce sulfate into sulfide, a compound that is beneficial to some insects, by improving development, increasing resistance to desiccation [176, 177], and aiding in amino acid syntheses. Among the candidate sulfate-reducers are *Cephaloticoccus* sp., dominant symbionts in turtle ant midguts [111, 137]. Coincidentally, other common midgut-inhabiting turtle ant symbionts include Campylobacterales, which were found here to possess the complete genetic machinery to oxidize sulfide anaerobically. Although the oxic regimes in turtle ant gut compartments at the microscopic scale are unknown, the common – though not systematic – co-occurrence of *Cephaloticoccus* sp. and Campylobacterales in turtle ant midguts hints to a possible sulfur syntrophy between *Cephaloticoccus*sp. sulfate reducers and Campylobacterales sulfide oxidizers. Similar sulfur-oriented syntrophy may exist in other symbiotic systems, as evidenced from metagenomics in gutless annelid worms [178].

*Cephaloticoccus* sp. grow *in vitro* in environments of 12-20% O_2_ [113], suggesting that at least a portion of turtle ant midguts is enriched in O_2_. In contrast, although *Cephalotes*-derived Campylobacterales have not been successfully cultured yet, some of their genome-encoded metabolisms are anaerobic, including sulfide oxidation, potentially coupled to anaerobic dissimilatory nitrate reduction and the rTCA cycle. This indicates that Campylobacterales symbionts may occupy O_2_-depleted midgut micro-environments. Therefore, we hypothesize that, as demonstrated in termite and honeybee hindguts [14,51,179], inwardly diffusing O_2_ is consumed in turtle ant midguts and/or hindguts, allowing anaerobic (or microaerophilic) bacteria, like Campylobacterales, to thrive in the center of midgut luminal region. If the environment inside these ants’ guts allows these reactions to occur, sulfide oxidation could then be a source of energy for carbon autotrophs, and would commonly be coupled with valuable reductive pathways, as seen in related Campylobacterales from aquatic anoxic and microaerophilic, sulfur-rich environments [116, 180]. Based on further genome inferences for this taxon, nitrate reduction could be a key mechanism to generate NH_3_, used in the production of amino acids and some B-vitamins, in addition to providing energy when oxidized to nitrite [181]. Additionally, given its relatively limited dietary carbohydrate catabolic abilities (Fig 8; **S16 Fig**), Campylobacterales Bin17 may be able to fix CO_2_ as a primary source of carbon, in autotrophic, or at least mixotrophic, processes.

### Partner fidelity and partner choice in the *Cephalotes*-symbiont systems

In adult turtle ant guts, partner fidelity between symbionts and hosts has previously been demonstrated [61, 62]. Such preferential conservation of symbiont strains is not accidental, but rather facilitated by several anatomical, behavioral and developmental adaptations from the host acting as ecological forces shaping turtle ant gut symbiont communities [182]. First is faithful vertical transmission, with the symbiont community from a parent colony likely being carried by foundress queens [66], and within-colony spread via social transmission, involving a newly-emerged adult ingesting rectal fluids from a mature nestmate [26]. The timing of this anal-oral trophallactic event means that the core symbionts of mature adults are among the first microbes to colonize the bare ecosystem that constitutes the gut of a newly emerged adult. These first colonizers are then protected from environmental threats and outside microbes, by a fine filter that develops over the restrictive proventricular organ, separating the crop from the specialized symbiont-dominated midgut, ileum, and rectum [26]. Partner fidelity could be reinforced by physiochemical properties providing adequate ecological niches to core symbionts – a potential form of environmental filtering. It is also possible that microbes interact in competitive or mutualistic manners [183], such as the potential midgut syntrophy documented in the present study, contributing to shape gut symbiont communities by competitive exclusions or facilitations.

While high partner fidelity between adult turtle ants and their gut symbionts is indisputable, the mechanisms enabling integration of microbe in the larval gut microbiome are more obscure. Turtle ant larval guts are more exposed to their environment, as no anatomical filter prevents colonization by microbial intruders [26], and are therefore prone to occupation by free-living microbes. In that sense, and as discussed above, we usually found here that genes from Enterobacteriales, Lactobacillales and *Staphylococcus* sp. sampled from turtle ant larvae do not group into *Cephalotes*-specific, monophyletic groupings, but are most closely related to free-living bacteria (*e.g.* **S5 and S17 Figs**) – fitting with results from a recent 16S rRNA amplicon sequencing study [66]. Nevertheless, it is possible that certain species, or even strains, of bacteria are preferentially kept by larvae. For example, in the present study, we showed that the same species of *Enterococcus* (JR029-101/Bin3) has been sampled in larvae from two different turtle ant species: *C. varians* and *C. texanus* (**S8 Fig**), indicating possible regular acquisition of the same environmental bacteria by turtle ant larvae. Recurrence of the same unique sequences for Enterobacteriales and *Enterococcus* across turtle ant species [66], further supports the possibility of strong gut selectivity. These larvae, or the bacteria in larval guts, could thus select for bacteria encoding beneficial nutritional and/or digestive functions through as yet described partner choice mechanisms [184–186] (**S2 Text**).

### The digestive site conundrum

We found clear retention of genes encoding fiber-degrading enzymes from adult-associated gut symbionts across *Cephalotes* species, with some potential cases of PFCWDE lateral gene transfer among symbionts (**S2 Text; S21 and S22 Fig**). Such conservation of bacterial enzyme-encoding genes in symbiotic bacteria that are thought to live nowhere outside of turtle ants [58,62,157], suggests that these functions play important roles for the *Cephalotes*-symbiont system – regardless of whether they provide direct benefits to hosts or, instead, support microbes that may be beneficial through other means.

The presence of genes encoding PFCWDEs in *Cephalotes* adult-associated bacteria is somewhat surprising. This extends from the observation that this community of specialized bacterial symbionts is enriched in the midgut and hindgut of adult turtle ants [111,136,137], where fibers are unlikely to transit. Indeed, the sophisticated and narrow proventriculus of adult *Cephalotes* ants prevents microbes, and presumably macromolecular particles, from reaching the symbionts in the midgut and ileum [26,187,188], where most solid food digestion unfolds in other adult insects [189]. This raises the questions on the active sites of digestion, *i.e.* where do dietary fibers and symbiont-produced digestive enzymes meet and interact in a digestive process? It is predicted that enzymes encoded by symbionts sampled downstream the proventriculus, or the symbionts themselves, must exit the rectum and, through trophallaxes, travel to other digestive chambers where they exert action.

One possibility could be that adult-associated symbionts, or their digestive enzymes, like in fungus-growing ants [190], are transferred from adult hindguts to larval digestive tracts, where they would supplement digestive processes conducted by larva-associated symbionts. Supporting this, some of our results and the results from another study show that core adult symbiont species, from the Burkholderiales, Pseudomonadales and Xanthomonadales, can be sampled in larval guts (**S2, S3 and S6 Figs;** [66]). But other symbionts with presumed digestive capacities, including *Cephaloticoccus* sp. and Sphingobacteriales, appear to be rare in the larval gut habitat.

Another possible explanation for the retention of fiber degradative capabilities in adult turtle ant symbionts found in downstream gut compartments could be the transfer to, and activity within the foregut (*i.e.* infrabuccal pocket and/or crop) of other adult nestmates. Indeed, *Cephalotes* adult gut symbionts were shown to be transferred to the foregut of other adult nestmates via anal-oral trophallaxes [136], where recalcitrant substrates are directly available for pre-digestive enzymatic activity. Along these lines, turtle ant workers are known to partially break-down fungal and arthropod pieces for their larvae [50] and to regurgitate infrabuccal pellets packed with pollen grains from their foregut, which are occasionally fed to larvae and other adults [48]. These pellets could thus contain energy- and nutrient-rich food fragments, pre-digested by symbiont actions in the adult foregut. Gene-expression or metabolite analyses across gut compartments would undeniably improve our understanding of symbiont digestive mechanisms in turtle ants.

### Conclusion

We provide evidence that the taxonomically divergent gut microbiota of larval *vs.* adult turtle ants encode impressive metabolic repertoires, supporting key aspects of N, carbohydrate, and sulfur metabolism. A subset of these symbionts has the potential to generate energy through a series of processes, including sulfide oxidation, dietary fiber deconstruction and fermentation, and some may produce acetyl-CoA and organic acids of potential use to host energy budgets. Most members of adult and larval gut microbiomes synthesize amino acids, and several can also make B-vitamins, potentially missing from the ant’s diet. While functions of all but amino acid synthesis seem partitioned across various members of the adult microbiome – meaning that no one symbiont retained the majority of metabolic diversity represented in the full microbiome – in larvae, symbionts from the Enterobacteriales possess a very large number of potentially useful metabolic functions.

Taken together, our results suggest that turtle ant larval gut symbionts likely participate in the digestion of plant and fungal recalcitrant nutrients, raising the possibility that symbionts – including several acquired from the environment – aid the digestive caste function of larvae in turtle ant colonies. Specialized gut symbionts of adult turtle ants can also participate in colony-level digestion, though the site for such process is still unknown. While our genomic and metagenomic approach, supplemented by *in vitro* assays, enabled the exploration of symbiont metabolisms and their likely contribution to turtle ant nutrition, the present study raises a clear need for future work combining multi-omics and experimentation to decipher the complex network of symbiont functions uncovered here, and to elucidate host physiological and fitness responses to microbial metabolism.

## Supporting information

Supporting_information_list

## Acknowledgements

We thank Corrie Moreau, Scott Powell, and Jignasha Rana who collected specimens used in this study. We also thank Nicole Buleza who helped in metagenomic pathway screening. Finally, we thank Rachel Ehrlich who worked on the assembly of PacBio sequences.

## Funding

This study was supported by NSF grant DoB-1442144 awarded to JTW and JAR.

## Author contributions

**Conceptualization:** Benoît Béchade, Yi Hu, John T. Wertz and Jacob A. Russell.

**Data curation:** Benoît Béchade, Yi Hu and John T. Wertz.

**Formal analysis:** Benoît Béchade and Yi Hu.

**Funding acquisition:** John T. Wertz and Jacob A. Russell.

**Investigation:** Benoît Béchade, Yi Hu, Jon G. Sanders, Christian S. Cabuslay, Piotr Łukasik, Bethany R. Williams, Valerie J. Fiers, Richard Lu.

**Methodology:** Benoît Béchade, Yi Hu, Jon G. Sanders, Christian S. Cabuslay, Piotr Łukasik, John T. Wertz and Jacob A. Russell.

**Project administration:** Benoît Béchade and Jacob A. Russell.

**Resources:** Jon G. Sanders, John T. Wertz and Jacob A. Russell.

**Software:** Benoît Béchade, Yi Hu, Jon G. Sanders and Piotr Łukasik.

**Supervision:** Benoît Béchade, John T. Wertz and Jacob A. Russell.

**Validation:** Benoît Béchade, Yi Hu and Jacob A. Russell.

**Visualization:** Benoît Béchade.

**Writing – Original draft preparation:** Benoît Béchade and Jacob A. Russell.

**Writing – Review and editing:** Benoît Béchade, Yi Hu, Jon G. Sanders, Piotr Łukasik, John T. Wertz and Jacob A. Russell.

**Data availability** Assembled genomes and metagenomes new to this study are available in IMG/M-ER under the following GOLD Sequencing Project IDs: Gp0095984, Gp0055707, Gp0224581, Gp0224584. Raw sequence data are available upon request from the authors. Supplementary data can be found in our data repository at: 10.6084/m9.figshare.15173730.

**Conflict of interest** The authors declare that they have no conflict of interest.

## Notes

### Competing Interest Statement

The authors have declared no competing interest.

