## Supporting_information_list for "In their sister’s footsteps: Taxonomic divergence obscures substantial functional overlap among the metabolically diverse symbiotic gut communities of adult and larval turtle ants"

**List of supporting information**

**S1 Fig. Experimental design.** Larval and adult gut metagenomes were generated, and 18 individual Metagenome-Assembled Genomes (MAGs) were produced from an adult and a larval metagenome originating from the same colony (PL010). Additionally, we cultured and isolated symbionts and sequenced the genome of 14 of these cultured Isolate Genomes (IGs). These IGs were isolated from *Cephalotes varians* and *C. rohweri* adults and two were from *C. texanus* larvae. Assembled (meta)genomes were annotated in IMG/M-ER and screened for metabolic pathway completeness, followed by rigorous curation and verification of the results (see **S4 Fig**). Finally, some of the isolates were tested for their metabolic potential in *in vitro* assays.

**S2 Fig. Taxonomic overview of the larval PL005 metagenome.** Assembled scaffolds from the metagenome were taxonomically annotated using the USEARCH-IMG/M-ER pipeline [79–81]. Guanine (G) and cytosine (C) content, which varies among core bacterial genomes, is displayed on the x-axis. Depth of sequencing coverage is a proxy for the relative abundance of core symbionts, shown on the y-axis. When compared to dominant taxa in our metagenomes with results from Hu et al. [66], who inferred the core gut bacterial composition of adults and larvae using 16S rRNA amplicon sequencing, spanning several colonies of *Cephalotes varians* and other *Cephalotes* species, the bacterial composition in this larval metagenome was found to be unusual. Therefore, we treated this metagenome as not representative of the larval stage in *C. varians*, and focused our analyses on PL010 metagenomes.

**S3 Fig. Relative abundance of bacterial 97%-OTUs found in the larval gut of multiple *Cephalotes* species.** The 16S rRNA amplicon sequencing data is from Hu et al. [66]. The three most common bacterial orders sampled in larvae from twelve different turtle ant species are Rhizobiales, Lactobacillales and Enterobacteriales. Although rare, some OTUs from conserved adult-associated symbionts (*e.g.* Burkholderiales, Xanthomonadales, *Cephaloticoccus* sp. (Opitutales) and Pseudomonadales) can be sampled in larval guts.

**S4 Fig. Genomic and** **metagenomic** **exploration workflow.** Focal metabolic pathways were first reviewed in KEGG and Metacyc websites, and in the literature to produce a list of KEGG KO and EC identifiers. PL010 metagenomes and all IGs from *Cephalotes* guts, previously uploaded in IMG/M-ER, were screened using these KO and EC lists. The resulting list was curated through different steps, to remove false positives and add false negatives. Additional verifications using BLASTPs were performed for nitrogen recycling and carbohydrate catabolisms, and genes for herbivore and fungivore digestive enzymes (PFCWDEs) were retrieved from multiple *Cephalotes* metagenomes followed by phylogenetic analyses. Blue ellipses: actions performed. Yellow squares: inputs and outputs. Orange ellipses: step targeting false positives and/or false negatives. Squares with white background: examples of inputs and outputs from specific pathway screened in our study, identified by font colors, with gene counts retained at each step of the workflow. The ellipses with white background depict number of genes identified as false positives removed or false negatives added.

**S5 Fig. Pectate lyase and larval polygalacturonase protein phylogenetic trees showing examples of annotations well supported, mildly supported, and not supported.** Rooted phylogenetic tree of **(A)** pectate lyase and **(B)** polygalacturonase amino acid sequences from *Cephalotes*-associated symbionts. Sequences were obtained from a screening in IMG/M-ER against seven *Cephalotes* metagenomes. These sequences were used as queries for an IMG/M-ER BLAST against the 20 *Cephalotes* metagenomes sequenced and the 14 *Cephalotes* IGs and for a BLASTP against custom NCBI databases. A maximum likelihood method was used with 999 bootstrap iterations to build the trees. All bootstrap values are shown. Colors of host *Cephalotes* species (outer strip) are from [61]. **(A)** The annotation as pectate lyase is only well supported for one gene from *Staphylococcus* sp. JDR108L-110-1 IG, falling into a monophyletic group along with other pectate lyase-encoding genes from non-*Cephalotes* associated Bacillales. **(B)** The annotation as polygalacturonase is mildly supported for these three Enterobacteriales genes given that they are closely related to genes with unprecise annotation (“glycoside hydrolase”).

**S6 Fig. Taxonomic description of MAGs in the *Cephalotes varians* PL010 larval gut metagenome**. Assembled scaffolds from the PL010 larval metagenome were taxonomically annotated using the USEARCH-IMG/M-ER pipeline [79–81]. See **S2 Fig** for a legend on GC-blobplots. The first GC-blobplot is a more detailed version of **Fig 2A**, with, additionally, scaffolds < 500 bp, and annotations of less represented taxonomic orders. The following pairs of GC-blobplots illustrate the possible classification of certain MAGs. Enterobacteriales and Lactobacillales MAGs could be classified at a lower taxonomic level, but it was not the case for Rhizobiales MAGs.

**S7 Fig. Taxonomic description of MAGs in the *Cephalotes varians* PL010 adult worker gut metagenome**. Assembled scaffolds from the PL010 adult metagenome were taxonomically annotated using the USEARCH-IMG/M-ER pipeline [79–81]. See **S2 Fig** for a legend on GC-blobplots. The first GC-blobplot is a more detailed version of **Fig 2A**, with, additionally, scaffolds < 500 bp, and annotations to less represented taxonomic orders. The following pairs of GC-blobplots illustrate the possible classification of certain MAGs. While Burkholderiales scaffolds are easy to assign to bacterial families, classifying Rhizobiales and Xanthomonadales MAGs is more difficult.

**S8 Fig. Pairwise Average Nucleotide Identity (ANI) calculated among *Cephalotes*-associated symbiont genomes.** Assembled scaffolds from *C. varians* PL010 metagenomes were assigned to MAGs (with the mention “Bin” or “Merg”) using the Anvi’o-CONCOCT pipeline. Additional IGs from *C. varians*, *C. texanus* and *C. rohweri* were included. The bacterial phylogenetic trees were inferred based on seven bacterial marker gene sequences using a maximum likelihood method and 999 bootstrap iterations. Bootstrap values <100 are shown on the tree. OrthoANI values were calculated as described in Yoon et al. (2017) [85]. Higher orthoANIs indicate higher levels of genomic similarity among symbionts.

**S9 Fig. B-vitamin biosynthesis pathways encoded in *Cephalotes*-associated genomes.** MAGs (with the mention “Bin” or “Merg”) and IGs next to an arrow indicates that they possess the genes for the focal step. Dashed squares correspond to pathways half encoded to almost complete. The color of turtle ant adults and larvae correspond to the species symbionts were sampled from, as in **S8 Fig**. Note that many *Cephalotes*-associated symbionts have retained genes for B-vitamin biosynthesis.

**S10 Fig. Organic acid metabolism pathways encoded in *Cephalotes*-associated genomes.** MAGs (with the mention “Bin” or “Merg”) and IGs next to an arrow indicates that they possess the genes for the focal step. Blue arrows are for steps that can be fermentative, taking place in anaerobic environments. The color of turtle ant adults and larvae correspond to the species symbionts were sampled from, as in **S8 Fig**. Note that many *Cephalotes*-associated symbionts have retained genes for organic acid metabolism, and many can ferment pyruvate to lactate and acetate. Symbionts from larvae (plus Rhizobiales JR021-5/Bin10) can additionally ferment citrate and ferment pyruvate to produce formate.

**S11 Fig. Gene-level N-recycling pathways in *Cephalotes*-associated symbiont genomes.** Presence of genes encoding steps of N-recycling in each symbiont genome is indicated in squares. Black squares without any gene name are for genes not found in focal genomes. Gene names with a white font and without background color correspond to genes not found in our screenings, but likely to be present because they were found in genomes of highly related *Cephalotes*-associated symbionts (see **S8 Fig**). For a visual description of enzymatic step numbers, see **Fig 6B**. The bacterial phylogenetic trees were inferred based on seven bacterial marker gene sequences using a maximum likelihood method and 999 bootstrap iterations. Bootstrap values <100 are shown on the tree.

**S12 Fig. Clusters of genes coding for steps of N-recycling pathways from larval and adult gut symbionts.** Step numbers and gene names correspond to the ones shown in **Fig 6B**.

**S13 Fig. Gene cluster from the larval gut metagenome including genes to import pectate and degrade galacturonate.** The scaffold is from the Enterobacteriales order, symbiont *Klebsiella* sp. MergB4 MAG. For a visual description of gene names and their encoded enzymes, see **Fig 7A**.

**S14 Fig. Carbohydrate Active Enzyme (CAZy) screening results for *Cephalotes varians* PL010 larval and adult worker gut metagenomes.** The metagenomes were screened against the HMMER, DIAMOND and Hotpep databases for CAZy families (CAZymes) using the dbCAN2 metaserver (v7, January 2019) [101]. Catabolic CAZymes were categorized based on the type of substrates the enzymes can degrade. Degradations are as follow: “St”: Starch, “HC”: Hemicellulose, “Ce”: Cellulose, “Pe”: Pectin, “Pf”: Polyfructans, “Gu”: Gums, “Ch”: Chitin, “Li”: Lignin.

**S15 Fig. Pathways for carbohydrate catabolism screened in *Cephalotes varians* PL010 larval and adult metagenomes.** Results at the symbiont genome-level can be seen in **S16 Fig**. Pink arrows are for steps encoded in the larval metagenome, and blue arrows are for steps encoded in the adult metagenome. Note that homogalacturonan catabolic pathway can be found in **Fig 7A**. “Es”: Esterification, “C1”: Cut on a polymer backbone, “C2”: Cut on a disaccharide, “De”: Debranching of a polymer side chain, “Im”: Import of disaccharides or monosaccharides in a bacterial cytoplasm, “P”: Phosphorylation of a carbohydrate, “C3”: Cut of a carbohydrate ring and transformation into an assimilable compound.

**S16 Fig.** **Carbohydrate catabolic profile of *Cephalotes*-associated symbionts in larval-associated symbionts.** Heatmap depicting carbohydrate catabolisms encoded in genomes of distinct *Cephalotes*-associated bacterial gut symbionts. Proportions of step categories encoded in symbiont genomes (columns) to perform various metabolisms (rows) is given by the shade within the squares of the heatmap, the darker the more complete the metabolic step is. Symbiont genomes were screened in IMG/M-ER for the presence of key enzyme- and transporter-encoding genes. Bacterial genes from the *C. varians* PL010 metagenomes that were not binned to MAGs (with the mention “Bin” or “Merg”) but having a predicted relevant function are pooled in the “Other bacteria in larval gut” and “Other bacteria from adult worker gut” columns, respectively. Additional IGs from *C. varians*, *C. texanus* and *C. rohweri* were included. The bacterial phylogenetic trees were inferred based on seven bacterial marker gene sequences using a maximum likelihood (ML) method and 999 bootstrap iterations. Bootstrap values <100 are shown on the tree. Hatched squares signal when step categories were retained as complete or incomplete but with contradicting evidence. See **S15 Fig** for details on abbreviations of enzyme/transporter types.

**S17 Fig. Protein phylogenetic trees for conserved genes encoding fiber degrading-enzymes in the Xanthomonadales adult associates.** Rooted phylogenetic tree of **(A)** beta-xylosidase, **(B)** chitinase and **(C)** alpha-amylase amino acid sequences from *Cephalotes*-associated symbionts. Sequences were obtained from a screening in IMG/M-ER against seven *Cephalotes* metagenomes. These sequences were used as queries for an IMG/M-ER BLAST against the 20 *Cephalotes* metagenomes sequenced and the 14 *Cephalotes* IGs and for a BLASTP against custom NCBI databases. A maximum likelihood method was used with 999 bootstrap iterations. Bootstrap values are shown on the tree. Colors of host *Cephalotes* species (outer circle) are from [61].

**S18 Fig. Results of *in vitro* metabolic assays for utilization of cellulose, chitin and starch derivatives.** The assays were conducted on cultured bacteria isolated from *Cephalotes varians* and *C. rohweri* adult guts and bacteria from *C. varians* and *C. texanus* larval guts. Each assay was run in triplicates. The phylogenetic tree was built from 16S rRNA sequences extracted from individual isolates using a ML method and 999 bootstrap iterations. Bootstrap values <100 are shown on the tree. Full results of *in vitro* metabolic assays can be found in **S8 Table**.

**S19 Fig. Protein phylogenetic trees for pectinolytic enzymes in the *Cephaloticoccus* adult associates.** Rooted phylogenetic tree of **(A)** rhamnogalacturonan endolyase, **(B)** rhamnogalacturonyl hydrolase and **(C)** polygalacturonase (adults only) amino acid sequences from *Cephalotes*-associated symbionts. Trees were built similarly as in **S16 Fig**. Grey and coral horizontal bar graphs show the GC content of scaffolds from *Cephalotes-*associated symbionts (coral color) or of entire genomes from NCBI (grey color). Orange horizontal bar graphs show the read depths of scaffolds from *Cephalotes-*associated symbionts. Note that genes coding for these two enzymes from metagenomes of many *Cephalotes* species group into monophyletic branches with most scaffolds having similar GC content (0.56 on average) and occasionally with high read depth. Genes with such high read depth fit the characteristics of *Cephaloticoccus* sp.

**S20 Fig. Origins, and functional and taxonomic assignments of cellulase-encoding genes from *Cephalotes*-associated symbionts. (A)** Phylogenetic tree of cellulase amino acid sequences from *Cephalotes*-associated symbionts. The tree was built similarly as in **S16 Fig** and rooted using a *Gloeocapsa* sp. (Cyanobacteria) sequence ([AFZ33259.1](https://www.ncbi.nlm.nih.gov/protein/AFZ33259.1?report=genbank&log$=protalign&blast_rank=1&RID=BSBFGKW0014)). Note that Hymenopteran, Opitutales and Rhizobiales sequences were removed from the tree due to a lack of support for their cellulase annotation. **(B)** Taxonomically annotated bacterial scaffolds with a cellulase-encoding gene in *Cephalotes* adult gut metagenomes. The two scaffolds at the bottom are the longest assembled scaffolds from *Cephalotes* gut metagenomes containing a cellulase-encoding gene. The colors of the genes correspond to their bacterial taxonomic annotation at the order level inferred with online NCBI BLASTNs.

Taxonomic annotation of single genes shows evidence of a cellulose-synthesis operon horizontally transferred from a Pseudomonadales bacterium to a Xanthomonadales *Cephalotes*-associated bacterial symbiont. The scaffold on the top shows the *wssA-J* cellulose-synthetase operon from the biofilm-producing *Pseudomonas fluorescens* SBW25 detailed in [151]. Note that a cellulase-encoding gene is naturally present on such cellulose-synthetase operon.

**S21 Fig. Gene-level taxonomy of symbiont genes for herbivore and fungivore digestive enzymes (PFCWDEs) across *Cephalotes* species.** This figure is similar to **Fig 8**, but the bacterial taxonomic classification was performed at the gene level here, instead of scaffold level. The pie charts show the proportion of genes from bacteria identified at the taxonomic order (colors) to code for specific enzymes (columns) in given *Cephalotes* hosts (rows). The *Cephalotes* phylogeny is adapted from [102]. Contrasting with **Fig 8**, the classification is less ambiguous, indicating potential horizontal gene transfers having occurred among *Cephalotes*-­associated gut bacteria.

**S22 Fig. Protein phylogenetic trees for putatively horizontally exchanged genes encoding fiber-degrading enzymes in the *Cephaloticoccus* adult associates.** Rooted phylogenetic tree of **(A)** alpha-arabinofuranosidase and **(B)** beta-fructosidase amino acid sequences from *Cephalotes*-associated symbionts. Trees were built similarly as in **S16 Fig**. Based on these reconstructed phylogenies, *Cephaloticoccus* sp. may have received an alpha-arabinofuranosidase-encoding gene from Alphaproteobacteria, and beta-fructosidase-encoding genes may have been transferred from Flavobacteriales or Sphingobacteriales to Xanthomonadales and *Cephaloticoccus* sp.

**S1 Table. Assembly statistics of metagenomic data.** Rows in bold indicate metagenomes new to this study.

**S2 Table. Assembly statistics of Cultured Isolate Genomes (IGs) and cultivation conditions of cultured bacteria.** Rows in bold indicate IGs new to this study.

**S3 Table. Summary of strain-level binning for gut metagenomes from *Cephalotes varians* workers and larvae in colony PL010.** Genes from *C. varians* PL010 larval and adult worker gut metagenomes were assigned to Metagenome-Assembled Genomes (MAGs, with the mention “Bin” or “Merg”) as described in [58] using the Anvi’o-CONCOCT pipeline. “PL010-L” corresponds to the larval gut metagenome, and “PL010-W” corresponds to the adult worker gut metagenome. Rows in bold indicate MAGs new to this study.

**S4 Table. List of 48 key plant and fungal cell wall digestive enzymes gathered from the literature** **and our previous carbohydrate catabolism screening.** The Function IDs were used in IMG/M-ER to screen adult gut metagenomes from 6 *Cephalotes* species and the larval PL010 *C.* *varians* gut metagenome.

**S5 Table. Binning results and taxonomic classification of metagenome-derived assembled scaffolds into MAGs.** Genes from *Cephalotes varians* PL010 larval and adult worker gut metagenomes were assigned to MAGs (with the mention “Bin” or “Merg”) as described in [58] using the Anvi’o-CONCOCT pipeline. “PL010-L” corresponds to the larval gut metagenome, and “PL010-W” corresponds to the adult worker gut metagenome.

**S6 Table. Bacterial gene counts per KEGG pathway in PL010 *Cephalotes varians* adult and larval gut metagenomes and statistical comparisons.** KEGG pathways are sorted from the most (top) to the least (bottom) enriched in both larval and adult metagenomes. Bacterial gene counts per KEGG pathway were normalized by the total count of KO-encoding bacterial genes. Corresponding *Z*-normalized log odds ratios between adult and larval metagenomes were calculated. “PL010-L” corresponds to the larval gut metagenome, and “PL010-W” corresponds to the adult worker gut metagenome. Significantly enriched KEGG pathways are indicated in bold and correspond to KEGG pathways with Benjamini-Hochberg critical value < 0.05.

**S7 Table. List of KEGG Orthologies (KOs) found in significantly enriched pathways in the *Cephalotes varians* PL010 larval (left table) and adult (right table) metagenomes when compared to one another.** For each KO (rows), the bacterial gene count encoding it, the bacterial gene proportion in the metagenome, the presence/absence (1/0) of this KO in significantly enriched pathways, and the number of significantly enriched pathways in the metagenome to have this KO are indicated in the different columns.

**S8 Table. *in vitro* metabolic assay results.** Bacteria isolates were by 16S rRNA gene sequencing. Taxonomic classifications for these bacteria were achieved by submitting these sequences to the Classifier program in the Ribosomal Database Project website ([rdp.cme.msu.edu](http://rdp.cme.msu.edu/)). All the assays were performed on GN3 microplates (Biolog, Hayward, CA, USA) except for pectin depolymerization assay where the plate assay method from Engel et al. (2012) [15] was followed, and for urease assay where Rapid Urea Broth (Becton Dickinson, Sparks, MD, USA) was used as described in [58]. Each assay was run in triplicates. “NT”: Not Tested.

**S9 Table. CAZy screening results for *Cephalotes varians* PL010 larval and adult worker gut metagenomes.** The metagenomes were screened against the HMMER, DIAMOND and Hotpep databases for Carbohydrate Active Enzyme families (CAZymes) using the dbCAN2 metaserver (v7, January 2019) [101]. Catabolic CAZymes were categorized based on the type of substrates the enzymes can degrade. “PL010-L” corresponds to the larval gut metagenome, and “PL010-W” corresponds to the adult worker gut metagenome. Degradations are as follow: “St”: Starch, “HC”: Hemicellulose, “Ce”: Cellulose, “Pe”: Pectin, “Pf”: Polyfructans, “Gu”: Gums, “Ch”: Chitin, “Li”: Lignin.

**S1 Text. Supplementary materials and methods.**

**S2 Text. Supplementary results and discussion.**
