## Supplementary material for "In their sister’s footsteps: Taxonomic divergence obscures substantial functional overlap among the metabolically diverse symbiotic gut communities of adult and larval turtle ants": S1_Fig

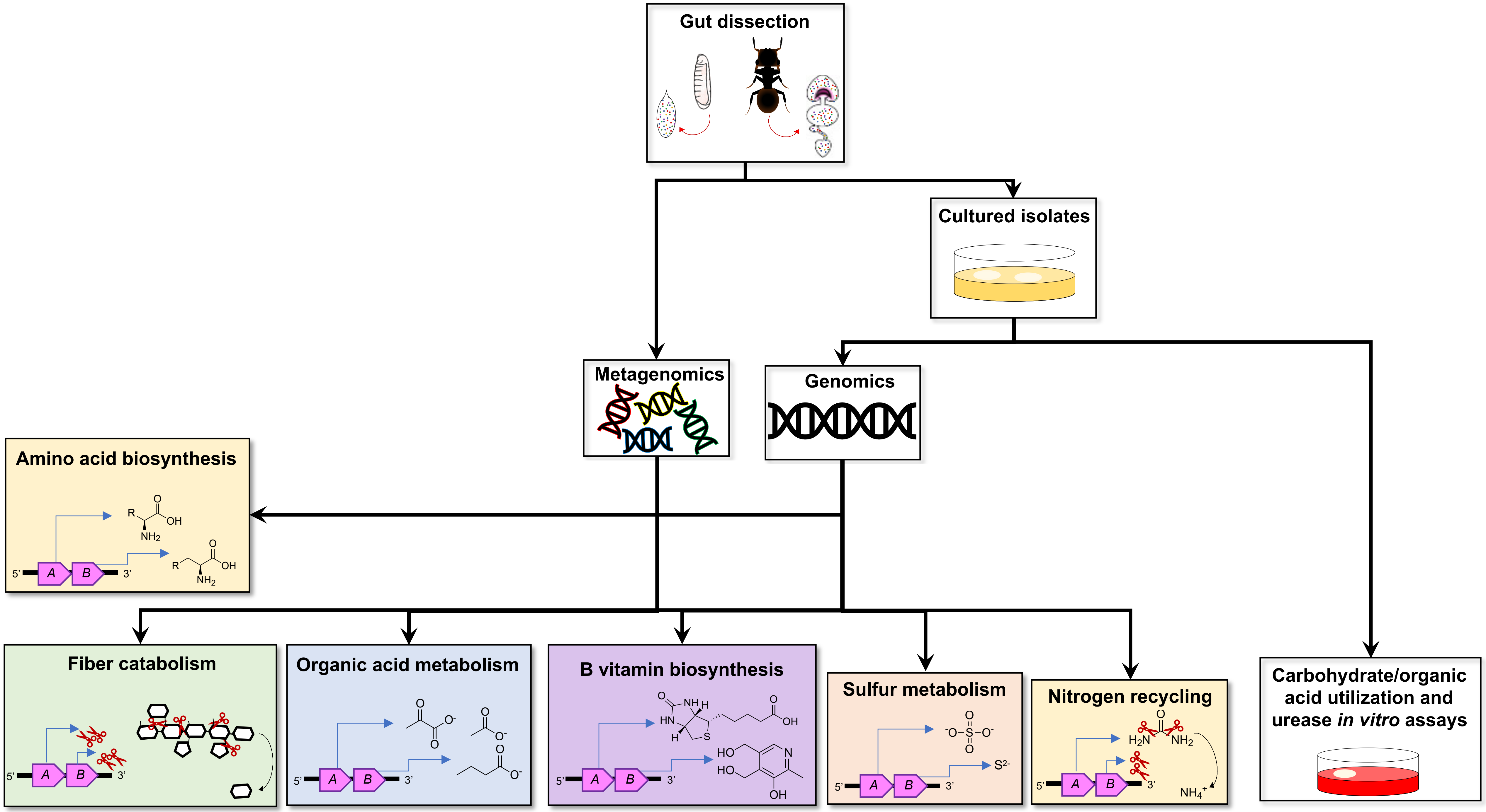

Gut dissection
Cultured isolates
Metagenomics
Genomics
Amino acid biosynthesis
Fiber catabolism
Organic acid metabolism
B vitamin biosynthesis
Sulfur metabolism
Nitrogen recycling
Carbohydrate/organic acid utilization and urease *in vitro* assays

**S1 Fig. Experimental design.** Larval and adult gut metagenomes were generated, and 18 individual Metagenome-Assembled Genomes (MAGs) were produced from an adult and a larval metagenome originating from the same colony (PL010). Additionally, we cultured and isolated symbionts and sequenced the genome of 14 of these cultured Isolate Genomes (IGs). These IGs were isolated from *Cephalotes varians* and *C. rohweri* adults and two were from *C. texanus* larvae. Assembled (meta)genomes were annotated in IMG/M-ER and screened for metabolic pathway completeness, followed by rigorous curation and verification of the results (see **S4 Fig**). Finally, some of the isolates were tested for their metabolic potential in *in vitro* assays.
