## Supplementary material for "In their sister’s footsteps: Taxonomic divergence obscures substantial functional overlap among the metabolically diverse symbiotic gut communities of adult and larval turtle ants": S2_Fig

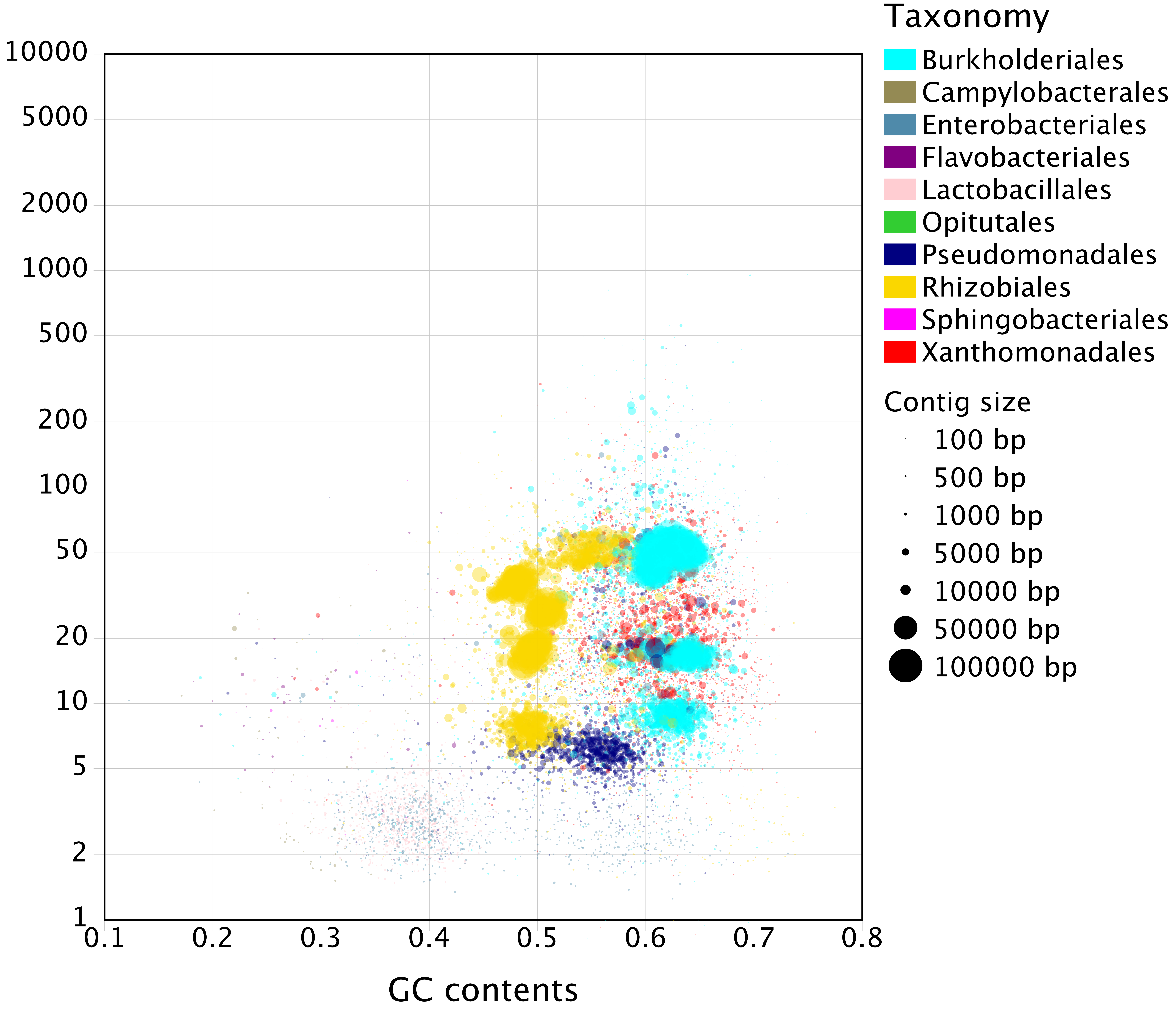

Taxonomy
Burkholderiales
Campylobacterales
Enterobacteriales
Flavobacteriales
Lactobacillales
Opitutales
Pseudomonadales
Rhizobiales
Sphingobacteriales
Xanthomonadales
Contig size
100 bp
500 bp
1000 bp
5000 bp
10000 bp
50000 bp
100000 bp
10000
5000
2000
1000
500
200
100
50
20
10
5
2
1
0.1
0.2
0.3
0.4
0.5
0.6
0.7
0.8
GC contents

**S2 Fig. Taxonomic overview of the larval PL005 metagenome.** Assembled scaffolds from the metagenome were taxonomically annotated using the USEARCH-IMG/M-ER pipeline [79–81]. Guanine (G) and cytosine (C) content, which varies among core bacterial genomes, is displayed on the x-axis. Depth of sequencing coverage is a proxy for the relative abundance of core symbionts, shown on the y-axis. When compared to dominant taxa in our metagenomes with results from Hu et al. [66], who inferred the core gut bacterial composition of adults and larvae using 16S rRNA amplicon sequencing, spanning several colonies of *Cephalotes varians* and other *Cephalotes* species, the bacterial composition in this larval metagenome was found to be unusual. Therefore, we treated this metagenome as not representative of the larval stage in *C. varians*, and focused our analyses on PL010 metagenomes.
