## Supplementary material for "In their sister’s footsteps: Taxonomic divergence obscures substantial functional overlap among the metabolically diverse symbiotic gut communities of adult and larval turtle ants": S3_Fig

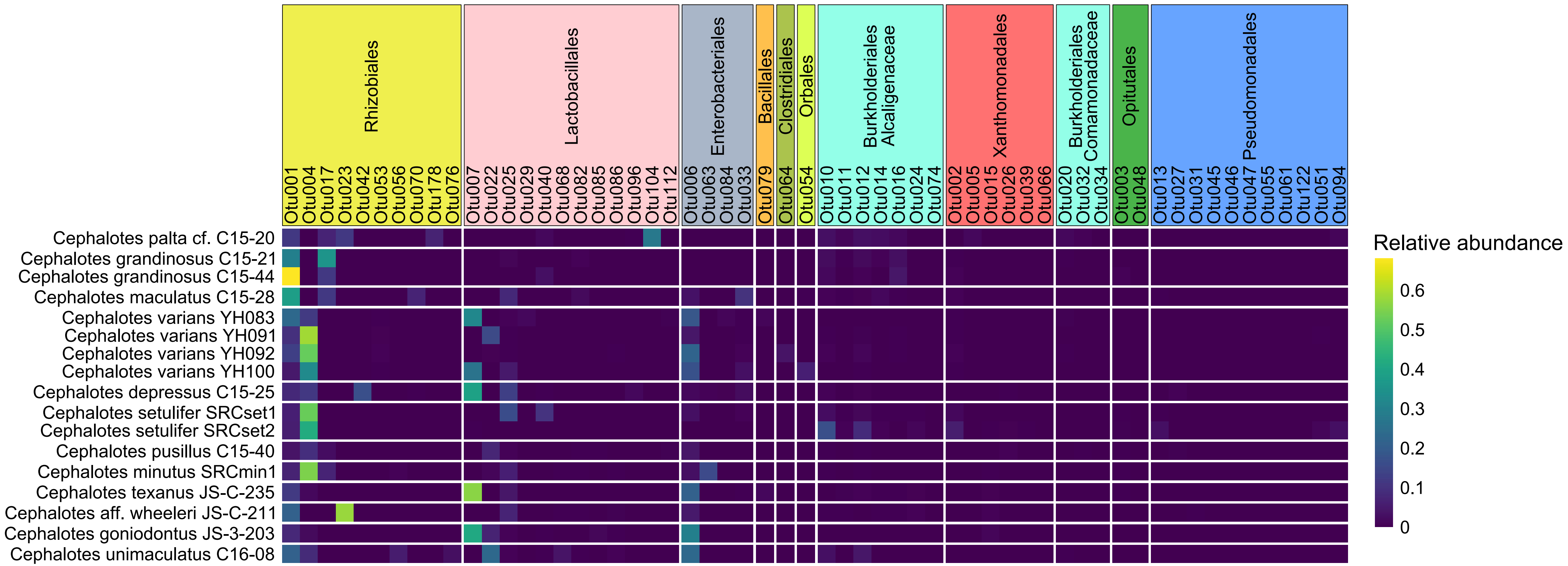
Rhizobiales
Lactobacillales
Enterobacteriales
Bacillales
Clostridiales
Orbales
Burkholderiales Alcaligenaceae
Xanthomonadales
Burkholderiales Comamonadaceae
Opitutales
Pseudomonadales
Otu001
Otu004
Otu017
Otu023
Otu042
Otu053
Otu056
Otu070
Otu178
Otu076
Otu007
Otu022
Otu025
Otu029
Otu040
Otu068
Otu082
Otu085
Otu086
Otu096
Otu104
Otu112
Otu006
Otu063
Otu084
Otu033
Otu079
Otu064
Otu054
Otu010
Otu011
Otu012
Otu014
Otu016
Otu024
Otu074
Otu002
Otu005
Otu015
Otu026
Otu039
Otu066
Otu020
Otu032
Otu034
Otu003
Otu048
Otu013
Otu027
Otu031
Otu045
Otu046
Otu047
Otu055
Otu061
Otu122
Otu051
Otu094
Cephalotes palta cf. C15-20
Cephalotes grandinosus C15-21
Cephalotes grandinosus C15-44
Cephalotes maculatus C15-28
Cephalotes varians YH083
Cephalotes varians YH091
Cephalotes varians YH092
Cephalotes varians YH100
Cephalotes depressus C15-25
Cephalotes setulifer SRCset1
Cephalotes setulifer SRCset2
Cephalotes pusillus C15-40
Cephalotes minutus SRCmin1
Cephalotes texanus JS-C-235
Cephalotes aff. wheeleri JS-C-211
Cephalotes goniodontus JS-3-203
Cephalotes unimaculatus C16-08
Relative abundance
0.6
0.5
0.4
0.3
0.2
0.1
0

**S3 Fig. Relative abundance of bacterial 97%-OTUs found in the larval gut of multiple *Cephalotes* species.** The 16S rRNA amplicon sequencing data is from Hu et al. [66]. The three most common bacterial orders sampled in larvae from twelve different turtle ant species are Rhizobiales, Lactobacillales and Enterobacteriales. Although rare, some OTUs from conserved adult-associated symbionts (*e.g.* Burkholderiales, Xanthomonadales, *Cephaloticoccus* sp. (Opitutales) and Pseudomonadales) can be sampled in larval guts.
