## Supplementary material for "In their sister’s footsteps: Taxonomic divergence obscures substantial functional overlap among the metabolically diverse symbiotic gut communities of adult and larval turtle ants": S4_Fig

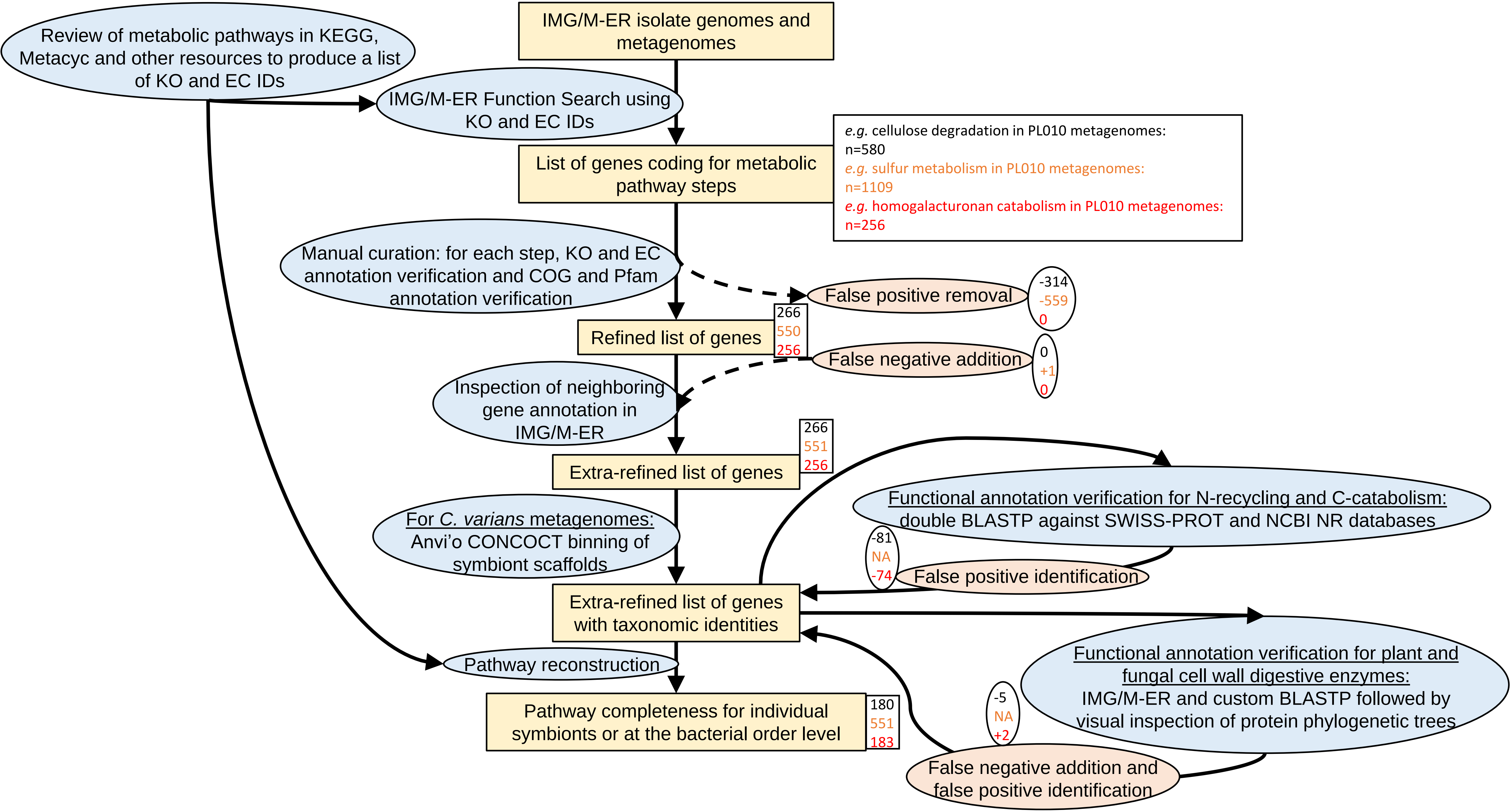

Review of metabolic pathways in KEGG, Metacyc and other resources to produce a list of KO and EC IDs
IMG/M-ER isolate genomes and metagenomes
IMG/M-ER Function Search using KO and EC IDs
List of genes coding for metabolic pathway steps
e.g. cellulose degradation in PL010 metagenomes: n=580
e.g. sulfur metabolism in PL010 metagenomes: n=1109
e.g. homogalacturonan catabolism in PL010 metagenomes: n=256
Manual curation: for each step, KO and EC annotation verification and COG and Pfam annotation verification
False positive removal
-314
-559
0
Refined list of genes
266
550
256
False negative addition
0
+1
0
Inspection of neighboring gene annotation in IMG/M-ER
Extra-refined list of genes
266
551
256
Functional annotation verification for N-recycling and C-catabolism: double BLASTP against SWISS-PROT and NCBI NR databases
For C. varians metagenomes: Anvi'o CONCOCT binning of symbiont scaffolds
-81
NA
-74
False positive identification
Extra-refined list of genes with taxonomic identities
Functional annotation verification for plant and fungal cell wall digestive enzymes: IMG/M-ER and custom BLASTP followed by visual inspection of protein phylogenetic trees
Pathway reconstruction
Pathway completeness for individual symbionts or at the bacterial order level
180
551
183
-5
NA
+2
False negative addition and false positive identification

**S4 Fig. Genomic and metagenomic exploration workflow.** Focal metabolic pathways were first reviewed in KEGG and Metacyc websites, and in the literature to produce a list of KEGG KO and EC identifiers. PL010 metagenomes and all IGs from *Cephalotes* guts, previously uploaded in IMG/M-ER, were screened using these KO and EC lists. The resulting list was curated through different steps, to remove false positives and add false negatives. Additional verifications using BLASTPs were performed for nitrogen recycling and carbohydrate catabolisms, and genes for herbivore and fungivore digestive enzymes (PFCWDEs) were retrieved from multiple *Cephalotes* metagenomes followed by phylogenetic analyses. Blue ellipses: actions performed. Yellow squares: inputs and outputs. Orange ellipses: step targeting false positives and/or false negatives. Squares with white background: examples of inputs and outputs from specific pathway screened in our study, identified by font colors, with gene counts retained at each step of the workflow. The ellipses with white background depict number of genes identified as false positives removed or false negatives added.
