## Supplementary material for "In their sister’s footsteps: Taxonomic divergence obscures substantial functional overlap among the metabolically diverse symbiotic gut communities of adult and larval turtle ants": S5_Fig

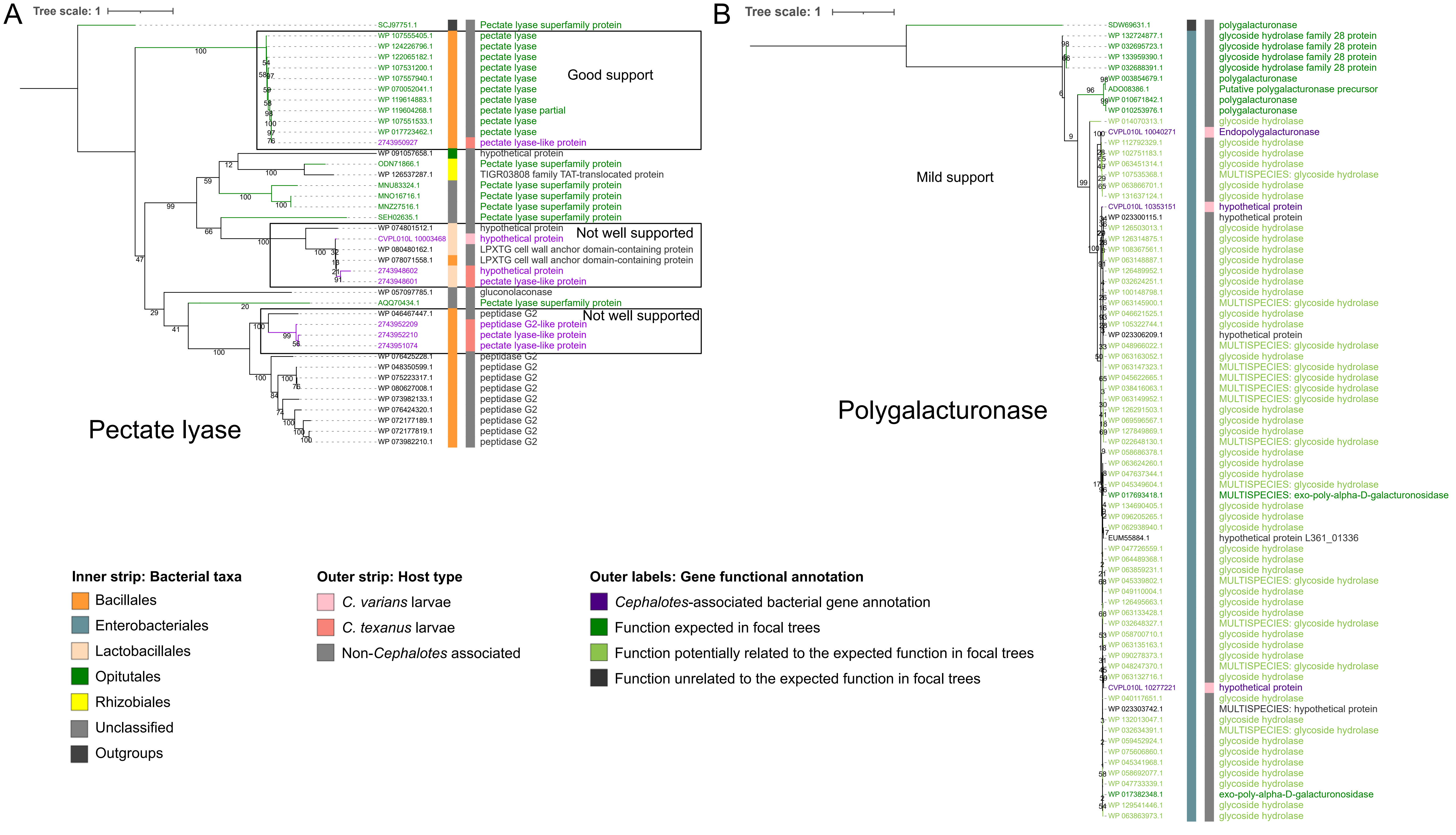
A
Tree scale: 1
SCJ97751.1 Pectate lyase superfamily protein
WP 107555405.1 pectate lyase
WP 124226796.1 pectate lyase
WP 122065182.1 pectate lyase
WP 107531200.1 pectate lyase
WP 107557940.1 pectate lyase
WP 070052041.1 pectate lyase
WP 119614883.1 pectate lyase
WP 119604268.1 pectate lyase partial
WP 107551533.1 pectate lyase
WP 017723462.1 pectate lyase
2743950927 pectate lyase-like protein
Good support
WP 091057658.1 hypothetical protein
ODN71866.1 Pectate lyase superfamily protein
WP 126537287.1 TIGR03808 family TAT-translocated protein
MNU83324.1 Pectate lyase superfamily protein
MNO16716.1 Pectate lyase superfamily protein
MNZ27516.1 Pectate lyase superfamily protein
SEH02635.1 Pectate lyase superfamily protein
WP 074801512.1 hypothetical protein
CVPL010L 10003468 hypothetical protein
WP 080480162.1 LPXTG cell wall anchor domain-containing protein
WP 078071558.1 LPXTG cell wall anchor domain-containing protein
2743948602 hypothetical protein
2743948601 pectate lyase-like protein
Not well supported
WP 057097785.1 gluconolaconase
AQQ70434.1 Pectate lyase superfamily protein
WP 046467447.1 peptidase G2
2743952209 peptidase G2-like protein
2743952210 pectate lyase-like protein
2743951074 pectate lyase-like protein
Not well supported
WP 076425228.1 peptidase G2
WP 048350599.1 peptidase G2
WP 075223317.1 peptidase G2
WP 080627008.1 peptidase G2
WP 073982133.1 peptidase G2
WP 076424320.1 peptidase G2
WP 072177189.1 peptidase G2
WP 072177819.1 peptidase G2
WP 073982210.1 peptidase G2
Pectate lyase
B
Tree scale: 1
SDW69631.1 polygalacturonase
WP 132724877.1 glycoside hydrolase family 28 protein
WP 032695723.1 glycoside hydrolase family 28 protein
WP 133959390.1 glycoside hydrolase family 28 protein
WP 032688391.1 glycoside hydrolase family 28 protein
WP 003854679.1 polygalacturonase
ADO08386.1 Putative polygalacturonase precursor
WP 010671842.1 polygalacturonase
WP 010253976.1 polygalacturonase
WP 014070313.1 glycoside hydrolase
CVPL010L 10040271 Endopolygalacturonase
WP 112792329.1 glycoside hydrolase
WP 102751183.1 glycoside hydrolase
WP 063451314.1 glycoside hydrolase
WP 107535368.1 MULTISPECIES: glycoside hydrolase
WP 063866701.1 glycoside hydrolase
WP 131637124.1 glycoside hydrolase
CVPL010L 10353151 hypothetical protein
WP 023300115.1 hypothetical protein
WP 126503013.1 glycoside hydrolase
WP 126314875.1 glycoside hydrolase
WP 108367561.1 glycoside hydrolase
WP 063148887.1 glycoside hydrolase
WP 126489952.1 glycoside hydrolase
WP 032624251.1 glycoside hydrolase
WP 100148798.1 glycoside hydrolase
WP 063145900.1 MULTISPECIES: glycoside hydrolase
WP 046621525.1 glycoside hydrolase
WP 105322744.1 glycoside hydrolase
WP 023306209.1 hypothetical protein
WP 048966022.1 MULTISPECIES: glycoside hydrolase
WP 063163052.1 glycoside hydrolase
WP 063147323.1 MULTISPECIES: glycoside hydrolase
WP 045622665.1 MULTISPECIES: glycoside hydrolase
WP 038416063.1 MULTISPECIES: glycoside hydrolase
WP 063149952.1 MULTISPECIES: glycoside hydrolase
WP 126291503.1 glycoside hydrolase
WP 069596567.1 glycoside hydrolase
WP 127849869.1 glycoside hydrolase
WP 022648130.1 MULTISPECIES: glycoside hydrolase
WP 058686378.1 glycoside hydrolase
WP 063624260.1 glycoside hydrolase
WP 047637344.1 glycoside hydrolase
WP 045349604.1 MULTISPECIES: glycoside hydrolase
WP 017693418.1 MULTISPECIES: exo-poly-alpha-D-galacturonosidase
WP 134690405.1 glycoside hydrolase
WP 096205265.1 glycoside hydrolase
WP 062938940.1 glycoside hydrolase
EUM55884.1 hypothetical protein L361_01336
WP 047726559.1 glycoside hydrolase
WP 064489368.1 glycoside hydrolase
WP 063859231.1 glycoside hydrolase
WP 045339802.1 MULTISPECIES: glycoside hydrolase
WP 049110004.1 glycoside hydrolase
WP 126495663.1 glycoside hydrolase
WP 063133428.1 glycoside hydrolase
WP 032648327.1 MULTISPECIES: glycoside hydrolase
WP 058700710.1 glycoside hydrolase
WP 063135163.1 glycoside hydrolase
WP 090278373.1 glycoside hydrolase
WP 048247370.1 MULTISPECIES: glycoside hydrolase
WP 063132716.1 glycoside hydrolase
CVPL010L 10277221 hypothetical protein
WP 040117651.1 glycoside hydrolase
WP 023303742.1 MULTISPECIES: hypothetical protein
WP 132013047.1 glycoside hydrolase
WP 032634391.1 MULTISPECIES: glycoside hydrolase
WP 059452924.1 glycoside hydrolase
WP 075606860.1 glycoside hydrolase
WP 045341968.1 glycoside hydrolase
WP 058692077.1 glycoside hydrolase
WP 047733339.1 glycoside hydrolase
WP 017382348.1 exo-poly-alpha-D-galacturonosidase
WP 129541446.1 glycoside hydrolase
WP 063863973.1 glycoside hydrolase
Mild support
Polygalacturonase
Inner strip: Bacterial taxa
Bacillales
Enterobacteriales
Lactobacillales
Opitutales
Rhizobiales
Unclassified
Outgroups
Outer strip: Host type
C. varians larvae
C. texanus larvae
Non-Cephalotes associated
Outer labels: Gene functional annotation
Cephalotes-associated bacterial gene annotation
Function expected in focal trees
Function potentially related to the expected function in focal trees
Function unrelated to the expected function in focal trees

**S5 Fig. Pectate lyase and larval polygalacturonase protein phylogenetic trees showing examples of annotations well supported, mildly supported, and not supported.** Rooted phylogenetic tree of **(A)** pectate lyase and **(B)** polygalacturonase amino acid sequences from *Cephalotes*-associated symbionts. Sequences were obtained from a screening in IMG/M-ER against seven *Cephalotes* metagenomes. These sequences were used as queries for an IMG/M-ER BLAST against the 20 *Cephalotes* metagenomes sequenced and the 14 *Cephalotes* IGs and for a BLASTP against custom NCBI databases. A maximum likelihood method was used with 999 bootstrap iterations to build the trees. All bootstrap values are shown. Colors of host *Cephalotes* species (outer strip) are from [61]. **(A)** The annotation as pectate lyase is only well supported for one gene from *Staphylococcus* sp. JDR108L-110-1 IG, falling into a monophyletic group along with other pectate lyase-encoding genes from non-*Cephalotes* associated Bacillales. **(B)** The annotation as polygalacturonase is mildly supported for these three Enterobacteriales genes given that they are closely related to genes with unprecise annotation ("glycoside hydrolase").
