## Supplementary material for "In their sister’s footsteps: Taxonomic divergence obscures substantial functional overlap among the metabolically diverse symbiotic gut communities of adult and larval turtle ants": S6_Fig

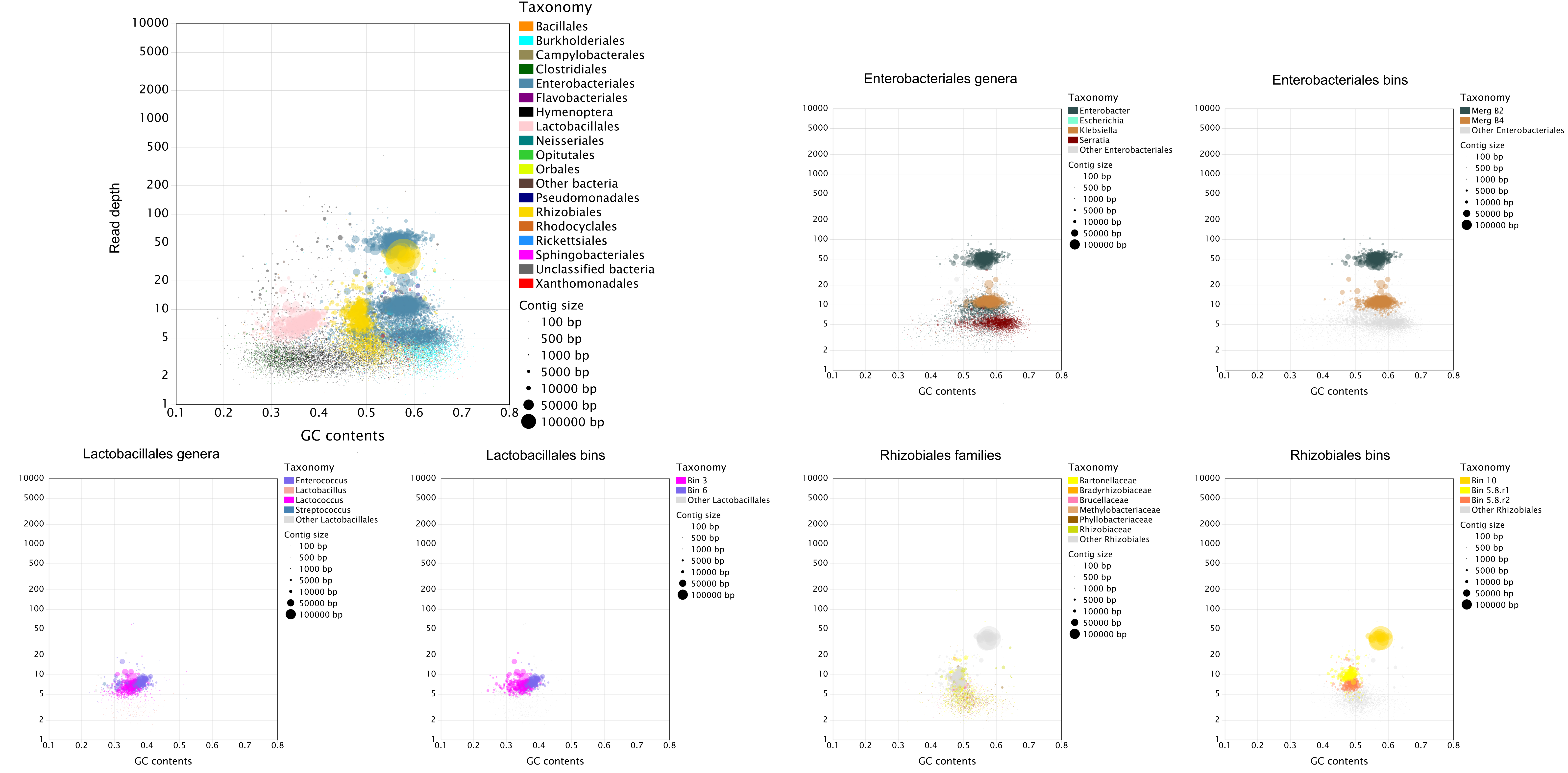
Taxonomy
Bacillales
Burkholderiales
Campylobacterales
Clostridiales
Enterobacteriales
Flavobacteriales
Hymenoptera
Lactobacillales
Neisseriales
Opitutales
Orbales
Other bacteria
Pseudomonadales
Rhizobiales
Rhodocyclales
Rickettsiales
Sphingobacteriales
Unclassified bacteria
Xanthomonadales
Contig size
100 bp
500 bp
1000 bp
5000 bp
10000 bp
50000 bp
100000 bp
Read depth
GC contents
Enterobacteriales genera
Enterobacter
Escherichia
Klebsiella
Serratia
Other Enterobacteriales
Enterobacteriales bins
Merg B2
Merg B4
Other Enterobacteriales
Lactobacillales genera
Enterococcus
Lactobacillus
Lactococcus
Streptococcus
Other Lactobacillales
Lactobacillales bins
Bin 3
Bin 6
Other Lactobacillales
Rhizobiales families
Bartonellaceae
Bradyrhizobiaceae
Brucellaceae
Methylobacteriaceae
Phyllobacteriaceae
Rhizobiaceae
Other Rhizobiales
Rhizobiales bins
Bin 10
Bin 5.8.r1
Bin 5.8.r2
Other Rhizobiales

**S6 Fig. Taxonomic description of MAGs in the *Cephalotes varians* PL010 larval gut metagenome**. Assembled scaffolds from the PL010 larval metagenome were taxonomically annotated using the USEARCH-IMG/M-ER pipeline [79–81]. See **S2 Fig** for a legend on GC-blobplots. The first GC-blobplot is a more detailed version of **Fig 2A**, with, additionally, scaffolds < 500 bp, and annotations of less represented taxonomic orders. The following pairs of GC-blobplots illustrate the possible classification of certain MAGs. Enterobacteriales and Lactobacillales MAGs could be classified at a lower taxonomic level, but it was not the case for Rhizobiales MAGs.
