## Supplementary material for "In their sister’s footsteps: Taxonomic divergence obscures substantial functional overlap among the metabolically diverse symbiotic gut communities of adult and larval turtle ants": S7_Fig

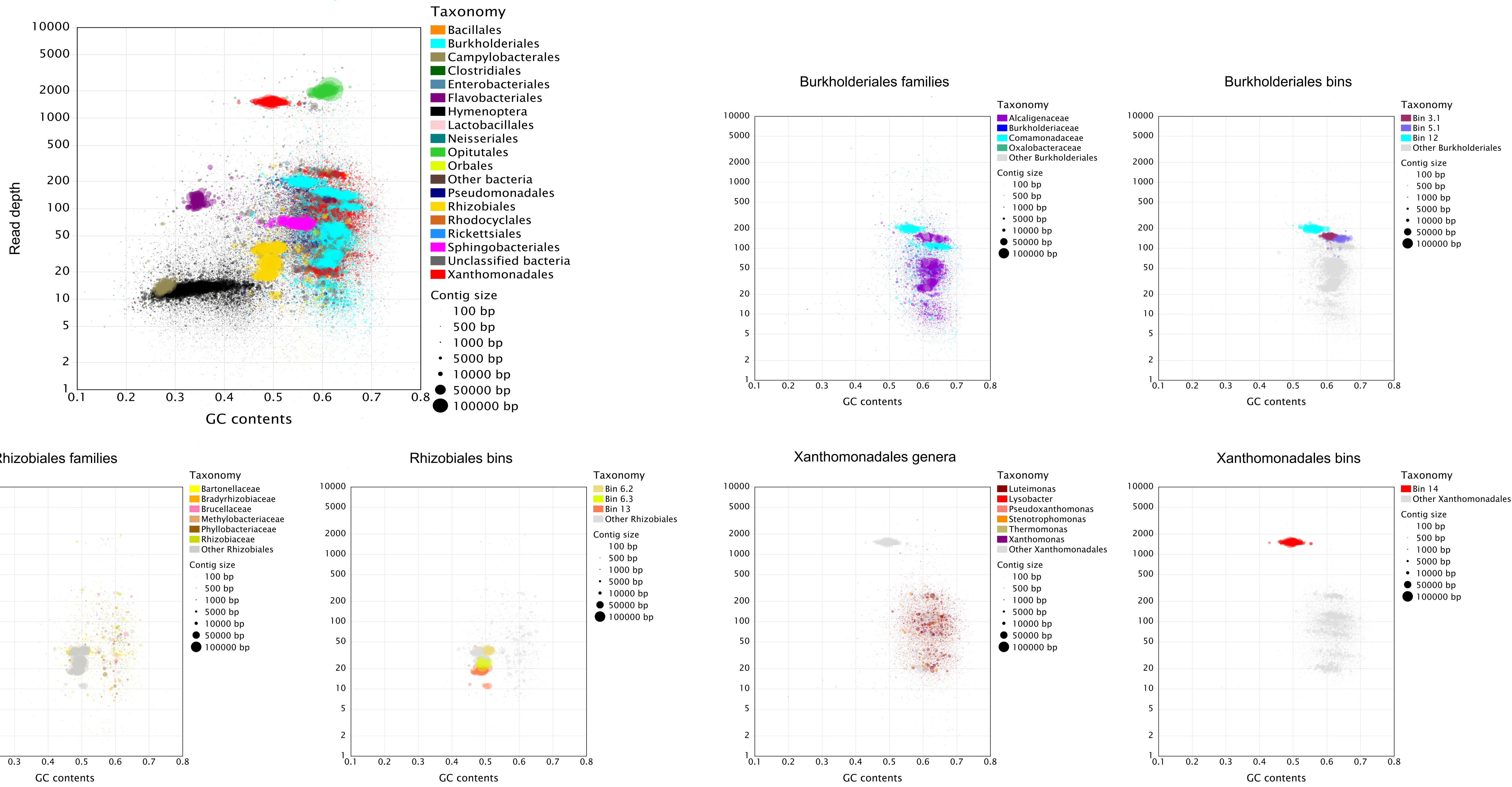
Taxonomy
Bacillales
Burkholderiales
Campylobacterales
Clostridiales
Enterobacteriales
Flavobacteriales
Hymenoptera
Lactobacillales
Neisseriales
Opitutales
Orbales
Other bacteria
Pseudomonadales
Rhizobiales
Rhodocyclales
Rickettsiales
Sphingobacteriales
Unclassified bacteria
Xanthomonadales
Contig size
100 bp
500 bp
1000 bp
5000 bp
10000 bp
50000 bp
100000 bp
Read depth
10000
5000
2000
1000
500
200
100
50
20
10
5
2
1
GC contents
0.1
0.2
0.3
0.4
0.5
0.6
0.7
0.8
Burkholderiales families
Taxonomy
Alcaligenaceae
Burkholderiaceae
Comamonadaceae
Oxalobacteraceae
Other Burkholderiales
Burkholderiales bins
Bin 3.1
Bin 5.1
Bin 12
Other Burkholderiales
Rhizobiales families
Bartonellaceae
Bradyrhizobiaceae
Brucellaceae
Methylobacteriaceae
Phyllobacteriaceae
Rhizobiaceae
Other Rhizobiales
Rhizobiales bins
Bin 6.2
Bin 6.3
Bin 13
Other Rhizobiales
Xanthomonadales genera
Luteimonas
Lysobacter
Pseudoxanthomonas
Stenotrophomonas
Thermomonas
Xanthomonas
Other Xanthomonadales
Xanthomonadales bins
Bin 14
Other Xanthomonadales

**S7 Fig. Taxonomic description of MAGs in the *Cephalotes varians* PL010 adult worker gut metagenome**. Assembled scaffolds from the PL010 adult metagenome were taxonomically annotated using the USEARCH-IMG/M-ER pipeline [79–81]. See **S2 Fig** for a legend on GC-blobplots. The first GC-blobplot is a more detailed version of **Fig 2A**, with, additionally, scaffolds < 500 bp, and annotations to less represented taxonomic orders. The following pairs of GC-blobplots illustrate the possible classification of certain MAGs. While Burkholderiales scaffolds are easy to assign to bacterial families, classifying Rhizobiales and Xanthomonadales MAGs is more difficult.
