## Supplementary material for "In their sister’s footsteps: Taxonomic divergence obscures substantial functional overlap among the metabolically diverse symbiotic gut communities of adult and larval turtle ants": S8_Fig

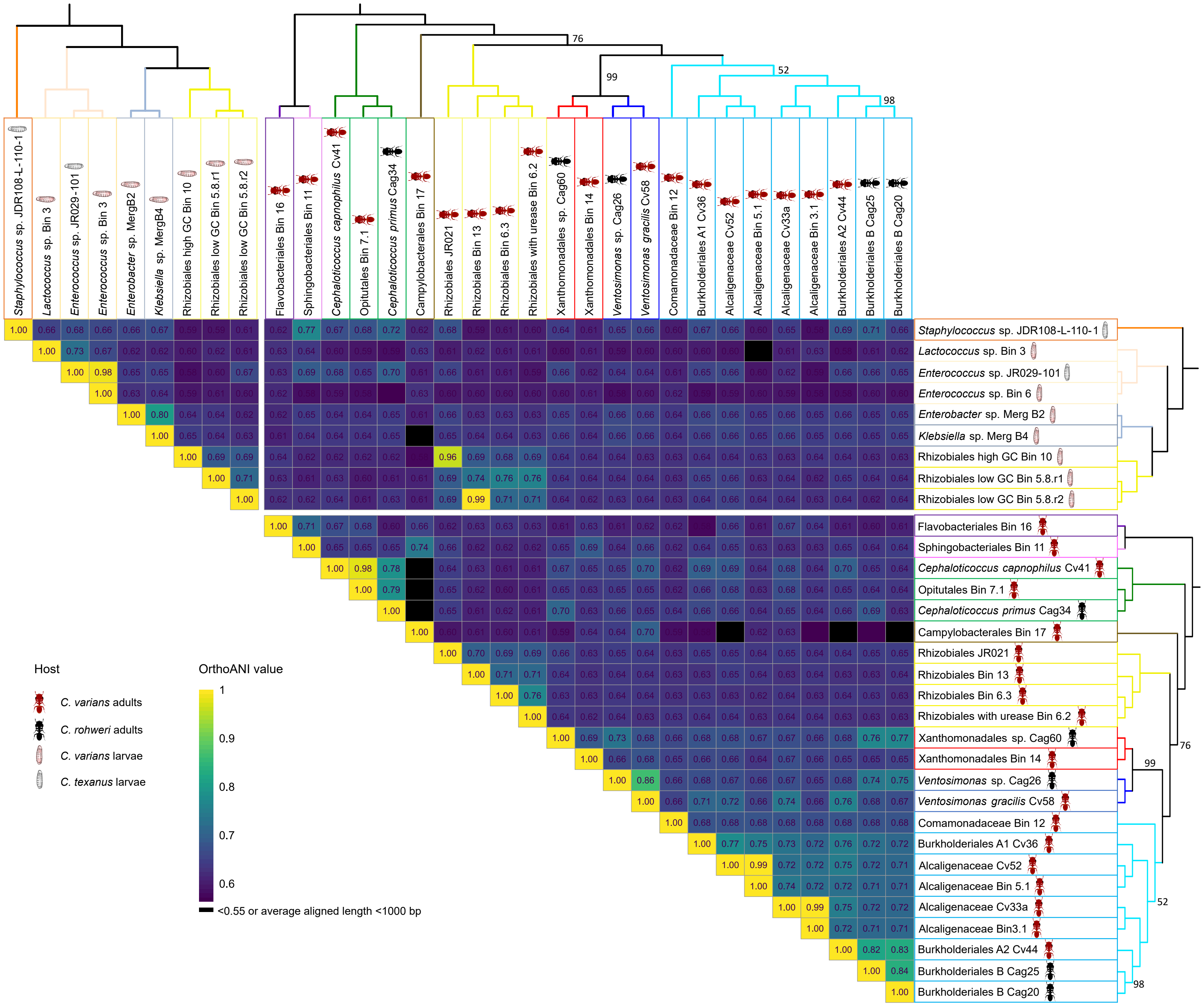
Staphylococcus sp. JDR108-L-110-1
Lactococcus sp. Bin 3
Enterococcus sp. JR029-101
Enterococcus sp. Bin 3
Enterococcus sp. Bin 6
Enterobacter sp. MergB2
Enterobacter sp. Merg B2
Klebsiella sp. MergB4
Klebsiella sp. Merg B4
Rhizobiales high GC Bin 10
Rhizobiales low GC Bin 5.8.r1
Rhizobiales low GC Bin 5.8.r2
Flavobacteriales Bin 16
Sphingobacteriales Bin 11
Cephaloticoccus capnophilus Cv41
Opitutales Bin 7.1
Cephaloticoccus primus Cag34
Campylobacterales Bin 17
Rhizobiales JR021
Rhizobiales Bin 13
Rhizobiales Bin 6.3
Rhizobiales with urease Bin 6.2
Xanthomonadales sp. Cag60
Xanthomonadales Bin 14
Ventosimonas sp. Cag26
Ventosimonas gracilis Cv58
Comamonadaceae Bin 12
Burkholderiales A1 Cv36
Alcaligenaceae Cv52
Alcaligenaceae Bin 5.1
Alcaligenaceae Cv33a
Alcaligenaceae Bin 3.1
Alcaligenaceae Bin3.1
Burkholderiales A2 Cv44
Burkholderiales B Cag25
Burkholderiales B Cag20
76
99
52
98
Host
C. varians adults
C. rohweri adults
C. varians larvae
C. texanus larvae
OrthoANI value
1
0.9
0.8
0.7
0.6
<0.55 or average aligned length <1000 bp

**S8 Fig. Pairwise Average Nucleotide Identity (ANI) calculated among *Cephalotes*-associated symbiont genomes.** Assembled scaffolds from *C. varians* PL010 metagenomes were assigned to MAGs (with the mention “Bin” or “Merg”) using the Anvi’o-CONCOCT pipeline. Additional IGs from *C. varians*, *C. texanus* and *C. rohweri* were included. The bacterial phylogenetic trees were inferred based on seven bacterial marker gene sequences using a maximum likelihood method and 999 bootstrap iterations. Bootstrap values <100 are shown on the tree. OrthoANI values were calculated as described in Yoon et al. (2017) [85]. Higher orthoANIs indicate higher levels of genomic similarity among symbionts.
