## Supplementary material for "In their sister’s footsteps: Taxonomic divergence obscures substantial functional overlap among the metabolically diverse symbiotic gut communities of adult and larval turtle ants": S9_Fig

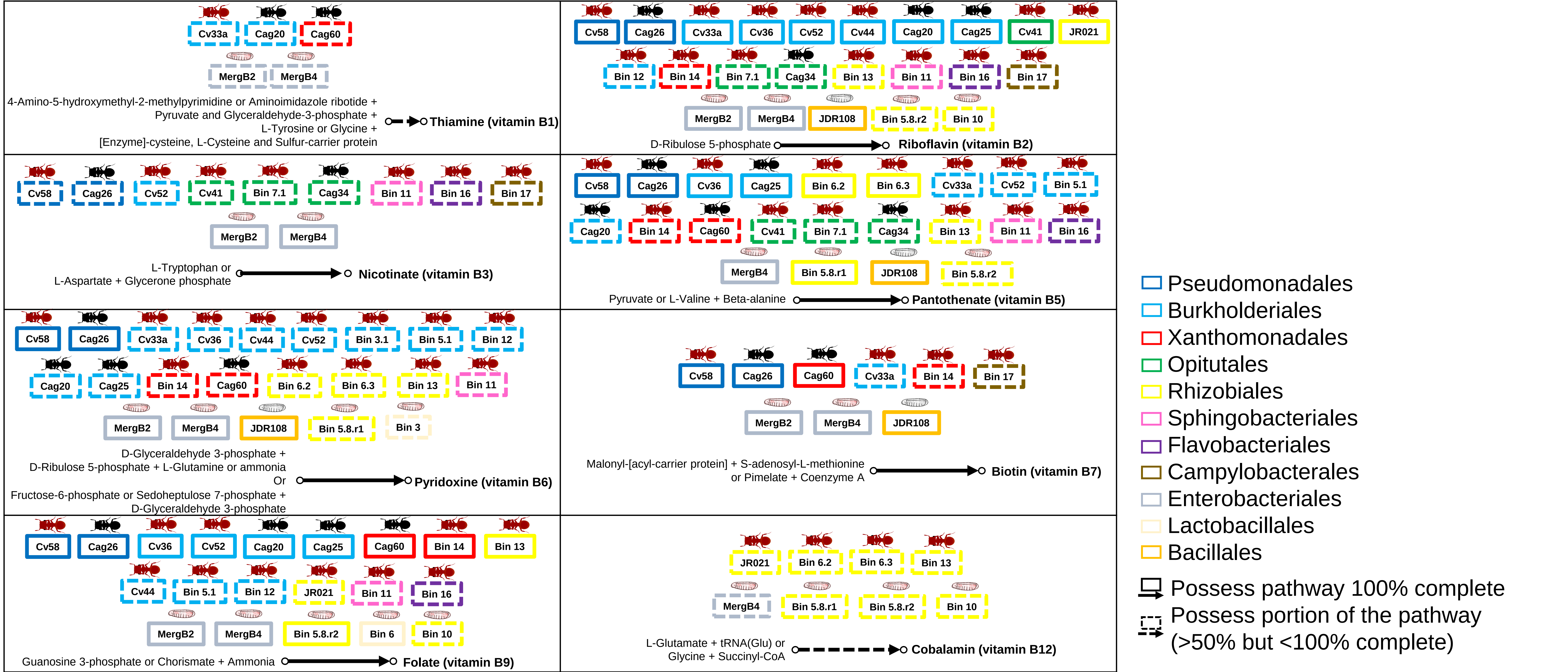
Cv33a
Cag20
Cag60
MergB2
MergB4
4-Amino-5-hydroxymethyl-2-methylpyrimidine or Aminoimidazole ribotide +
Pyruvate and Glyceraldehyde-3-phosphate +
L-Tyrosine or Glycine +
[Enzyme]-cysteine, L-Cysteine and Sulfur-carrier protein
Thiamine (vitamin B1)
Cv58
Cag26
Cv33a
Cv36
Cv52
Cv44
Cag20
Cag25
Cv41
JR021
Bin 12
Bin 14
Bin 7.1
Cag34
Bin 13
Bin 11
Bin 16
Bin 17
MergB2
MergB4
JDR108
Bin 5.8.r2
Bin 10
D-Ribulose 5-phosphate
Riboflavin (vitamin B2)
Cv58
Cag26
Cv52
Cv41
Bin 7.1
Cag34
Bin 11
Bin 16
Bin 17
MergB2
MergB4
L-Tryptophan or
L-Aspartate + Glycerone phosphate
Nicotinate (vitamin B3)
Cv58
Cag26
Cv36
Cag25
Bin 6.2
Bin 6.3
Cv33a
Cv52
Bin 5.1
Cag20
Bin 14
Cag60
Cv41
Bin 7.1
Cag34
Bin 13
Bin 11
Bin 16
MergB4
Bin 5.8.r1
JDR108
Bin 5.8.r2
Pyruvate or L-Valine + Beta-alanine
Pantothenate (vitamin B5)
Cv58
Cag26
Cv33a
Cv36
Cv44
Cv52
Bin 3.1
Bin 5.1
Bin 12
Cag20
Cag25
Bin 14
Cag60
Bin 6.2
Bin 6.3
Bin 13
Bin 11
MergB2
MergB4
JDR108
Bin 5.8.r1
Bin 3
D-Glyceraldehyde 3-phosphate +
D-Ribulose 5-phosphate + L-Glutamine or ammonia
Or
Fructose-6-phosphate or Sedoheptulose 7-phosphate +
D-Glyceraldehyde 3-phosphate
Pyridoxine (vitamin B6)
Cv58
Cag26
Cag60
Cv33a
Bin 14
Bin 17
MergB2
MergB4
JDR108
Malonyl-[acyl-carrier protein] + S-adenosyl-L-methionine
or Pimelate + Coenzyme A
Biotin (vitamin B7)
Cv58
Cag26
Cv36
Cv52
Cag20
Cag25
Cag60
Bin 14
Bin 13
Cv44
Bin 5.1
Bin 12
JR021
Bin 11
Bin 16
MergB2
MergB4
Bin 5.8.r2
Bin 6
Bin 10
Guanosine 3-phosphate or Chorismate + Ammonia
Folate (vitamin B9)
JR021
Bin 6.2
Bin 6.3
Bin 13
MergB4
Bin 5.8.r1
Bin 5.8.r2
Bin 10
L-Glutamate + tRNA(Glu) or
Glycine + Succinyl-CoA
Cobalamin (vitamin B12)
Pseudomonadales
Burkholderiales
Xanthomonadales
Opitutales
Rhizobiales
Sphingobacteriales
Flavobacteriales
Campylobacterales
Enterobacteriales
Lactobacillales
Bacillales
Possess pathway 100% complete
Possess portion of the pathway
(>50% but <100% complete)

**S9 Fig. B-vitamin biosynthesis pathways encoded in *Cephalotes*-associated genomes.** MAGs (with the mention "Bin" or "Merg") and IGs next to an arrow indicates that they possess the genes for the focal step. Dashed squares correspond to pathways half encoded to almost complete. The color of turtle ant adults and larvae correspond to the species symbionts were sampled from, as in **S8 Fig**. Note that many *Cephalotes*-associated symbionts have retained genes for B-vitamin biosynthesis.
