## Supplementary material for "In their sister’s footsteps: Taxonomic divergence obscures substantial functional overlap among the metabolically diverse symbiotic gut communities of adult and larval turtle ants": S10_Fig

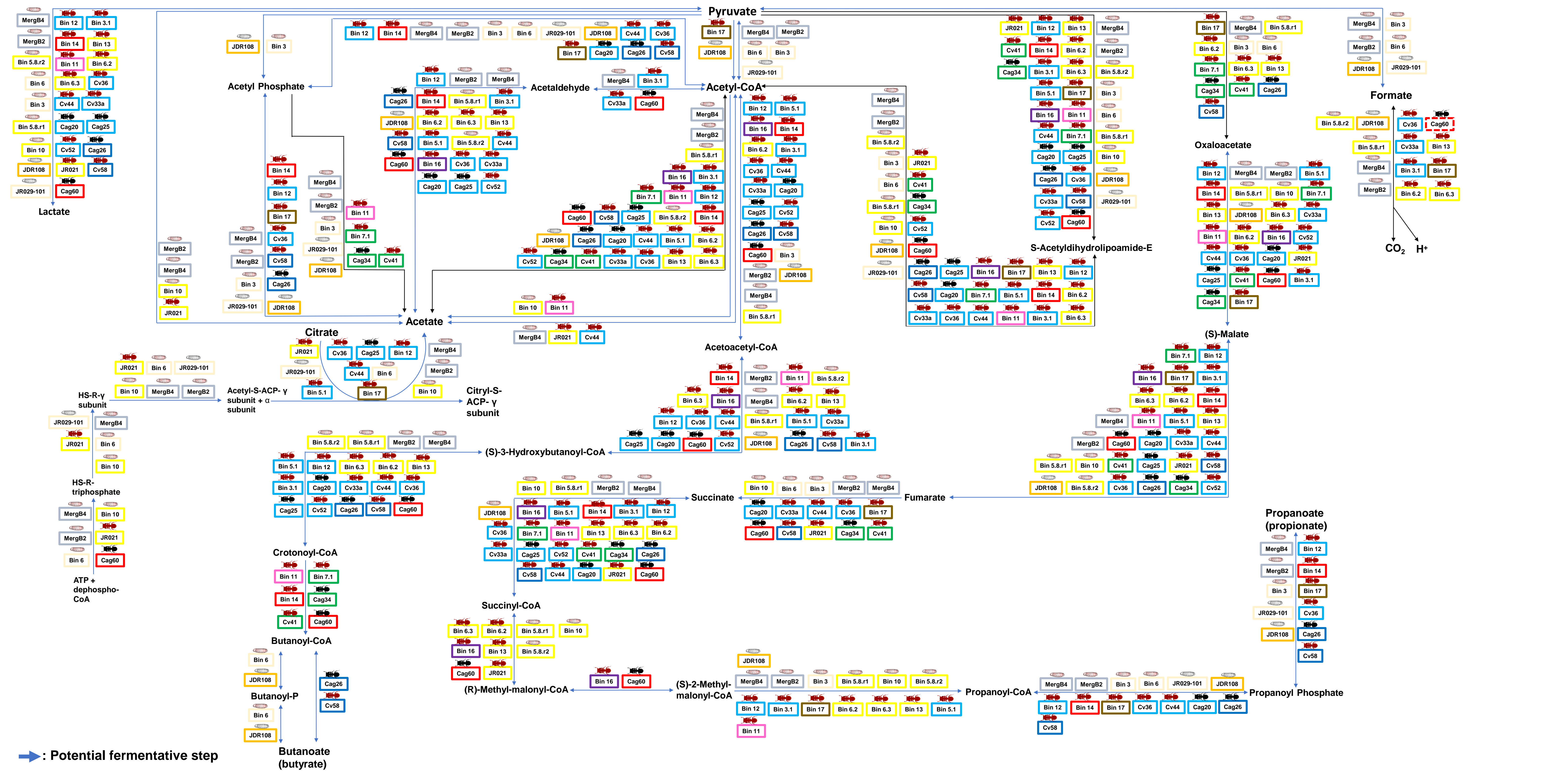

**S10 Fig. Organic acid metabolism pathways encoded in *Cephalotes*-associated genomes.** MAGs (with the mention “Bin” or “Merg”) and IGs next to an arrow indicates that they possess the genes for the focal step. Blue arrows are for steps that can be fermentative, taking place in anaerobic environments. The color of turtle ant adults and larvae correspond to the species symbionts were sampled from, as in **S8 Fig**. Note that many *Cephalotes*-associated symbionts have retained genes for organic acid metabolism, and many can ferment pyruvate to lactate and acetate. Symbionts from larvae (plus Rhizobiales JR021-5/Bin10) can additionally ferment citrate and ferment pyruvate to produce formate.
