## Supplementary material for "In their sister’s footsteps: Taxonomic divergence obscures substantial functional overlap among the metabolically diverse symbiotic gut communities of adult and larval turtle ants": S11_Fig

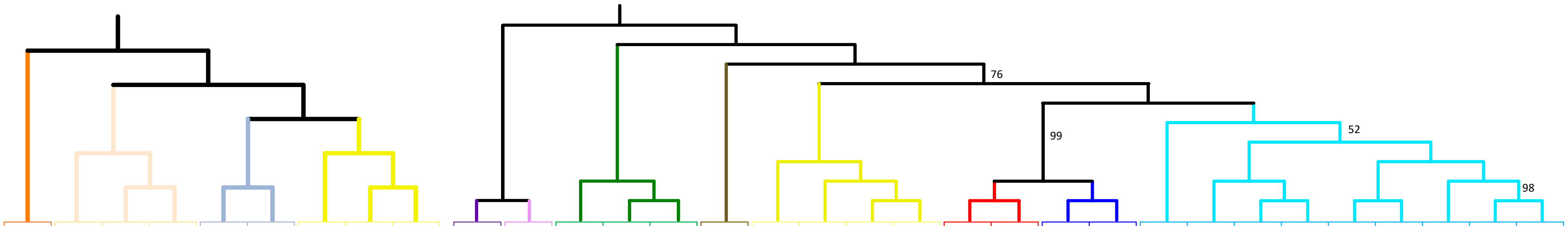


| Pathway | Steps | *Staphylococcus* sp. JDR108-L-110-1 | Lactobacillales Bin 3 | *Enterococcus* sp. JR029-101 | Lactobacillales Bin 6 | Enterobacteriales Merg B2 | Enterobacteriales Merg B4 | Rhizobiales high GC Bin 10 | Rhizobiales low GC Bin 5.8.r1 | Rhizobiales low GC Bin 5.8.r2 | Flavobacteriales Bin 16 | Sphingobacteriales Bin 11 | *Cephaloticoccus capnophilus* Cv41 | Opitutales Bin 7.1 | *Cephaloticoccus primus* Cag34 | Campylobacterales Bin 17 | Rhizobiales JR021 | Rhizobiales Bin 13 | Rhizobiales Bin 6.3 | Rhizobiales with urease Bin 6.2 | Xanthomonadales sp. Cag60 | Xanthomonadales Bin 14 | *Ventosimonas* sp. Cag26 | *Ventosimonas gracilis* Cv58 | Burkholderiales Bin 12 | Burkholderiales A1 Cv36 | Burkholderiales C Cv52 | Burkholderiales Bin 5.1 | Burkholderiales B Cv33a | Burkholderiales Bin 3.1 | Burkholderiales A2 Cv44 | Burkholderiales B Cag25 | Burkholderiales B Cag20 | Host genome |
|---|---|---|---|---|---|---|---|---|---|---|---|---|---|---|---|---|---|---|---|---|---|---|---|---|---|---|---|---|---|---|---|---|---|---|
| Purine degradation | step1 | | | *ade* | *ade* | | *ade* | | *ade* | | | | | | | | | | | *ade* | | | | | | | | | | | | | | |
| | step2 | | | *guaD* | *guaD* | *guaD* | *guaD* | | | | | | | | | | | | | | | | | | | | | | *guaD* | *guaD* | | | | *guaD* |
| Uricolytic pathway | step3 | | | *xdhABC* | | *hpxDE* | *hpxDE* | | | | | | | | | | | | | | *xdhABC* | | | | | | | | *xdhABC* | *xdhABC* | | *xdhABC* | | XDH |
| | step4 | | | | | *hpxO* | *hpxO* | | | | | | | | | | | | | | | | | | *puuD* | | | | *puuD* | *puuD* | | | | *uaZ* |
| | step5 | | | | | *hpxT* | *hpxT* | | | | | | | | | | | | | | | | | | *uraH* | | | | *uraH* | *uraH* | | | | *uraH* |
| | step6 | | | | | *hpxQ* | *hpxQ* | | | | | | | | | | | | | | | PRHOXNB | | | PRHOXNB | | PRHOXNB | PRHOXNB | PRHOXNB | PRHOXNB | PRHOXNB | PRHOXNB | PRHOXNB | |
| | step6' | | | | | *hpxA* | *hpxA* | | | | | | | | | | | | | | *hpxA* | *hpxA* | | | *hpxA* | | | | *hpxA* | *hpxA* | *hpxA* | | | |
| | step7 | *allB* | | *allB* | *allB* | *hpxB* | *hpxB* | | *allB* | *allB* | | | | | | | | *allB* | | | | *puuE* | | | *puuE* | | | | *puuE* | *puuE* | *puuE* | *puuE* | *puuE* | *allB* |
| | step8 | | | | | | | | | | | | | | | | | | | | | | | | | | | | *alc* | *alc* | | | | |
| Ureidoglycine/ureidoglycolate route | step9 | *allC* | | *allC* | *allC* | *hpxK* | *hpxK* | | *allC* | *allC* | | | | | | | | *allC* | | | | | | | | | *allC* | *allC* | *allC* | *allC* | *allC* | *allC* | *allC* | |
| | step10 | *pucG* | *pucG* | *pucG* | *pucG* | *hpxJ* | *hpxJ* | | | | | | *pucG* | *pucG* | *pucG* | *pucG* | | | | | | | | | | | | | | | | | | |
| | step11 | *ylbA* | | *ylbA* | *ylbA* | | | | | | | | | | | | | | | *ylbA* | | | | | | | | | *ylbA* | *ylbA* | | | | |
| | step12 | | | | | | | | | | | | | | | | | | | | | | | | | *allA* | | | *allA* | *allA* | | | *ugl* | |
| Urease | step13 | *ureABC* | | | | *ureABC* | *ureABC* | | | | | | *ureABC* | *ureABC* | *ureABC* | | | | | *ureABC* | *ureABC* | | | | | | | | | | | | | |
| Oxalurate pathway | step14 | *allD* | | *allD* | *allD* | | | | | | | | | | | | | | | | | | | | | | | | | | | | | |
| | step15 | | | | | *hpxY* | *hpxY* | | | | | | | | | | | | | | | | | | | | | | | | | | | |
| | step16 | | | | | *hpxW* | *hpxW* | | | | | | | | | | | | | | | | | | | | | | | | | | | |
| | step17 | *arcC* | *arcC* | *arcC* | *arcC* | *arcC* | | | | | | | | | | | | | | | | | | | | | | | | | | | | |

**S11 Fig. Gene-level N-recycling pathways in *Cephalotes*-associated symbiont genomes.** Presence of genes encoding steps of N-recycling in each symbiont genome is indicated in squares. Black squares without any gene name are for genes not found in focal genomes. Gene names with a white font and without background color correspond to genes not found in our screenings, but likely to be present because they were found in genomes of highly related *Cephalotes*-associated symbionts (see **S8 Fig**). For a visual description of enzymatic step numbers, see **Fig 6B**. The bacterial phylogenetic trees were inferred based on seven bacterial marker gene sequences using a maximum likelihood method and 999 bootstrap iterations. Bootstrap values <100 are shown on the tree.
