## Supplementary material for "In their sister’s footsteps: Taxonomic divergence obscures substantial functional overlap among the metabolically diverse symbiotic gut communities of adult and larval turtle ants": S12_Fig

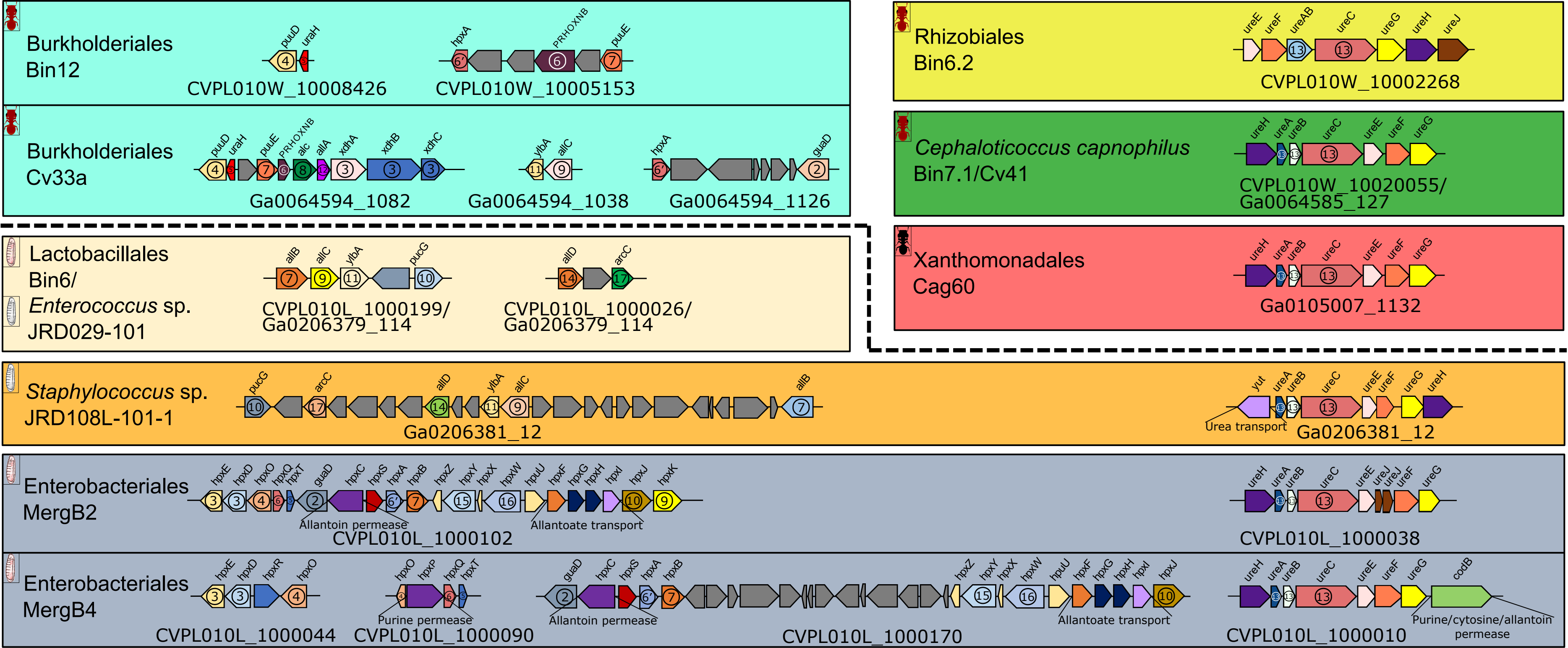

Burkholderiales Bin12
puuD uraH
CVPL010W_10008426
hpxA PRHOXNB puuE
CVPL010W_10005153
Burkholderiales Cv33a
puuD uraH puuE PRHOXNB alc allA xdhA xdhB xdhC
Ga0064594_1082
ylbA allC
Ga0064594_1038
hpxA guaD
Ga0064594_1126
Rhizobiales Bin6.2
ureE ureF ureAB ureC ureG ureH ureJ
CVPL010W_10002268
Cephaloticoccus capnophilus Bin7.1/Cv41
ureH ureA ureB ureC ureE ureF ureG
CVPL010W_10020055/ Ga0064585_127
Lactobacillales Bin6/ Enterococcus sp. JRD029-101
allB allC ylbA pucG
CVPL010L_1000199/ Ga0206379_114
allD arcC
CVPL010L_1000026/ Ga0206379_114
Xanthomonadales Cag60
ureH ureA ureB ureC ureE ureF ureG
Ga0105007_1132
Staphylococcus sp. JRD108L-101-1
pucG arcC allD ylbA allC allB
Ga0206381_12
yut ureA ureB ureC ureE ureF ureG ureH
Urea transport
Ga0206381_12
Enterobacteriales MergB2
hpxE hpxD hpxO hpxQ hpxT guaD hpxC hpxS hpxA hpxB hpxZ hpxY hpxX hpxW hpuU hpxF hpxG hpxH hpxI hpxJ hpxK
Allantoin permease
Allantoate transport
CVPL010L_1000102
ureH ureA ureB ureC ureE ureJ ureJ ureF ureG
CVPL010L_1000038
Enterobacteriales MergB4
hpxE hpxD hpxR hpxO
CVPL010L_1000044
hpxO hpxP hpxQ hpxT
Purine permease
CVPL010L_1000090
guaD hpxC hpxS hpxA hpxB hpxZ hpxY hpxX hpxW hpuU hpxF hpxG hpxH hpxI hpxJ
Allantoin permease
Allantoate transport
CVPL010L_1000170
ureH ureA ureB ureC ureE ureF ureG codB
Purine/cytosine/allantoin permease
CVPL010L_1000010

**S12 Fig. Clusters of genes coding for steps of N-recycling pathways from larval and adult gut symbionts.** Step numbers and gene names correspond to the ones shown in **Fig 6B**.
