## Supplementary material for "In their sister’s footsteps: Taxonomic divergence obscures substantial functional overlap among the metabolically diverse symbiotic gut communities of adult and larval turtle ants": S13_Fig

Scaffold: CVPL010L_1000153, MergB4 Enterobacteriales

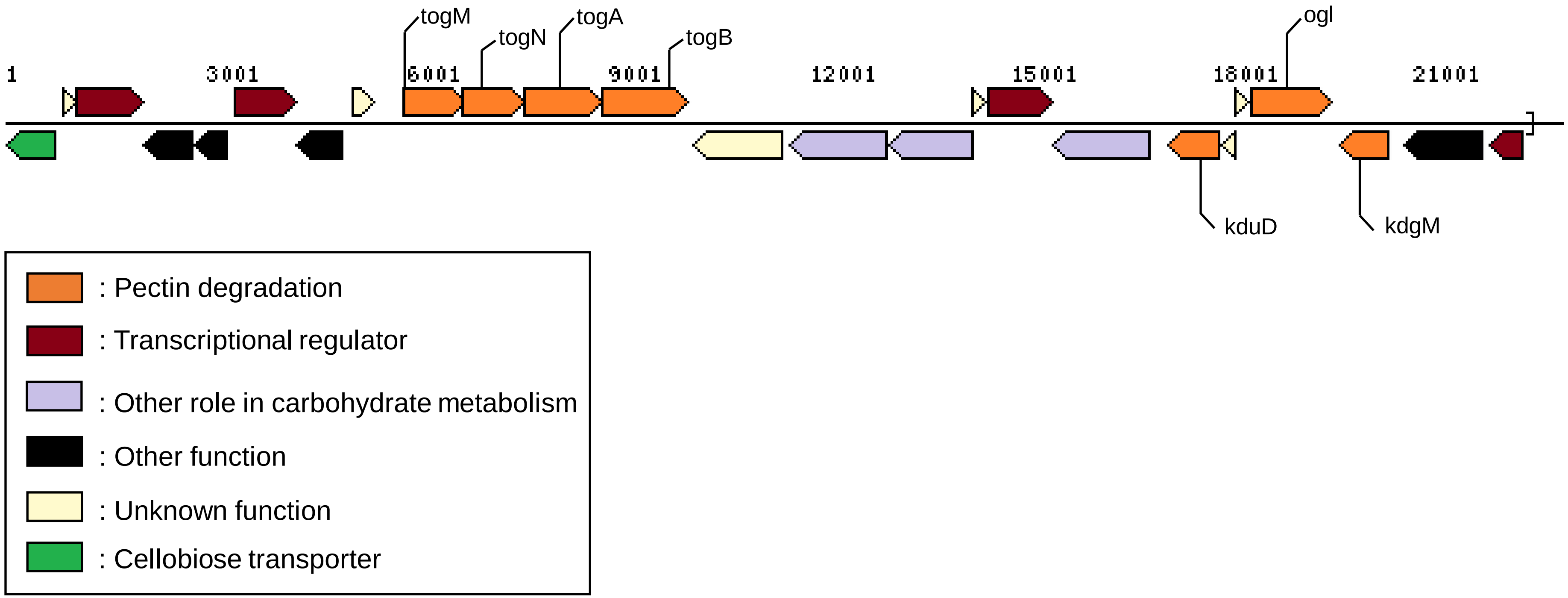

**S13 Fig. Gene cluster from the larval gut metagenome including genes to import pectate and degrade galacturonate.** The scaffold is from the Enterobacteriales order, symbiont *Klebsiella* sp. MergB4 MAG. For a visual description of gene names and their encoded enzymes, see **Fig 7A**.
