## Supplementary material for "In their sister’s footsteps: Taxonomic divergence obscures substantial functional overlap among the metabolically diverse symbiotic gut communities of adult and larval turtle ants": S14_Fig

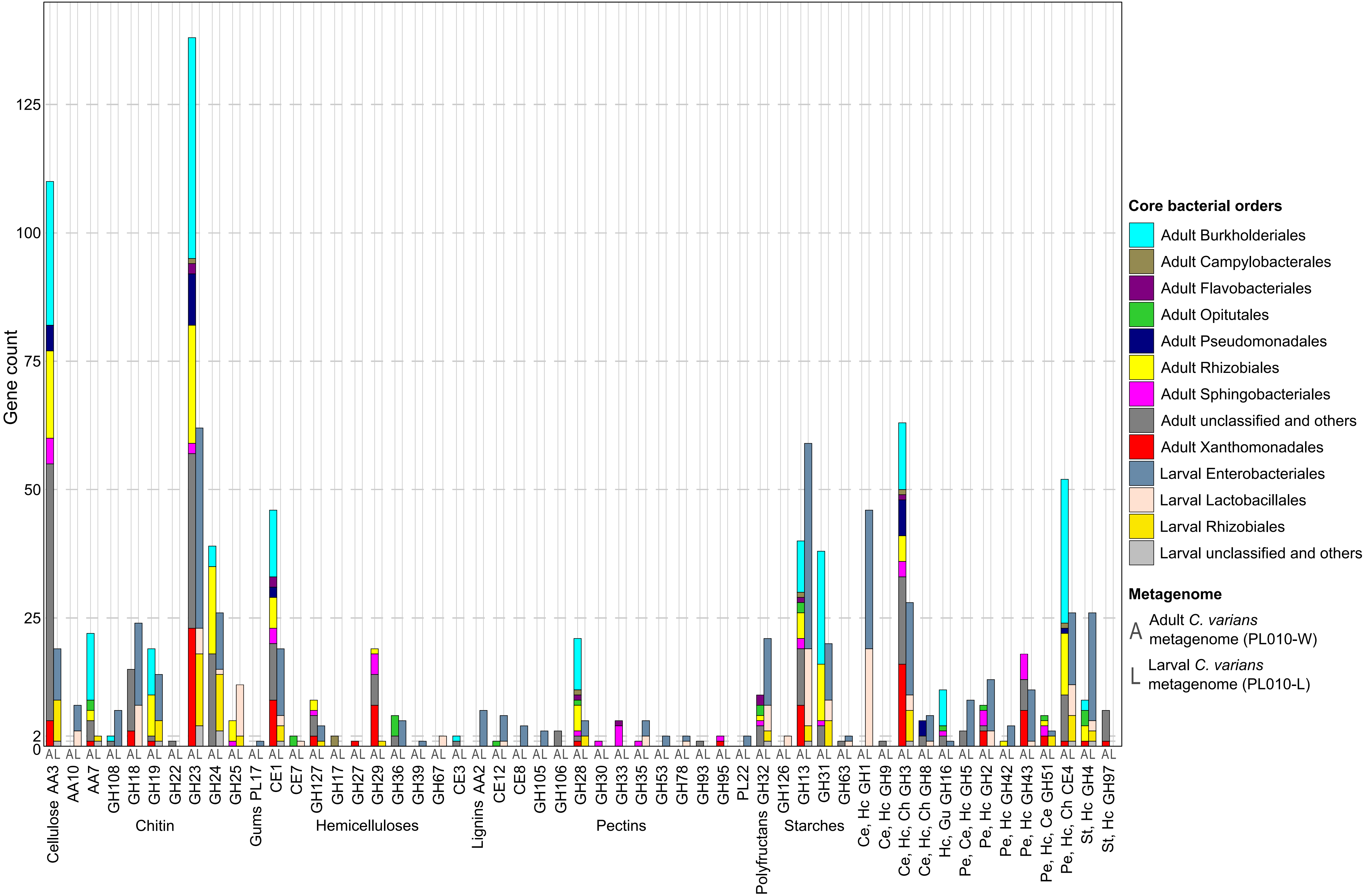

Gene count
Cellulose AA3
AA10
AA7
GH108
GH18
GH19
GH22
GH23
GH24
GH25
Chitin
Gums PL17
CE1
CE7
GH127
GH17
GH27
GH29
GH36
GH39
GH67
CE3
Hemicelluloses
Lignins AA2
CE12
CE8
GH105
GH106
GH28
GH30
GH33
GH35
GH53
GH78
GH93
GH95
PL22
Pectins
Polyfructans GH32
GH126
GH13
GH31
GH63
Starches
Ce, Hc GH1
Ce, Hc GH9
Ce, Hc, Ch GH3
Ce, Hc, Ch GH8
Hc, Gu GH16
Pe, Ce, Hc GH5
Pe, Hc GH2
Pe, Hc GH42
Pe, Hc GH43
Pe, Hc, Ce GH51
Pe, Hc, Ch CE4
St, Hc GH4
St, Hc GH97
Core bacterial orders
Adult Burkholderiales
Adult Campylobacterales
Adult Flavobacteriales
Adult Opitutales
Adult Pseudomonadales
Adult Rhizobiales
Adult Sphingobacteriales
Adult unclassified and others
Adult Xanthomonadales
Larval Enterobacteriales
Larval Lactobacillales
Larval Rhizobiales
Larval unclassified and others
Metagenome
A Adult C. varians metagenome (PL010-W)
L Larval C. varians metagenome (PL010-L)

**S14 Fig. Carbohydrate Active Enzyme (CAZy) screening results for *Cephalotes varians* PL010 larval and adult worker gut metagenomes.** The metagenomes were screened against the HMMER, DIAMOND and Hotpep databases for CAZy families (CAZymes) using the dbCAN2 metaserver (v7, January 2019) [101]. Catabolic CAZymes were categorized based on the type of substrates the enzymes can degrade. Degradations are as follow: "St": Starch, "HC": Hemicellulose, "Ce": Cellulose, "Pe": Pectin, "Pf": Polyfructans, "Gu": Gums, "Ch": Chitin, "Li": Lignin.
