## Supplementary material for "In their sister’s footsteps: Taxonomic divergence obscures substantial functional overlap among the metabolically diverse symbiotic gut communities of adult and larval turtle ants": S15_Fig

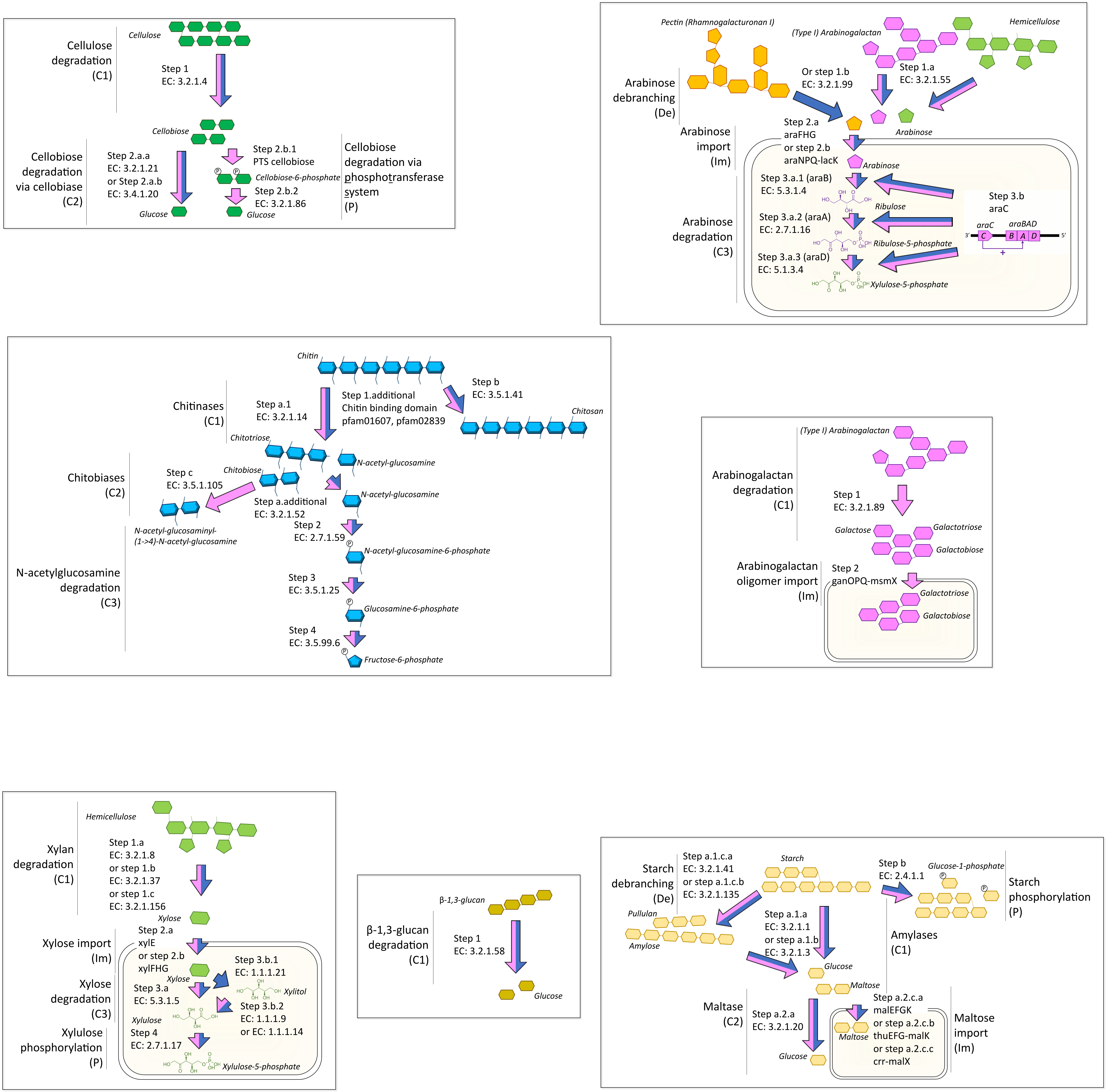
Cellulose degradation (C1)
Cellulose
Step 1
EC: 3.2.1.4
Cellobiose
Cellobiose degradation via cellobiase (C2)
Step 2.a.a
EC: 3.2.1.21
or Step 2.a.b
EC: 3.4.1.20
Glucose
Step 2.b.1
PTS cellobiose
Cellobiose-6-phosphate
Step 2.b.2
EC: 3.2.1.86
Glucose
Cellobiose degradation via phosphotransferase system (P)
Pectin (Rhamnogalacturonan I)
(Type I) Arabinogalactan
Hemicellulose
Arabinose debranching (De)
Or step 1.b
EC: 3.2.1.99
Step 1.a
EC: 3.2.1.55
Arabinose
Arabinose import (Im)
Step 2.a
araFHG
or step 2.b
araNPQ-lacK
Arabinose
Arabinose degradation (C3)
Step 3.a.1 (araB)
EC: 5.3.1.4
Ribulose
Step 3.a.2 (araA)
EC: 2.7.1.16
Ribulose-5-phosphate
Step 3.a.3 (araD)
EC: 5.1.3.4
Xylulose-5-phosphate
Step 3.b
araC
araC
araBAD
3'
C
B
A
D
5'
+
Chitin
Chitinases (C1)
Step a.1
EC: 3.2.1.14
Step 1.additional
Chitin binding domain
pfam01607, pfam02839
Step b
EC: 3.5.1.41
Chitosan
Chitotriose
N-acetyl-glucosamine
Chitobiases (C2)
Step c
EC: 3.5.1.105
Chitobiose
Step a.additional
EC: 3.2.1.52
N-acetyl-glucosamine
N-acetyl-glucosaminyl-(1->4)-N-acetyl-glucosamine
Step 2
EC: 2.7.1.59
N-acetyl-glucosamine-6-phosphate
N-acetylglucosamine degradation (C3)
Step 3
EC: 3.5.1.25
Glucosamine-6-phosphate
Step 4
EC: 3.5.99.6
Fructose-6-phosphate
(Type I) Arabinogalactan
Arabinogalactan degradation (C1)
Step 1
EC: 3.2.1.89
Galactose
Galactotriose
Galactobiose
Arabinogalactan oligomer import (Im)
Step 2
ganOPQ-msmX
Galactotriose
Galactobiose
Hemicellulose
Xylan degradation (C1)
Step 1.a
EC: 3.2.1.8
or step 1.b
EC: 3.2.1.37
or step 1.c
EC: 3.2.1.156
Xylose
Xylose import (Im)
Step 2.a
xylE
or step 2.b
xylFHG
Xylose
Xylose degradation (C3)
Step 3.a
EC: 5.3.1.5
Step 3.b.1
EC: 1.1.1.21
Xylitol
Step 3.b.2
EC: 1.1.1.9
or EC: 1.1.1.14
Xylulose
Xylulose phosphorylation (P)
Step 4
EC: 2.7.1.17
Xylulose-5-phosphate
β-1,3-glucan
β-1,3-glucan degradation (C1)
Step 1
EC: 3.2.1.58
Glucose
Starch debranching (De)
Step a.1.c.a
EC: 3.2.1.41
or step a.1.c.b
EC: 3.2.1.135
Starch
Pullulan
Amylose
Step a.1.a
EC: 3.2.1.1
or step a.1.b
EC: 3.2.1.3
Step b
EC: 2.4.1.1
Glucose-1-phosphate
Starch phosphorylation (P)
Amylases (C1)
Glucose
Maltose
Maltase (C2)
Step a.2.a
EC: 3.2.1.20
Glucose
Step a.2.c.a
malEFGK
or step a.2.c.b
thuEFG-malK
or step a.2.c.c
crr-malX
Maltose
Maltose import (Im)

**S15 Fig. Pathways for carbohydrate catabolism screened in *Cephalotes varians* PL010 larval and adult metagenomes.** Results at the symbiont genome-level can be seen in **S16 Fig**. Pink arrows are for steps encoded in the larval metagenome, and blue arrows are for steps encoded in the adult metagenome. Note that homogalacturonan catabolic pathway can be found in **Fig 7A**. “Es”: Esterification, “C1”: Cut on a polymer backbone, “C2”: Cut on a disaccharide, “De”: Debranching of a polymer side chain, “Im”: Import of disaccharides or monosaccharides in a bacterial cytoplasm, “P”: Phosphorylation of a carbohydrate, “C3”: Cut of a carbohydrate ring and transformation into an assimilable compound.
