## Supplementary material for "In their sister’s footsteps: Taxonomic divergence obscures substantial functional overlap among the metabolically diverse symbiotic gut communities of adult and larval turtle ants": S16_Fig

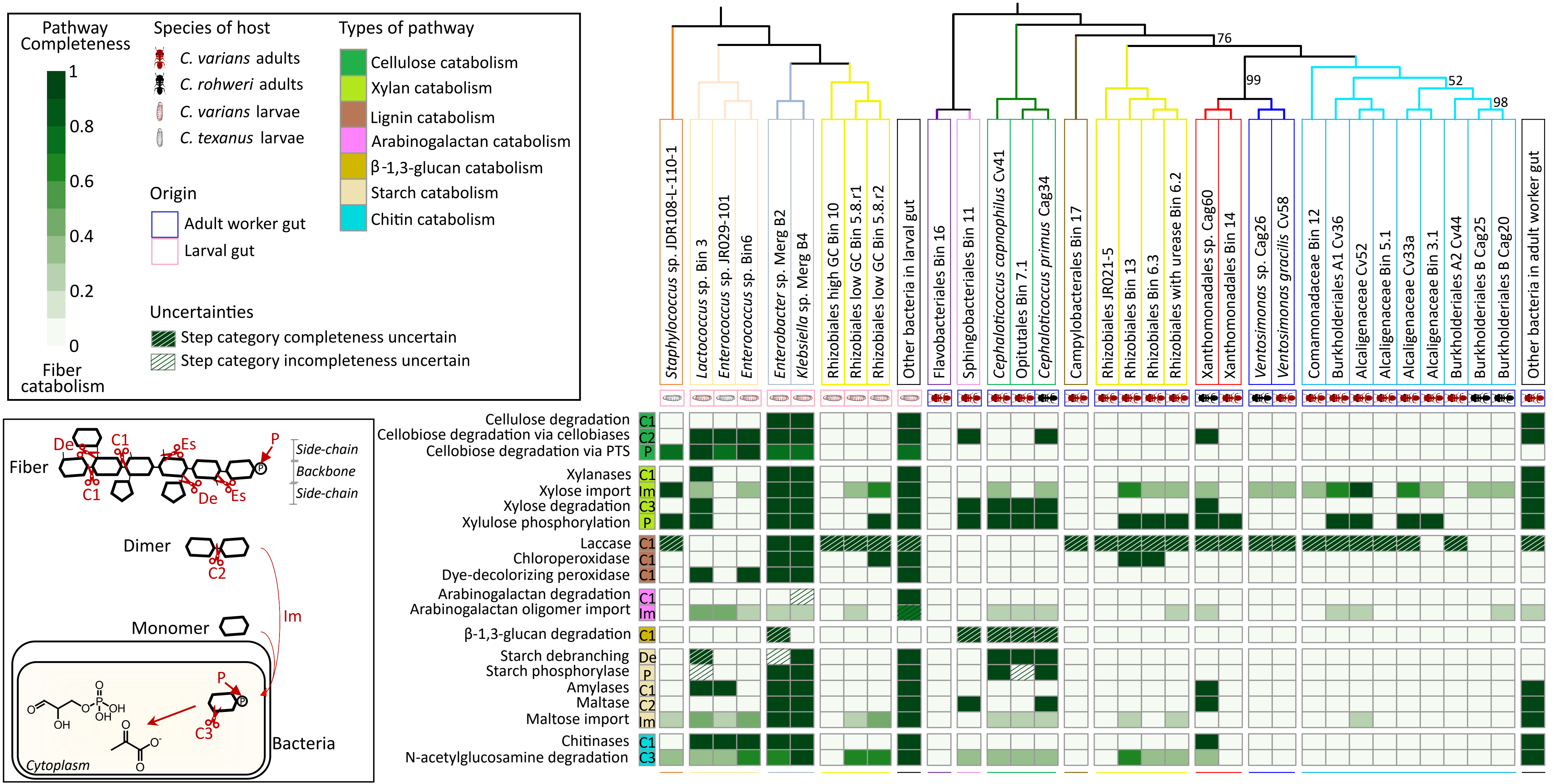

Pathway Completeness
1
0.8
0.6
0.4
0.2
0
Fiber catabolism
Species of host
C. varians adults
C. rohweri adults
C. varians larvae
C. texanus larvae
Origin
Adult worker gut
Larval gut
Types of pathway
Cellulose catabolism
Xylan catabolism
Lignin catabolism
Arabinogalactan catabolism
β-1,3-glucan catabolism
Starch catabolism
Chitin catabolism
Uncertainties
Step category completeness uncertain
Step category incompleteness uncertain
Fiber
De
C1
Es
P
Side-chain
Backbone
Side-chain
Dimer
C2
Im
Monomer
C3
Bacteria
Cytoplasm
76
99
52
98
Staphylococcus sp. JDR108-L-110-1
Lactococcus sp. Bin 3
Enterococcus sp. JR029-101
Enterococcus sp. Bin6
Enterobacter sp. Merg B2
Klebsiella sp. Merg B4
Rhizobiales high GC Bin 10
Rhizobiales low GC Bin 5.8.r1
Rhizobiales low GC Bin 5.8.r2
Other bacteria in larval gut
Flavobacteriales Bin 16
Sphingobacteriales Bin 11
Cephaloticoccus capnophilus Cv41
Opitutales Bin 7.1
Cephaloticoccus primus Cag34
Campylobacterales Bin 17
Rhizobiales JR021-5
Rhizobiales Bin 13
Rhizobiales Bin 6.3
Rhizobiales with urease Bin 6.2
Xanthomonadales sp. Cag60
Xanthomonadales Bin 14
Ventosimonas sp. Cag26
Ventosimonas gracilis Cv58
Comamonadaceae Bin 12
Burkholderiales A1 Cv36
Alcaligenaceae Cv52
Alcaligenaceae Bin 5.1
Alcaligenaceae Cv33a
Alcaligenaceae Bin 3.1
Burkholderiales A2 Cv44
Burkholderiales B Cag25
Burkholderiales B Cag20
Other bacteria in adult worker gut
Cellulose degradation C1
Cellobiose degradation via cellobiases C2
Cellobiose degradation via PTS P
Xylanases C1
Xylose import Im
Xylose degradation C3
Xylulose phosphorylation P
Laccase C1
Chloroperoxidase C1
Dye-decolorizing peroxidase C1
Arabinogalactan degradation C1
Arabinogalactan oligomer import Im
β-1,3-glucan degradation C1
Starch debranching De
Starch phosphorylase P
Amylases C1
Maltase C2
Maltose import Im
Chitinases C1
N-acetylglucosamine degradation C3

**S16 Fig. Carbohydrate catabolic profile of *Cephalotes*-associated symbionts in larval-associated symbionts.** Heatmap depicting carbohydrate catabolisms encoded in genomes of distinct *Cephalotes*-associated bacterial gut symbionts. Proportions of step categories encoded in symbiont genomes (columns) to perform various metabolisms (rows) is given by the shade within the squares of the heatmap, the darker the more complete the metabolic step is. Symbiont genomes were screened in IMG/M-ER for the presence of key enzyme- and transporter-encoding genes. Bacterial genes from the *C. varians* PL010 metagenomes that were not binned to MAGs (with the mention "Bin" or "Merg") but having a predicted relevant function are pooled in the "Other bacteria in larval gut" and "Other bacteria from adult worker gut" columns, respectively. Additional IGs from *C. varians*, *C. texanus* and *C. rohweri* were included. The bacterial phylogenetic trees were inferred based on seven bacterial marker gene sequences using a maximum likelihood (ML) method and 999 bootstrap iterations. Bootstrap values <100 are shown on the tree. Hatched squares signal when step categories were retained as complete or incomplete but with contradicting evidence. See **S15 Fig** for details on abbreviations of enzyme/transporter types.
