## Supplementary material for "In their sister’s footsteps: Taxonomic divergence obscures substantial functional overlap among the metabolically diverse symbiotic gut communities of adult and larval turtle ants": S17_Fig

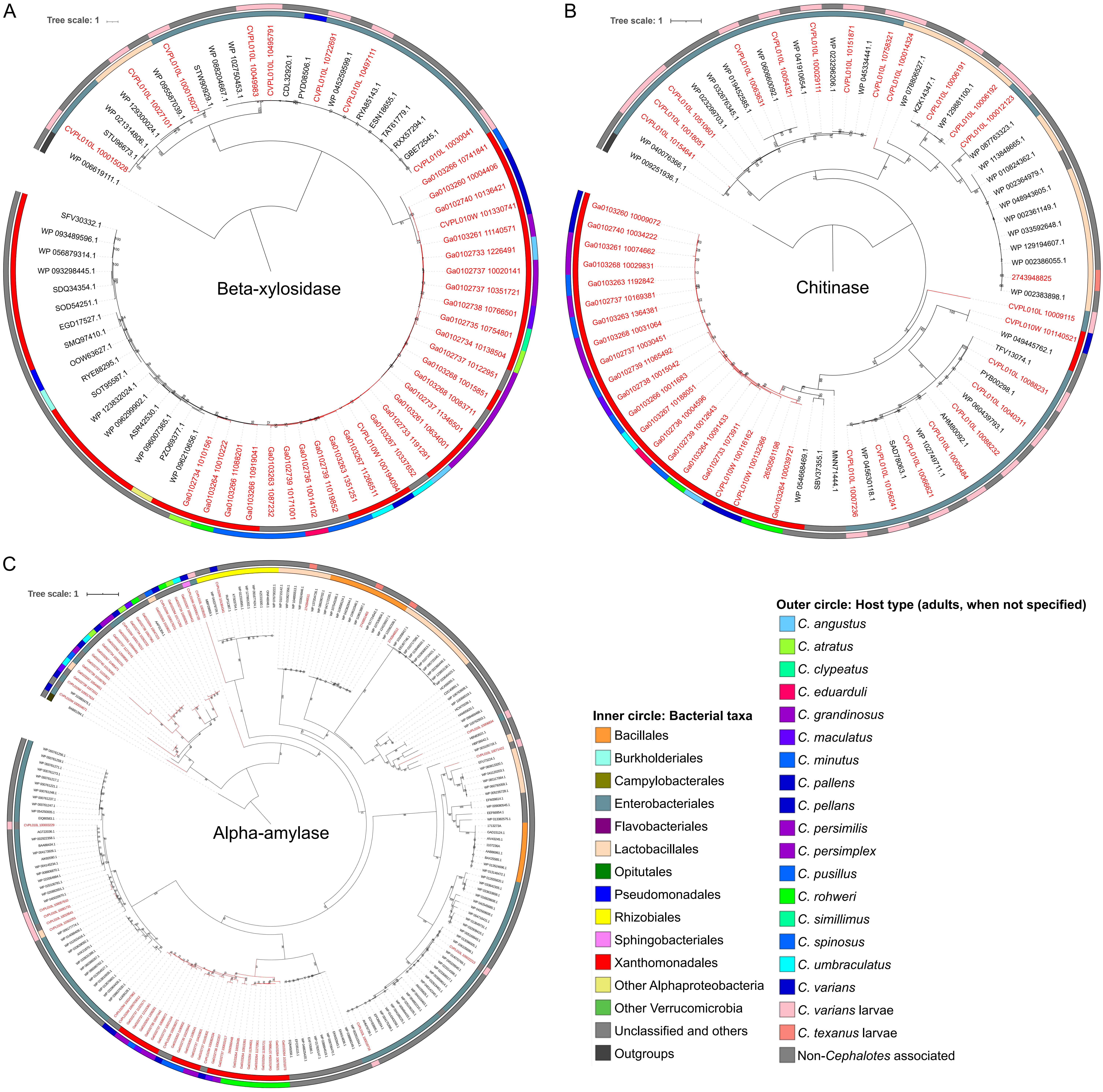

A
Tree scale: 1
Beta-xylosidase
CVPL010L 100015028
WP 006619111.1
STU96673.1
WP 021314806.1
WP 129300024.1
CVPL010L 10027101
WP 095587039.1
CVPL010L 100015027
STW90929.1
WP 088204667.1
WP 102750453.1
CVPL010L 100499883
CVPL010L 10495791
CDL32920.1
PYD08506.1
CVPL010L 10722691
WP 045259599.1
CVPL010L 10497111
RYA85143.1
ESN18655.1
TAT61779.1
RXX57294.1
GBE72545.1
CVPL010L 10030041
Ga0103266 10741841
Ga0103260 10004406
Ga0102740 10136421
CVPL010W 101330741
Ga0103261 11140571
Ga0102733 1226491
Ga0102737 10020141
Ga0102737 10351721
Ga0102738 10766501
Ga0102735 10754801
Ga0102734 10138504
Ga0102737 10122951
Ga0103268 10015851
Ga0103268 10083711
Ga0102737 11346501
Ga0103261 10634001
Ga0102733 1191291
Ga0103267 10337652
CVPL010W 100194094
Ga0103267 11266511
Ga0103263 1351251
Ga0102739 11019852
Ga0102736 10014102
Ga0102739 10171001
Ga0103263 1087232
Ga0103266 10919041
Ga0103268 11088201
Ga0103264 10010222
Ga0102734 10101561
WP 096210656.1
PZO69377.1
WP 096007365.1
ASR42530.1
WP 096299902.1
WP 123832024.1
SOT95587.1
RYE88295.1
OOW63627.1
SMQ97410.1
EGD17527.1
SOD54251.1
SDQ34354.1
WP 093298445.1
WP 056879314.1
WP 093489596.1
SFV30332.1
B
Tree scale: 1
Chitinase
WP 009251936.1
WP 040076366.1
CVPL010L 10154641
CVPL010L 10018051
CVPL010L 10510601
WP 023299703.1
WP 032676345.1
WP 019452585.1
CVPL010L 10063631
WP 060660092.1
CVPL010L 10054321
WP 041910654.1
CVPL010L 100029111
WP 023296206.1
CVPL010L 10151871
WP 045334441.1
CVPL010L 10758321
CVPL010L 100014324
WP 078806527.1
KZK14347.1
CVPL010L 10006191
WP 129881100.1
CVPL010L 10006192
CVPL010L 10012123
WP 087763323.1
WP 113848665.1
WP 010824362.1
WP 002364979.1
WP 048943605.1
WP 002361149.1
WP 033592648.1
WP 129194607.1
WP 002386055.1
2743948825
WP 002383898.1
CVPL010L 10009115
CVPL010W 101140521
WP 049445762.1
TFV13074.1
CVPL010L 10088231
PYB00298.1
CVPL010L 10040311
WP 060439793.1
CVPL010L 10088232
AHM80092.1
CVPL010L 10005484
WP 102749711.1
CVPL010L 10066621
SAD78063.1
CVPL010L 10156241
WP 045630118.1
CVPL010L 10007236
MNN71444.1
SBV37355.1
WP 054668469.1
Ga0103264 100039721
2650561198
CVPL010W 100132366
CVPL010W 100116162
Ga0102733 1073911
Ga0103264 10091433
Ga0102739 10012643
Ga0102736 10004596
Ga0103267 10188051
Ga0103266 10011683
Ga0102738 10015042
Ga0102739 11065492
Ga0102737 10030451
Ga0103268 10031064
Ga0103263 1364381
Ga0102737 10169381
Ga0103263 1192842
Ga0103268 10029831
Ga0103261 10074662
Ga0102740 10034222
Ga0103260 10009072
C
Tree scale: 1
Alpha-amylase
Inner circle: Bacterial taxa
Bacillales
Burkholderiales
Campylobacterales
Enterobacteriales
Flavobacteriales
Lactobacillales
Opitutales
Pseudomonadales
Rhizobiales
Sphingobacteriales
Xanthomonadales
Other Alphaproteobacteria
Other Verrucomicrobia
Unclassified and others
Outgroups
Outer circle: Host type (adults, when not specified)
C. angustus
C. atratus
C. clypeatus
C. eduarduli
C. grandinosus
C. maculatus
C. minutus
C. pallens
C. pellans
C. persimilis
C. persimplex
C. pusillus
C. rohweri
C. simillimus
C. spinosus
C. umbraculatus
C. varians
C. varians larvae
C. texanus larvae
Non-Cephalotes associated

**S17 Fig. Protein phylogenetic trees for conserved genes encoding fiber degrading-enzymes in the Xanthomonadales adult associates.** Rooted phylogenetic tree of **(A)** beta-xylosidase, **(B)** chitinase and **(C)** alpha-amylase amino acid sequences from *Cephalotes*-associated symbionts. Sequences were obtained from a screening in IMG/M-ER against seven *Cephalotes* metagenomes. These sequences were used as queries for an IMG/M-ER BLAST against the 20 *Cephalotes* metagenomes sequenced and the 14 *Cephalotes* IGs and for a BLASTP against custom NCBI databases. A maximum likelihood method was used with 999 bootstrap iterations. Bootstrap values are shown on the tree. Colors of host *Cephalotes* species (outer circle) are from [61].
