## Supplementary material for "In their sister’s footsteps: Taxonomic divergence obscures substantial functional overlap among the metabolically diverse symbiotic gut communities of adult and larval turtle ants": S18_Fig

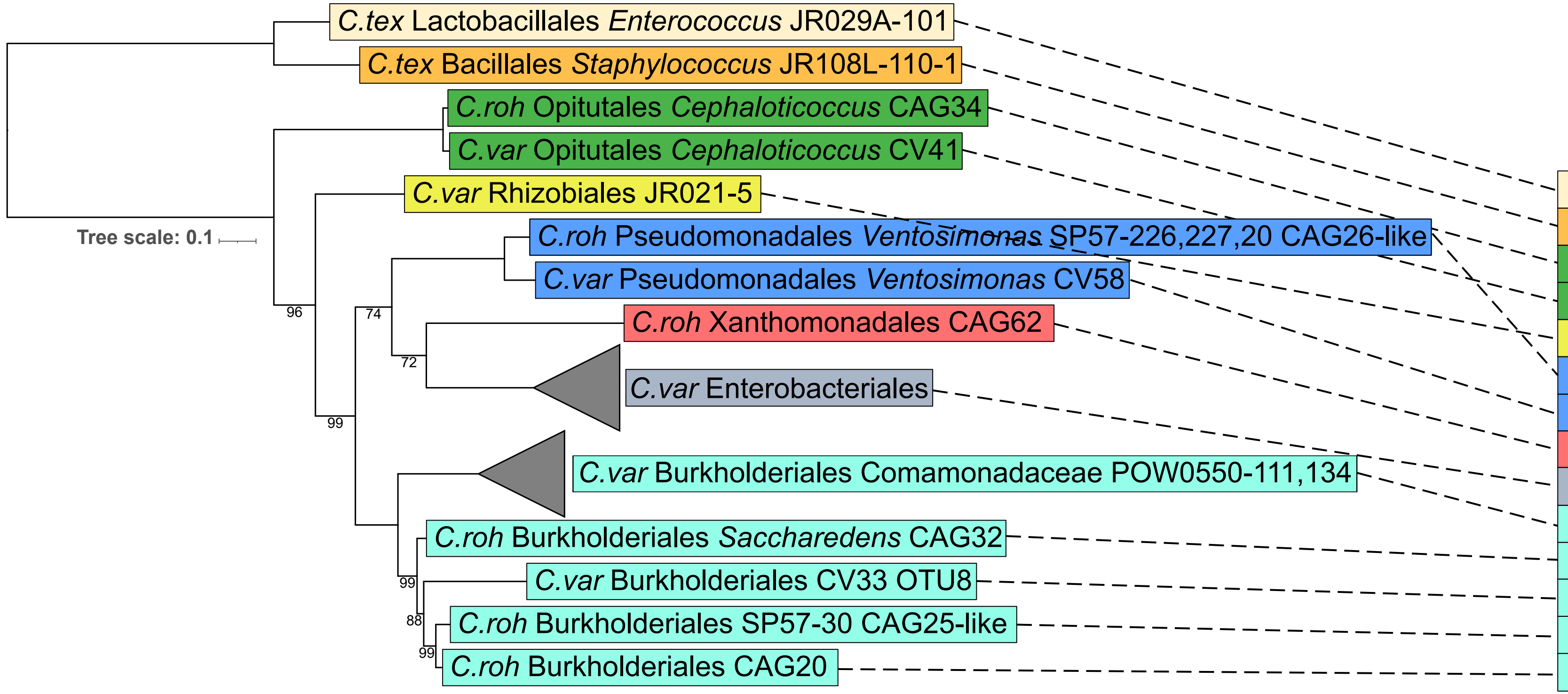


| Cultured symbiont strain | Host species and stage | D-Cellobiose (cellulose dimer) | N-Acetyl-D-Glucosamine (chitin monomer) | Amylose (starch backbone) | D-Maltose (starch dimer) |
|---|---|---|---|---|---|
| JR029A | | P | N | Ø | P |
| JR108L | | P | P | Ø | P |
| CAG34 | | P | N | Ø | P |
| CV41 | | P | N | Ø | P |
| JR021-5 | | N | N | Ø | P |
| CAG26-like | | N | N | Ø | N |
| CV58 | | N | N | Ø | N |
| CAG62 | | Ø | Ø | P | Ø |
| Ent. | | 15/16 | 16/16 | 2/8 | 15/16 |
| Com. | | Ø | Ø | 0/2 | Ø |
| CAG32 | | N | N | Ø | N |
| CV33 | | N | N | Ø | N |
| CAG25-like | | Ø | Ø | N | Ø |
| CAG20 | | N | N | Ø | N |

**S18 Fig. Results of *in vitro* metabolic assays for utilization of cellulose, chitin and starch derivatives.** The assays were conducted on cultured bacteria isolated from *Cephalotes varians* and *C. rohweri* adult guts and bacteria from *C. varians* and *C. texanus* larval guts. Each assay was run in triplicates. The phylogenetic tree was built from 16S rRNA sequences extracted from individual isolates using a ML method and 999 bootstrap iterations. Bootstrap values <100 are shown on the tree. Full results of *in vitro* metabolic assays can be found in **S8 Table**.
