## Supplementary material for "In their sister’s footsteps: Taxonomic divergence obscures substantial functional overlap among the metabolically diverse symbiotic gut communities of adult and larval turtle ants": S19_Fig

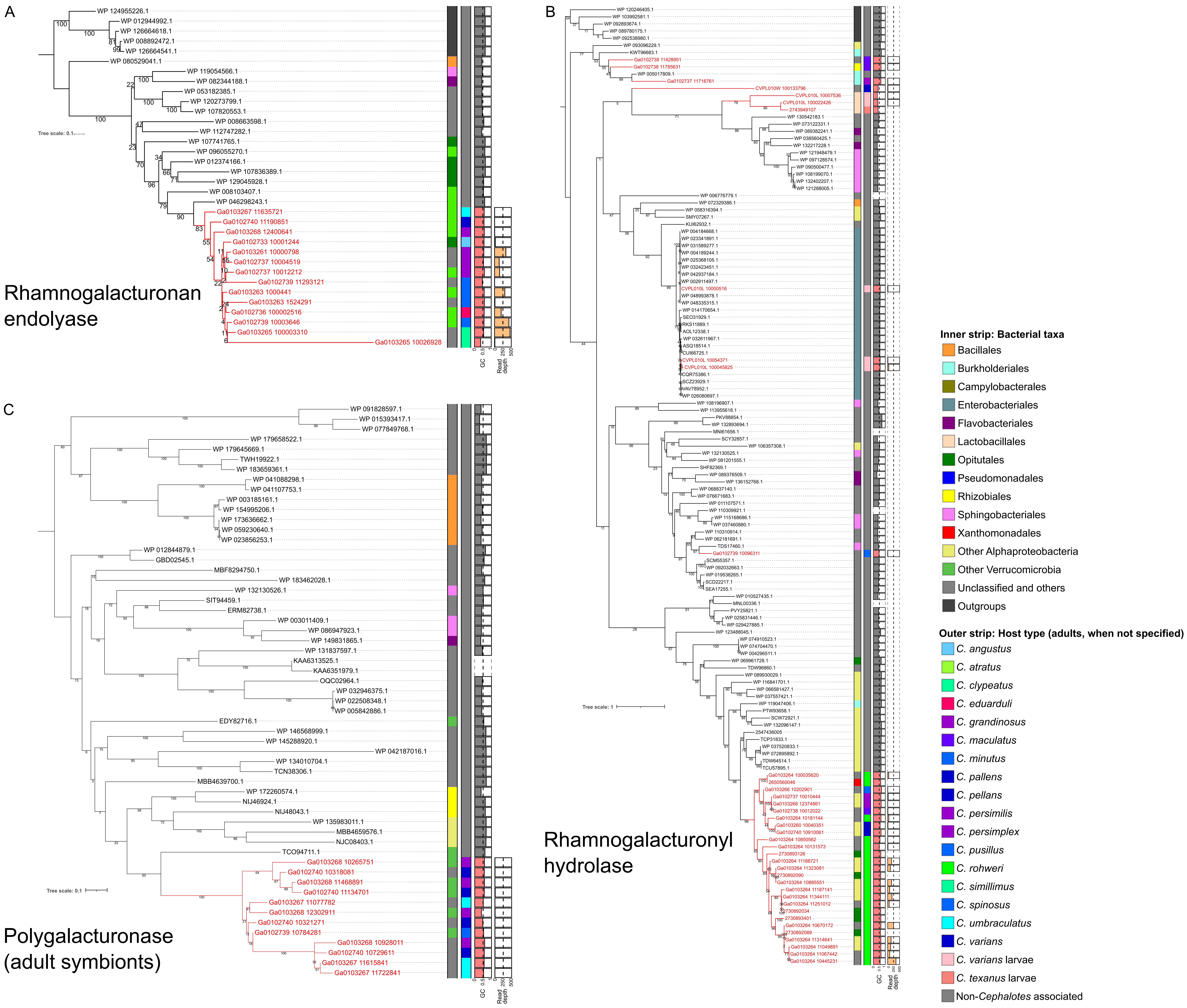

A
Rhamnogalacturonan endolyase
Tree scale: 0.1
GC
Read depth
B
Rhamnogalacturonyl hydrolase
Tree scale: 1
C
Polygalacturonase (adult symbionts)
Tree scale: 0.1
Inner strip: Bacterial taxa
Bacillales
Burkholderiales
Campylobacterales
Enterobacteriales
Flavobacteriales
Lactobacillales
Opitutales
Pseudomonadales
Rhizobiales
Sphingobacteriales
Xanthomonadales
Other Alphaproteobacteria
Other Verrucomicrobia
Unclassified and others
Outgroups
Outer strip: Host type (adults, when not specified)
C. angustus
C. atratus
C. clypeatus
C. eduarduli
C. grandinosus
C. maculatus
C. minutus
C. pallens
C. pellans
C. persimilis
C. persimplex
C. pusillus
C. rohweri
C. simillimus
C. spinosus
C. umbraculatus
C. varians
C. varians larvae
C. texanus larvae
Non-Cephalotes associated

**S19 Fig. Protein phylogenetic trees for pectinolytic enzymes in the *Cephaloticoccus* adult associates.** Rooted phylogenetic tree of **(A)** rhamnogalacturonan endolyase, **(B)** rhamnogalacturonyl hydrolase and **(C)** polygalacturonase (adults only) amino acid sequences from *Cephalotes*-associated symbionts. Trees were built similarly as in **S16 Fig**. Grey and coral horizontal bar graphs show the GC content of scaffolds from *Cephalotes*-associated symbionts (coral color) or of entire genomes from NCBI (grey color). Orange horizontal bar graphs show the read depths of scaffolds from *Cephalotes*-associated symbionts. Note that genes coding for these two enzymes from metagenomes of many *Cephalotes* species group into monophyletic branches with most scaffolds having similar GC content (0.56 on average) and occasionally with high read depth. Genes with such high read depth fit the characteristics of *Cephaloticoccus* sp.
