## Supplementary material for "In their sister’s footsteps: Taxonomic divergence obscures substantial functional overlap among the metabolically diverse symbiotic gut communities of adult and larval turtle ants": S20_Fig

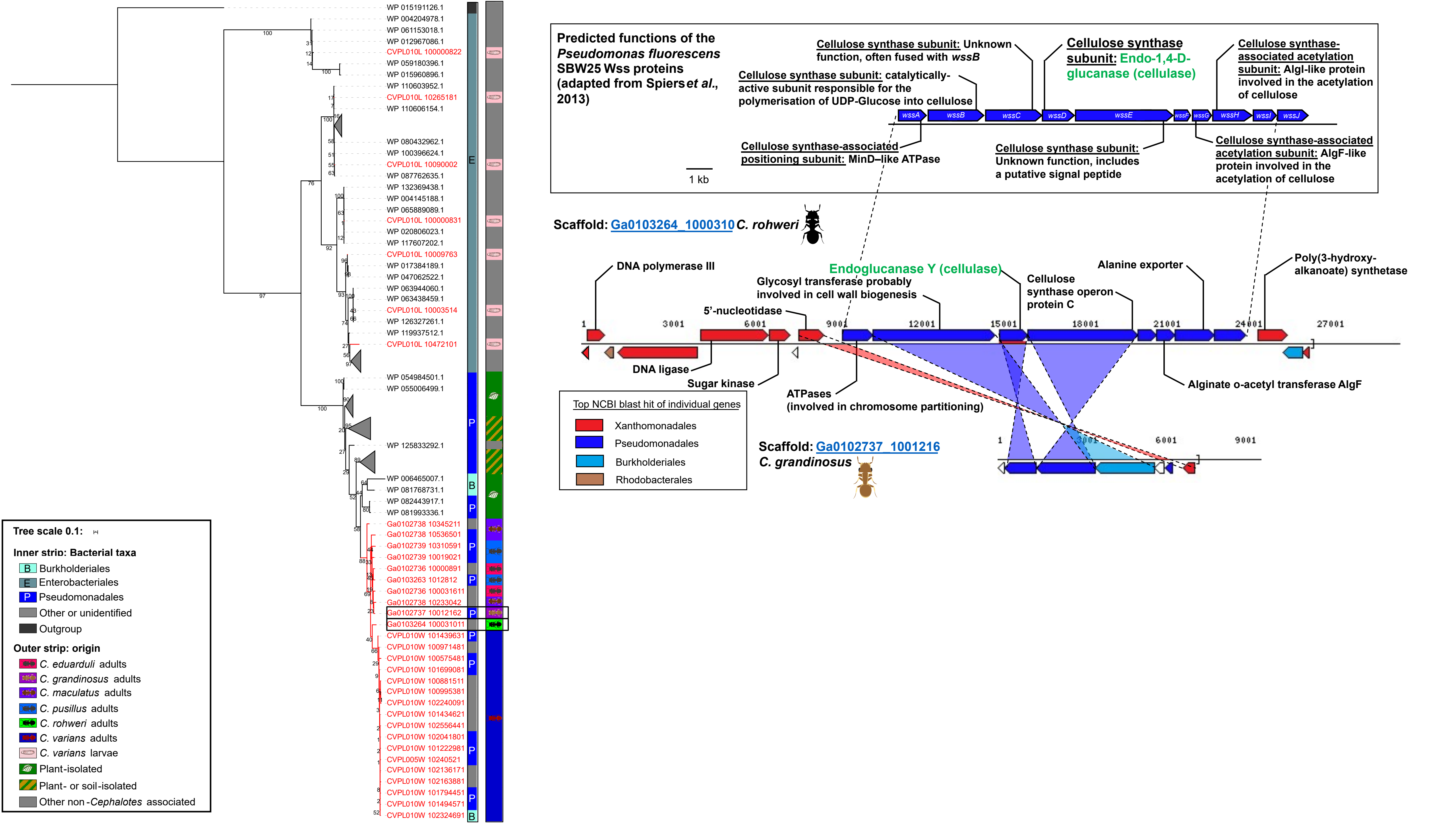

WP 015191126.1
WP 004204978.1
WP 061153018.1
WP 012967086.1
CVPL010L 100000822
WP 059180396.1
WP 015960896.1
WP 110603952.1
CVPL010L 10265181
WP 110606154.1
WP 080432962.1
WP 100396624.1
CVPL010L 10090002
WP 087762635.1
WP 132369438.1
WP 004145188.1
WP 065889089.1
CVPL010L 100000831
WP 020806023.1
WP 117607202.1
CVPL010L 10009763
WP 017384189.1
WP 047062522.1
WP 063944060.1
WP 063438459.1
CVPL010L 10003514
WP 126327261.1
WP 119937512.1
CVPL010L 10472101
WP 054984501.1
WP 055006499.1
WP 125833292.1
WP 006465007.1
WP 081768731.1
WP 082443917.1
WP 081993336.1
Ga0102738 10345211
Ga0102738 10536501
Ga0102739 10310591
Ga0102739 10019021
Ga0102736 10000891
Ga0103263 1012812
Ga0102736 100031611
Ga0102738 10233042
Ga0102737 10012162
Ga0103264 100031011
CVPL010W 101439631
CVPL010W 100971481
CVPL010W 100575481
CVPL010W 101699081
CVPL010W 100881511
CVPL010W 100995381
CVPL010W 102240091
CVPL010W 101434621
CVPL010W 102556441
CVPL010W 102041801
CVPL010W 101222981
CVPL005W 10240521
CVPL010W 102136171
CVPL010W 102163881
CVPL010W 101794451
CVPL010W 101494571
CVPL010W 102324691
E
P
B
Tree scale 0.1:
Inner strip: Bacterial taxa
B Burkholderiales
E Enterobacteriales
P Pseudomonadales
Other or unidentified
Outgroup
Outer strip: origin
C. eduarduli adults
C. grandinosus adults
C. maculatus adults
C. pusillus adults
C. rohweri adults
C. varians adults
C. varians larvae
Plant-isolated
Plant- or soil-isolated
Other non-Cephalotes associated
Predicted functions of the Pseudomonas fluorescens SBW25 Wss proteins (adapted from Spiers et al., 2013)
Cellulose synthase subunit: Unknown function, often fused with wssB
Cellulose synthase subunit: Endo-1,4-D-glucanase (cellulase)
Cellulose synthase-associated acetylation subunit: AlgI-like protein involved in the acetylation of cellulose
Cellulose synthase subunit: catalytically-active subunit responsible for the polymerisation of UDP-Glucose into cellulose
wssA
wssB
wssC
wssD
wssE
wssF
wssG
wssH
wssI
wssJ
Cellulose synthase-associated positioning subunit: MinD–like ATPase
Cellulose synthase subunit: Unknown function, includes a putative signal peptide
Cellulose synthase-associated acetylation subunit: AlgF-like protein involved in the acetylation of cellulose
1 kb
Scaffold: Ga0103264_1000310 C. rohweri
DNA polymerase III
Endoglucanase Y (cellulase)
Glycosyl transferase probably involved in cell wall biogenesis
Alanine exporter
Poly(3-hydroxy-alkanoate) synthetase
Cellulose synthase operon protein C
5’-nucleotidase
1
3001
6001
9001
12001
15001
18001
21001
24001
27001
DNA ligase
Sugar kinase
ATPases (involved in chromosome partitioning)
Alginate o-acetyl transferase AlgF
Top NCBI blast hit of individual genes
Xanthomonadales
Pseudomonadales
Burkholderiales
Rhodobacterales
Scaffold: Ga0102737_1001216 C. grandinosus
1
3001
6001
9001

**S20 Fig. Origins, and functional and taxonomic assignments of cellulase-encoding genes from *Cephalotes*-associated symbionts. (A)** Phylogenetic tree of cellulase amino acid sequences from *Cephalotes*-associated symbionts. The tree was built similarly as in **S16 Fig** and rooted using a *Gloeocapsa* sp. (Cyanobacteria) sequence (AFZ33259.1). Note that Hymenopteran, Opitutales and Rhizobiales sequences were removed from the tree due to a lack of support for their cellulase annotation. **(B)** Taxonomically annotated bacterial scaffolds with a cellulase-encoding gene in *Cephalotes* adult gut metagenomes. The two scaffolds at the bottom are the longest assembled scaffolds from *Cephalotes* gut metagenomes containing a cellulase-encoding gene. The colors of the genes correspond to their bacterial taxonomic annotation at the order level inferred with online NCBI BLASTNs.

Taxonomic annotation of single genes shows evidence of a cellulose-synthesis operon horizontally transferred from a Pseudomonadales bacterium to a Xanthomonadales *Cephalotes*-associated bacterial symbiont. The scaffold on the top shows the *wssA-J* cellulose-synthetase operon from the biofilm-producing *Pseudomonas fluorescens* SBW25 detailed in [151]. Note that a cellulase-encoding gene is naturally present on such cellulose-synthetase operon.
