## Supplementary material for "In their sister’s footsteps: Taxonomic divergence obscures substantial functional overlap among the metabolically diverse symbiotic gut communities of adult and larval turtle ants": S21_Fig

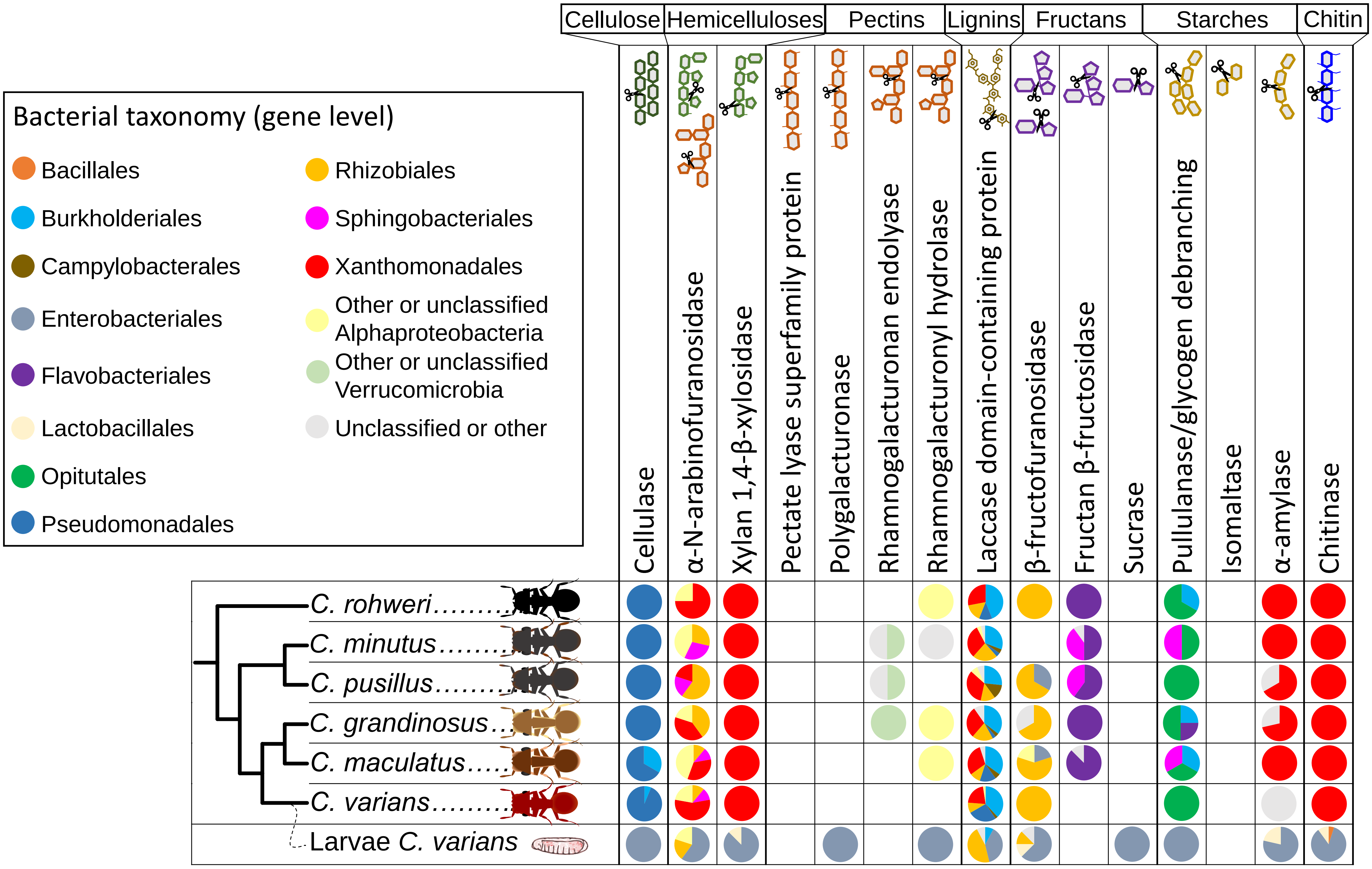

Cellulose
Hemicelluloses
Pectins
Lignins
Fructans
Starches
Chitin
Cellulase
α-N-arabinofuranosidase
Xylan 1,4-β-xylosidase
Pectate lyase superfamily protein
Polygalacturonase
Rhamnogalacturonan endolyase
Rhamnogalacturonyl hydrolase
Laccase domain-containing protein
β-fructofuranosidase
Fructan β-fructosidase
Sucrase
Pullulanase/glycogen debranching
Isomaltase
α-amylase
Chitinase
Bacterial taxonomy (gene level)
Bacillales
Burkholderiales
Campylobacterales
Enterobacteriales
Flavobacteriales
Lactobacillales
Opitutales
Pseudomonadales
Rhizobiales
Sphingobacteriales
Xanthomonadales
Other or unclassified Alphaproteobacteria
Other or unclassified Verrucomicrobia
Unclassified or other
C. rohweri
C. minutus
C. pusillus
C. grandinosus
C. maculatus
C. varians
Larvae C. varians

**S21 Fig. Gene-level taxonomy of symbiont genes for herbivore and fungivore digestive enzymes (PFCWDEs) across *Cephalotes* species.** This figure is similar to **Fig 8**, but the bacterial taxonomic classification was performed at the gene level here, instead of scaffold level. The pie charts show the proportion of genes from bacteria identified at the taxonomic order (colors) to code for specific enzymes (columns) in given *Cephalotes* hosts (rows). The *Cephalotes* phylogeny is adapted from [102]. Contrasting with **Fig 8**, the classification is less ambiguous, indicating potential horizontal gene transfers having occurred among *Cephalotes*-associated gut bacteria.
