## Supplementary material for "In their sister’s footsteps: Taxonomic divergence obscures substantial functional overlap among the metabolically diverse symbiotic gut communities of adult and larval turtle ants": S22_Fig

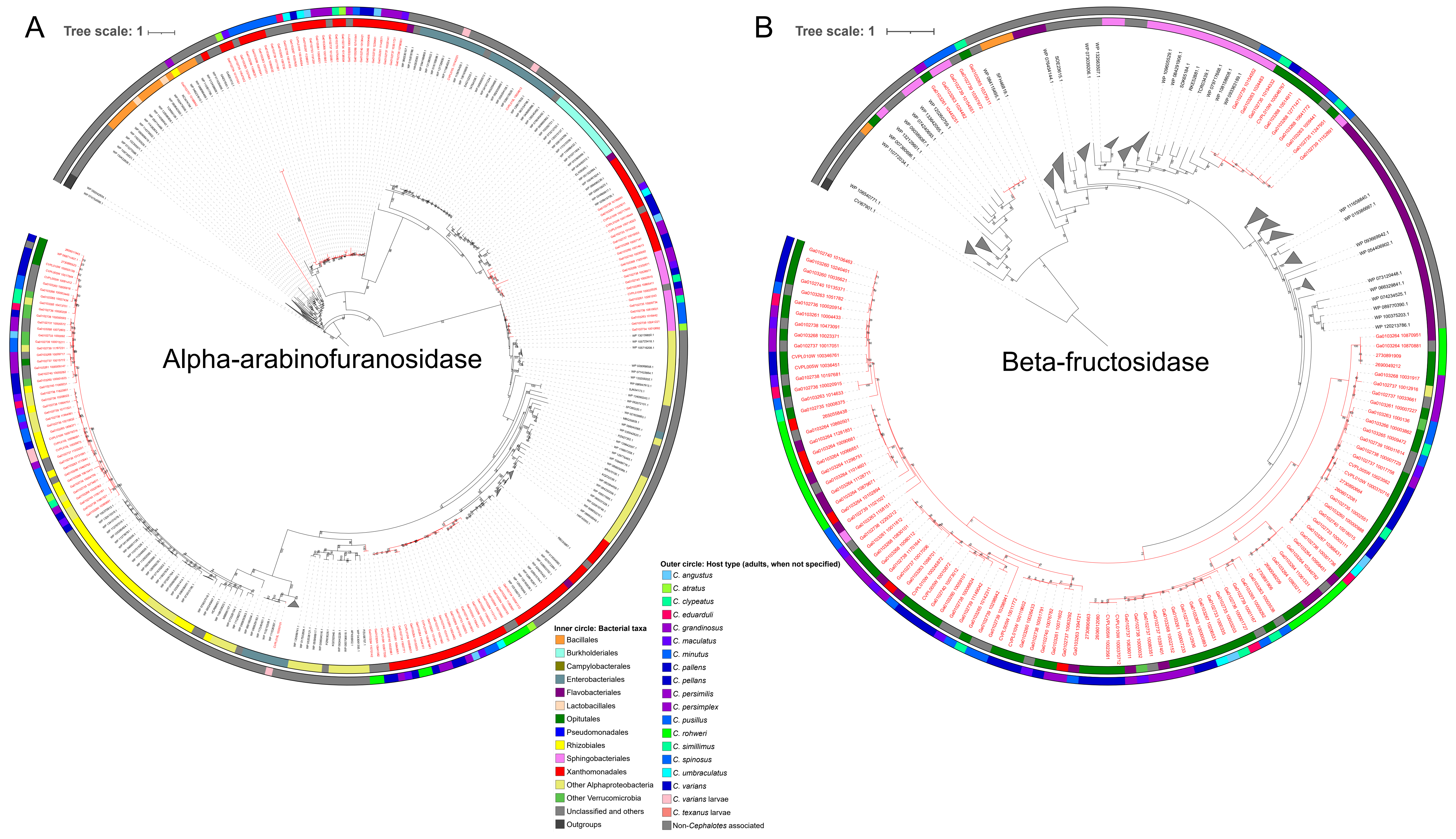

A
Tree scale: 1
Alpha-arabinofuranosidase
B
Tree scale: 1
Beta-fructosidase
Inner circle: Bacterial taxa
Bacillales
Burkholderiales
Campylobacterales
Enterobacteriales
Flavobacteriales
Lactobacillales
Opitutales
Pseudomonadales
Rhizobiales
Sphingobacteriales
Xanthomonadales
Other Alphaproteobacteria
Other Verrucomicrobia
Unclassified and others
Outgroups
Outer circle: Host type (adults, when not specified)
C. angustus
C. atratus
C. clypeatus
C. eduarduli
C. grandinosus
C. maculatus
C. minutus
C. pallens
C. pellans
C. persimilis
C. persimplex
C. pusillus
C. rohweri
C. simillimus
C. spinosus
C. umbraculatus
C. varians
C. varians larvae
C. texanus larvae
Non-Cephalotes associated

**S22 Fig. Protein phylogenetic trees for putatively horizontally exchanged genes encoding fiber-degrading enzymes in the *Cephaloticoccus* adult associates.** Rooted phylogenetic tree of **(A)** alpha-arabinofuranosidase and **(B)** beta-fructosidase amino acid sequences from *Cephalotes*-associated symbionts. Trees were built similarly as in **S16 Fig**. Based on these reconstructed phylogenies, *Cephaloticoccus* sp. may have received an alpha-arabinofuranosidase-encoding gene from Alphaproteobacteria, and beta-fructosidase-encoding genes may have been transferred from Flavobacteriales or Sphingobacteriales to Xanthomonadales and *Cephaloticoccus* sp.
