## Supplementary material for "In their sister’s footsteps: Taxonomic divergence obscures substantial functional overlap among the metabolically diverse symbiotic gut communities of adult and larval turtle ants": S1_Text

S1 Text – Supplementary materials and methods

Content

### 1 Cultured bacterial isolates

#### 1.1 Isolation, cultivation, and identification of bacteria

To obtain bacterial isolates, wild turtle ants were collected from a range of habitats in North, Central and South America, through excavations of hollowed tree branches. Dissected guts of adults and larvae with a visible dark food mass in their digestive systems were surface-sterilized in 70% isopropanol and homogenized in a glass tissue homogenizer in sterile trypticase soy broth (TSB; Difco BD) alone or supplemented with 250 μl Mallantoin (Sigma). Afterwards, the homogenate was serially diluted and spread onto trypticase soy agar (TSA; Difco BD) and TSA supplemented with 10% (v/v) sheep blood (Fisher Scientific). The plates were incubated in air, or in atmosphere-controlled glove boxes (Coy Labs) under hypoxic (2% O_2_, 5% CO_2_, and 93% N_2_) and anoxic (5% H_2_, 5% CO_2_, and 90% N_2_) conditions at room temperature for approximately one month. Resulting bacterial colonies were isolated, purified, and identified by 16S rRNA gene sequencing using the universal primers 63F (5′-CAG GCC TAA CAC ATG CAA GTC-3′) and 1389R (5′-ACG GGC GGT GTG TAC AAG-3′) [1]. Taxonomic classifications for these bacteria were achieved by submitting these sequences to the Classifier program in the Ribosomal Database Project website ([rdp.cme.msu.edu](http://rdp.cme.msu.edu/)) [2].

As a complement to our genomic efforts, we studied metabolic functions of cultivable bacteria from larvae and adults. Isolates utilized included those newly cultivated for the present study and several of those from Hu et al. 2018 [3]. The below-described assays were conducted in triplicate for each isolated bacterium tested.

#### 1.2 Pectin plate assay

To assess pectin depolymerization capabilities from the isolated bacteria, the pectin plate assay method from Engel et al. (2012) [4] was followed. Briefly, isolated bacteria were grown on TSA at 25°C for three days. Test media were prepared by combining 1.5% w/v poly-d-galacturonic acid methyl ester (from apple, Sigma), 0.5% w/v agar (Difco, BD) in 80 ml of Tris-HCl pH 8.6 and 70 ml of purified water. The solution was sterilized by autoclaving, and 1.5 ml of the solution was pipetted into each individual well of a 24-well plate (Corning). Once the agar had solidified, a small amount of bacteria was transferred from a single TSA plate onto the middle of a single well using a sterile loop. This was done until all bacteria had been placed in single wells. After two days growth at 25°C, each well was flooded with 0.75 ml of a 1% w/v solution of cetyltrimethylammonium bromide (CTAB, Sigma). Development of a clear zone around the bacterial colony was considered evidence of pectinase activity.

#### 1.3 Colorimetric assay

Carbohydrate and organic acid degradation (depolymerization or oxidation) was determined by using GN3 microplates (Biolog) according to manufacturer’s instructions. Bacteria were inoculated by taking a small sample from a TSA plate with a sterile inoculator swab (Biolog). The bacteria were re-suspended in inoculating fluid A (IF-A, Biolog) until the transmittance (measured at 600nm in a spectrophotometer) was 98%. A 150 µl aliquot of inoculating fluid was placed into each well on the plate.

### 2 Genomic and metagenomic analyses

#### 2.1 Functional dissimilarities in larval *vs.* adult symbionts

KO assignments were downloaded from IMG/M-ER [8] for *Cephalotes* varians PL010 larval and adult worker gut metagenomes and for all cultured isolate genomes. In Excel, for each of the 32 larval and adult symbiont genomes (Metagenome-Assembled Genomes (MAGs) and Cultured-Isolate Genomes (IGs)), gene counts per KO were calculated and normalized by the total number of KO-encoding genes. The resulting matrix was imported in R v3.6.3 [9]. Non-Metric Multi-Dimensional Scaling (NMDS) was performed using the *metaMDS* command from the ‘vegan’ R package [10], with the Bray-Curtis dissimilarity index, 2 dimensions, 500 maximum numbers of random starts in search of stable solution (a solution was reached at the 20^th^ run) and 9,999 maximum number of iterations in the single NMDS run. The goodness of fit to the two NMDS axes for each of the 32 symbionts was verified using the *goodness* and *stressplot* commands from the ‘vegan’ package. The graphical representation in **Fig 1.B** was drawn using the ‘ggplot2’ R package [11] and annotated using Inkscape 0.92 (<https://inkscape.org/release/inkscape-0.92/?latest=1>).

To test the difference in KO enrichment between larval and adult symbiont genomes a Permutational Multivariate Analysis of Variance (PERMANOVA) was performed using the *adonis* command from the ‘vegan’ package [10] with the Bray-Curtis distance and 9,999 permutations. The NMDS was performed a second time, using the same approach, after removing genomes with >95% OrthoANI (see **S8 Fig**). The NMDS graph was very similar to the original, helping to verify that pseudoreplication did not drive the observed patterns. R codes are available in the repository.

#### 2.2 Gene count per KEGG pathway in *Cephalotes varians* PL010 adult worker and larval gut metagenomes

To evaluate the relative weight of functional pathways in the *Cephalotes varians* PL010 adult worker and larval gut metagenomes, bacterial gene count per KEGG pathway was extracted from IMG/M-ER. The genes from single metagenomes were loaded in the Gene Cart. To extract only bacterial genes from IMG/M-ER, scaffolds from genes in the Gene Cart were loaded to the Scaffold Cart and a filter was applied to keep only bacterial scaffolds. Then genes from scaffolds in our filtered Scaffold Cart were *de novo* loaded in our empty Gene Cart

and saved as Gene Sets. For each Gene Set, the Gene Set Function Profile (KEGG_Pathway_KO) was generated and exported as an Excel file. Since the limit of genes in the Gene Cart is 20,000, we replicated this saving of Gene Sets and exporting of Gene Set Function Profile results multiple times for metagenomes, concatenating the output Excel files.

To normalize the bacterial gene counts for each KEGG pathway and facilitate the comparison between *Cephalotes varians* PL010 adult worker and larval gut metagenomes, we used *Z*-normalized log odds ratios (*Z*-LOR). To calculate the this statistic, we applied a modified formula available in the IMG/M-ER Function Comparisons tool: *Z*-LOR = log_10_{[(x_1_ / n_1_) / ((n_1_ - x_1_) / n_1_)] / [(x_2_ / n_2_) / ((n_2_ - x_2_) / n_2_)] / [ sqrt(1 / x_1_ + 1 / not_x_1_ + 1 / x_2_ + 1 / not_x_2_)]}, with x_1_ = gene count for the KEGG pathway in metagenome#1, x_2_ = gene count for the this pathway in metagenome#2, n_1_ = total count of genes possessing a KO assigned to this pathway from metagenome#1, n_2_ = total count of genes possessing a KO from the same pathway metagenome#2, not_x_1_ = n_1_ - x_1_ and not_x_2_ = n_2_ - x_2_. To avoid dividing by zeros, 0.0000001 was added to x_1_, x_2_, n_1_ and n_2_ before the calculations.

The significance of the normalized gene count per pathway between *Cephalotes varians* PL010 adult worker and larval gut metagenomes was evaluated by calculating the *p*-value as followed: *p*-value = 1 - pnorm(abs(*Z*-LOR)), with pnorm = probability distribution function for *N*(0, 1) and abs = absolute. The normalized gene count for a pathway was determined to be significantly different between the *Cephalotes varians* PL010 adult worker and larval gut metagenomes when the *p*-value was below the 0.05 threshold after applying the False Discovery Rate correction of Benjamini-Hochberg [12] using the *p.adjust* command in R [9]. The graphical representation in **Fig 1.C** was drawn using the ‘plotly’ package [13], with the R script being available in as a repository file.

#### 2.3 CAZy screening of *Cephalotes varians* PL010 adult worker and larval gut metagenomes

The *Cephalotes varians* PL010 adult worker and larval gut metagenomes were screened for carbohydrate-active enzymes (*i.e.* CAZy, <http://www.cazy.org/>: [14]) using the dbCAN2 metaserver (<http://bcb.unl.edu/dbCAN2/blast.php>: [15]). This metaserver combines three databases (HMMER, DIAMOND and Hotpep) to annotate genes from sequences uploaded as FASTA files.

Since the maximum file size that can be uploaded is 20 MB, the metagenome amino acid sequences in FASTA format that were downloaded from IMG/M-ER were split into smaller FASTA files in order to be uploaded in dbCAN2. The output files were concatenated into a single table per metagenome at the end of the screenings. For each metagenome, the output file contained three columns corresponding to each database (HMMER, DIAMOND and Hotpep). The annotations from HMMER were the first to be kept. Then, the genes not annotated in the HMMER database but annotated in the DIAMOND database were kept. Finally, the genes that were not annotated in the HMMER database, nor in the DIAMOND database, but in the Hotpep database, were kept.

Scaffold-level taxonomic annotations, read depths and gene counts from IMG/M-ER were then added to each gene with a CAZy annotation. The CAZy families found in the metagenomes were categorized by the type of substrate the enzymes can degrade based on the review of their CAZypedia description or CAZy activities in the CAZy website [14]. For each CAZy family, the sum of gene counts per bacterial order was calculated. The bar graphs in **S14 Fig** were drawn using the ‘ggplot2’ R package [11] (R script available in the repository database) and annotated using Inkscape 0.92 (<https://inkscape.org/release/inkscape-0.92/?latest=1>).

#### 2.4 Manual curation of key metabolic functions encoded within *Cephalotes*-associated cultured isolate genomes and *C. varians* colony PL010 metagenomes

Building upon the above analyses, we aimed to more carefully define the abilities of symbiotic bacteria to fulfill a focused set of metabolic functions of known [3] or likely importance in their symbioses with turtle ants. Below we detail the methodologies used to identify the involved genes and the strength of evidence supporting their functional annotations, using a protocol that is outlined in **S4 Fig**.

##### 2.4.1 Focal metabolic functions – identification and review

We hypothesized a role for *Cephalotes* gut symbionts in a range of digestive and nutritive functions, exploring the distributions of these functions via genomic inferences, across adults and larvae. Based on known contributions of symbiotic microbes to ants [3] or other insects [*e.g.* 20–24], we focused on symbiont catabolism of plant fibers, N-recycling, organic acid and sulfur metabolism, and both B-vitamin and amino acid biosynthesis.

To explore the abilities of *Cephalotes*-associated bacteria to catabolize polymers from the ant host diet – a fairly broad set of functions – we first investigated the potential food sources of the ant hosts by reviewing the literature. We focused on records of natural elements being collected by *Cephalotes* ants. The main fibers constituting these potential food sources were further considered as substrates of interest. We chose to focus on complex carbohydrates (pectins, cellulose, hemicelluloses, arabinogalactan, beta-1,3-glucans, starches and chitin) and lignins because these fibers are usually abundant staples in a generalist herbivorous diet. For fibers known to have variable structures, we chose to focus our primary analyses on the simplest types of structures, such as homogalacturonan for pectins [21] and xylan for hemicelluloses [22].

To identify the specific pathways and the component enzymatic/transport ‘steps’ for the aforementioned symbiotic functions, we used the KEGG database (<https://www.genome.jp/kegg/kegg2.html> [23]). The KEGG website provides regularly updated maps of metabolic pathways, giving insights into the known biochemical mechanisms used in the catabolism and anabolism from a wide range of organisms. IMG/M-ER includes data from 399 KEGG pathways, each including numerous biochemical networks, and transport mechanisms, for the transformation and movement of substrates inside and outside of cells. Identifying the KEGG pathways, the pertinent reaction chains contained within, and the genes of relevance for our focal functions was assisted by literature review.

To identify useful search terms in our below-performed Function Searches we also utilized individual web pages devoted to substrates, enzymes, KEGG KOs, and full pathways, obtaining additional information that is sometimes missing from KEGG pathway maps for eventual use in our screening (below). In particular, the “Reaction(IUBMB)” and “Comments” sections on the “ENZYME” pages were especially useful, listing alternative enzymes for a given step of a metabolic pathway, the levels of enzyme specificity, and the organisms synthesizing each enzyme.

##### 2.4.2 Genomic and metagenomic metabolic screening in IMG/M-ER

Next, to identify genes/proteins of relevance in our examined symbiotic functions (*i.e.* those encoding ‘steps’ of the focal metabolic pathways), we separately added the full set of 14 Cultured-Isolate Genomes (IGs), the *Cephalotes* *varians* PL010 adult worker metagenome, or the PL010 larval gut metagenome to the IMG/M-ER Genome Cart. We then performed an IMG/M-ER Function Search using Enzyme Commission (EC) numbers (for all focal pathways except amino acid biosynthesis) and KEGG Orthology (KO) numbers (for all focal pathways) as keywords. The selected ECs and KOs corresponded to the genes/proteins fulfilling each enzymatic transformation or cross-membrane transport event (*i.e.* ‘step’) in our selected focal pathways.

For several Function Searches, we used additional keywords corresponding to the full names of enzymes or the abbreviations of genes encoding membrane transporters, as retrieved from individual KEGG ENZYME webpages (<https://www.genome.jp/kegg/annotation/enzyme.html> [24]). This approach was applied to enzymes encoding metabolic transformations demonstrated *in vitro* (*e.g.* homogalacturonan catabolism, N-recycling) that were initially absent after our KOs and EC screening. For instance, given the lack of evidence for pectinase KOs and ECs, despite positive *in vitro* pectinase assays on a number of isolates (**Fig. 6.C**), we further searched for pectin methylesterase enzymes using the keywords “pectin methylesterase”, “pectin methyl esterase”, “pectinesterase”, “pectin demethoxylase”, “pectase” and “pectinoesterase”.

Results of Function Searches were added to the IMG/M-ER Gene Cart. These data were downloaded as an Excel table, with each row corresponding to one gene from a genome or metagenome that was annotated to encode a step of a focal metabolic pathway. Columns of these excel files contained information such as gene and scaffold identifiers, lengths, scaffold read depth, GC% and gene functional annotation from Pfam, COG, KEGG databases, and EC number. As we describe below, results of several annotation methods were then used to more confidently predict gene presence/absence, aiding in our metabolic reconstruction.

##### 2.4.3 Filtering our screening results

We, next, applied priority criteria to filter ambiguous annotations, helping to refine the aforementioned Function Search results. Our goal was to target enzymes that are active on a narrow spectrum of substrates, thereby improving confidence in our metabolic reconstructions. For this purpose, we prioritized the KEGG KO and EC number annotations. These classify enzymes into more accurate categories than Pfam, COG, or CAZy annotations, which can group proteins with different substrates into broad families.

We, therefore, took the presence of genes/proteins assigning to an EC or KO number from a focal pathway to indicate step fulfillment in the given genome/metagenome; and all such genes were retained. For each gene/protein without an EC or KO assignment, and for those linked to an EC or KO from outside the given focal pathway, we examined COG category assignments, in an effort to link more genes to focal pathway steps. When no EC number, KEGG KO, or COG ID annotation linked the gene to the step/function of interest, the Pfam annotation was examined for each remaining gene. All genes lacking assignments to the expected pathway steps based on these four aforementioned databases, were removed from our tables and from further analyses. The retained genes were given a step number corresponding to their order within their focal metabolic pathway.

##### 2.4.4 Taxonomic information

For metagenome datasets, taxonomic information for each scaffold containing a retained, focal pathway step-encoding gene was combined with the information downloaded from after IMG/M-ER Function Searches and filtering. To achieve this, we first downloaded the IMG/M-ER metagenome data by loading a given metagenome in the IMG Genome Cart and selecting “Download Genomes”. From this folder, we opened the “phylodist” file in Excel (accessible as a data repository), separating the classification data into multiple columns. Using a “VLOOKUP()” function, we populated a new column with order-level classifications of the scaffolds. Additionally, we added columns showing the binning/assignment of some scaffolds to Anvio’ CONCOCT Metagenome-Assembled Genomes (MAG), by using the “VLOOKUP()” function from the Anvio’ CONCOCT binning Excel file (**S5 Table**).

##### 2.4.5 Further annotation verification for carbohydrate catabolism and N-recycling pathways

To further verify the gene functional annotations of carbohydrate catabolism and N-recycling pathways – two functions of special interest in our study – a double BLASTP against the SWISS-PROT and NCBI non-redundant (nr) databases was performed using the FASTA amino acid sequences of our filtered Function Search results. The top ten hits per gene with a significant match (Hsp Expected <0.05) were retained (raw BLAST results are available as repositories). For each gene, the functional annotation of the top hit in the two databases was added in two new columns in the Excel file generated as described above. Since the SWISS-PROT database is rigorously and regularly curated [25], the BLAST results from this database that matched our expectations based on our filtered/curated annotations were retained. For example, when there were no hits with the NCBI nr database but hits with the SWISS-PROT database or when there were conflicts between the top hits of these two databases, we chose to keep only the hits with the SWISS-PROT database. When there were no hits from the SWISS-PROT database, we retained the hits from the NCBI nr database. When there were no adequate hits from both databases, genes were kept in the files, but considered as misannotated and not included in final figures.

##### 2.4.6 Post-processing and visualization

From these verified screening results, we computed completeness for our focal pathways, doing so for each cultured IG, each MAG, and then for the ‘left-over’ scaffolds, not assigned to MAGs, from each of our two *C. varians* metagenomes.

For carbohydrate degradation pathways, steps were classified in step categories based on the type of action of the putative enzymes. Step categories included esterases (“Es”); debranching enzymes (“De”) that remove side-chains from the backbone of a targeted fiber; phosphorylases (“P”); fiber backbone-cutting enzymes (“C1”); dimer-cutting enzymes (“C2”); and monomer-catabolizing enzymes (“C3”); and molecule import (“Im”). The pathway step category completeness was calculated as the proportion of steps of metabolic pathways in single step categories encoded by genes of MAGs and IGs. Some step categories only have one step (e.g. “Pectin esterase” or “Laccase”), thus their proportion was either 0 or 1. The heatmap was drawn using the ‘pheatmap’ R package [26] (R scripts and input tables are available in the repository database) and both the heatmap and N-recycling pathway map were edited in Inkscape 0.92 (<https://inkscape.org/release/inkscape-0.92/?latest=1>).

#### 2.5 Focus on key plant and fungal cell wall digestive enzymes (C1, De and Es) conserved across ant species

##### 2.5.1 Preparation of a list of digestive enzymes

To determine whether the initial steps of fiber degradation are encoded by bacterial gut symbionts hosted by adult workers of other *Cephalotes* species throughout the *Cephalotes* phylogeny, we first generated a list of 48 key polymer-degrading enzymes (**S4 Table**) based on the screening in *Cephalotes varians* adult worker and larval gut metagenomes described in the previous section, keeping only the enzymes involved in the first few steps of fiber carbohydrate catabolism. To this list, we added enzymes screened in other studies investigating the role of social insect mutualists [4,27] and we kept enzymes degrading substrates known to be consumed by *Cephalotes* ants [28–31]. For example, enzymes involved in the transformation of ceramide and kojibiose were removed from the list since we do not expect these molecules to be major constituents of the *Cephalotes* diet. Therefore, in addition to the enzymes initiating the degradation of fibers mentioned in the previous section (homogalacturonan for pectins, cellulose, xylan for hemicelluloses, arabinogalactan, beta-1,3-glucans, starches chitin, and lignin), we added enzymes acting on other types of pectin (*e.g.* rhamnogalacturonan-I) and hemicellulose (*e.g.* xyloglucan, glucuronoarabinoxylan, galactomannan) and on arabinosides, fructosides, fucosides and sucrose.

To avoid having identifiers from different databases (KO, EC, COG, Pfam) for a single enzyme, we selected only an identifier from one database for each enzyme, chosen with the same criteria as in *2.4.3*, with our KEGG prioritization. Thus, KEGG KOs and EC numbers were preferentially chosen. Some exceptions included enzymes that were not clearly annotated in IMG/M-ER with the KEGG KOs and EC numbers (*e.g.* pectate lyase) for which Pfam or COG IDs were kept.

##### 2.5.2 Metagenomic screening

The list of enzymes identified by EC numbers, KEGG KOs, COGs and Pfams was imported into IMG/M-ER as a new Function Set. A Function Profile was performed on five adult worker gut metagenomes from *Cephalotes* species spanning a good portion of the *Cephalotes* phylogeny (*Cephalotes rohweri*, *Cephalotes grandinosus*, *Cephalotes maculatus*, *Cephalotes pusillus* and *Cephalotes minutus*), in addition to the adult and larval *Cephalotes varians* PL010 metagenomes and the 14 isolated *Cephalotes* bacterial genomes. A Function Profile gives the gene count per function for each genome and metagenome. The genes obtained from the IMG/M-ER Function Profile were loaded in the IMG Gene Cart and exported as gene data in tab-delimited format.

Taxonomic assignments were given to each gene based on based on scaffold annotation from IMG/M-ER as described in *2.4.4.* In some cases, no genes were found encoding an enzyme strongly expected to be part of the given (meta)genome based on previous screenings (see *2.4*). In these cases, the list of enzymes was revised once using the Function Profile output to identify aberrant results. For example, for laccase the EC number EC:1.10.3.2 was never found in the metagenomes. The laccase EC was then replaced by the KEGG K05810, supposedly less specific because encoding various polyphenol oxidases. Although we tried to be as specific as possible by keeping in priority EC numbers and KEGG KOs, some enzymes are known for having a broad spectrum of potential substrates (*e.g.* beta-glucosidase EC:3.2.1.21 is active on beta-D-glucosides, beta-D-galactosides, alpha-L-arabinosides, beta-D-xylosides or beta-D-fucosides: <https://www.genome.jp/dbget-bin/www_bget?enzyme+3.2.1.21>).

##### 2.5.3 Exploring enzyme relatedness using protein phylogenetic trees

To identify relatedness of candidate fiber-degrading genes to others from symbionts of *Cephalotes* ants and those described previously in the literature, we constructed a series of phylogenetic trees. These were used, in several cases, to revise annotations, as described below. All the protein phylogenetic trees can be seen in the data repository.

Amino acid sequences for these genes were retrieved from IMG/M-ER and used as queries in two different BLAST searches. First, the queries were used in BLASTp searches against NCBI Identical Protein Groups databases. To obtain each Identical Protein Groups database, the generic name of the focal enzyme was used as a keyword to search against all NCBI databases (<https://www.ncbi.nlm.nih.gov/>). Then, all representatives from the corresponding Identical Protein Group(s) were downloaded as a FASTA file. Local BLASTP searches were performed with our *Cephalotes* symbiont-derived enzymes against their corresponding databases using the standalone BLAST software (<ftp://ftp.ncbi.nlm.nih.gov/blast/executables/LATEST/>). The top ten hits for each query were obtained in an output file and the sequences with an E-value below 0.05 were retained for the alignment.

The second BLASTp was performed in IMG/M-ER. All *Cephalotes* adult worker gut metagenomes and IGs in IMG/M-ER from a prior study [3], in addition to *Cephalotes varians* PL010 larval gut metagenome and the *Staphylococcus* sp. JDR108L-110-1 and *Enterococcus* sp. JR029-101 IGs new from this study, were loaded in the Genome Cart to serve as the database for this BLAST. We then used the same query genes as for our aforementioned local BLASTp. Hits with a percentage of similarity higher than 70% were kept for the alignment.

This value was decreased for alpha-amylase, since seemingly relevant hits had lower percentages of similarity in the blast results.

Sequences of all queries and top hits from the two BLASTs were aligned using the ClustalW multiple alignment algorithm in SeaView [32]. Alignments were uploaded into the CIPRES Science Gateway [33] to generate a maximum likelihood/rapid bootstrapping phylogeny using RAxML Black Box [34] with 999 bootstraps. The resulting tree with branch labels was uploaded to iTOL [35] for rooting and graphical annotation.

From these trees, genes in clades with a long branch that clearly dissociated from the other clades were used as queries for online NCBI BLASTp searches against the nr database. When all sequences from such dissociating clades yielded to hits with unexpected annotations (*i.e.* not the focal function), sequences from this clade were removed from the tree and considered as functionally misannotated.
