## Supplementary material for "In their sister’s footsteps: Taxonomic divergence obscures substantial functional overlap among the metabolically diverse symbiotic gut communities of adult and larval turtle ants": S2_Text

S2 Text – Supplementary results and discussion

Content

1. **Turtle ant gut bacterial communities**

Hu et al. [1] characterized dominant taxa across larvae of varying size for 12 *Cephalotes* species through 16S rRNA amplicon sequencing. Our shotgun-based data suggested a high abundance of Rhizobiales for both PL005 (**S2 Fig**) and PL010 (**Fig 1A**) colony larvae – matching overall trends of the amplicon-based study. But the colony PL005 differed in exhibiting a high abundance of Burkholderiales symbionts, a modest abundance of Pseudomonadales and Xanthomonadales symbionts, and low abundance of Enterobacteriales and Lactobacillales. While Hu et al. [1] showed that the latter two symbionts are not abundant in all larvae (**S3 Fig**) – and are, in fact, rare in those at younger stages – the average relative abundance of all Burkholderiales-assigned 97%-OTUs in larvae, across 12 *Cephalotes* species, was 0.046. Only one such Burkholderiales OTU ever exceeded an average relative abundance of 0.1 in larvae. Pseudomonadales were found in larvae of only 7 out of 12 surveyed *Cephalotes* species, showing an average relative abundance of 0.01, when present. Only one of the four *Cephalotes varians* colonies included in that study harbored this latter bacterial order in larvae, while Burkholderiales comprised an average of only 0.008 of all sequence reads from larvae in these same *Cephalotes varians* colonies.

In contrast, the amplicon-based study revealed that Lactobacillales and Enterobacteriales were abundant in larvae, accounting for average relative abundances of 0.22 and 0.11, respectively (**S3 Fig**). Larval gut communities enriched with these two bacterial orders, like our PL010 larval gut metagenome (**Fig 1A**)**,** are typical of older larval gut microbiomes [1]. For these reasons, we placed most of our focus on the PL010 larval metagenome to infer functions most likely representative of late larval stages, when larvae possess large masses of solid food in their gut.

2. **Patterns of horizontal gene transfer in the *Cephalotes*-symbiont system**

Horizontal gene transfer among *Cephalotes* symbiont lineages was suggested by differences in scaffold- *vs.* gene-level IMG/M-ER taxonomic annotations (**Fig 8; S21 Fig**) and by clustering patterns on protein phylogenetic trees (**S22 Fig)**. Illustrating this, verrucomicrobial (including *Cephaloticoccus* sp.) arabinofuranosidase-encoding genes seemed to be of alphaproteobacterial origin, based on gene-level taxonomy (**S22 Fig**), as were the rhamnogalacturonyl hydrolase-encoding genes encoded by this same abundant core symbiont group (**S19 Fig**). Similarly, genes coding for fructan beta-fructosidase may have been transferred on several occasions from Sphingobacteriales or Flavobacteriales to *Cephaloticoccus* sp. and relatives in the Opitutales, and to unnamed core symbiont species in the Xanthomonadales (**S22 Fig**).

3. **CO_2_ fixation by the Alcaligenaceae Cv33a/Bin3.1**

It is unclear how energy derived from sulfide oxidation in the Alcaligenaceae Cv33a/Bin3.1 would be utilized, since this symbiont is not capable of completing nitrate reduction or the rTCA cycle for CO_2_ fixation. This Burkholderiales symbiont might use other pathways coupled with sulfide oxidation for CO_2_ fixation or may be heterotrophic for its carbon sources [2]. As in other Betaproteobacteria, Alcaligenaceae Cv33a/Bin3.1 could, for instance, potentially perform anaerobic CO_2_ reduction using formate dehydrogenase [3]. Its presence in anoxic gut microhabitats – adult or larva – has not been confirmed, revealing more need for further investigation.

4. **Hypotheses on partner choice mechanisms in larval guts**

In the present study, we found that, compared to adults, and beside carbohydrate digestive capabilities, the gut microbiome from late-stage turtle ant larvae was enriched for genes encoding transporters and defensive functions (**Fig 1C**). This investment in extracellular interactions could be characteristic of bacterial symbionts having to survive in a highly competitive gut environment, where new microbial comers may regularly challenge the establishment of the larval gut bacterial community. But some mechanisms for the retention of specific larval gut symbionts may alternatively be contributed by adult-derived symbionts.

As documented in honeybees, iron is vital for bacterial symbiont gut colonization [4]. A recent investigation of biosynthetic gene clusters encoded by adult *Cephalotes*-associated symbionts showed that members of the Burkholderiales have retained their genes coding for iron-sequestering siderophores [5], that could putatively be secreted in adult gut lumens and transferred to larval guts. In our investigations on larval gut microbiomes, we found that the most encoded set of genes was the *fhu* genes, involved in the transport of iron-complexes, such as siderophores. Thus, although this remains speculative, it is possible that the aforementioned conserved Rhizobiales and environmentally-acquired Enterobacteriales and Lactobacillales can establish in larval guts thanks to their ability to evade competitive exclusion, harvest Burkholderiales-derived siderophores and provide digestive and nutritional benefits to their larval host.
